# Non-covalent reversibly photoconvertible fluorescent tags for wash-free protein labeling

**DOI:** 10.64898/2026.05.08.723694

**Authors:** Mrinal Mandal, Yuriy Shpinov, Alienor Lahlou, Fivos Pham, Lina El Hajji, Ian Coghill, Eve Laureau, Marie-Aude Plamont, Franck Perez, Thomas Le Saux, Isabelle Aujard, Arnaud Gautier, Ludovic Jullien

## Abstract

Reversibly photoswitchable fluorophores are widely used in advanced bioimaging but their design remains demanding. Here, we introduce a new series spanning the whole visible range, which results from combining a large set of fluorogens with the FAST protein scaffold. We first demonstrate that these well-established labeling fluorescent protein tags turn into negative reversible photoswitchers upon decreasing the fluorogen concentration and increasing light intensity. We then show that using not anymore one but two fluorogens adds new responses to illumination. Thus, we obtain positive reversible photoswitchers, that increase their brightness under illumination. We also generate a palette of non-covalent reversibly photoconvertible fluorescent proteins changing their fluorescence color upon illumination, a reversible behavior that still remains absent in regular fluorescent proteins. This light-induced color change opens the possibility to discriminate six spectrally similar **FAST** variants in live cells upon demonstrating the superiority of using multiple spectral channels for exploiting the time dependence of the fluorescence response to illumination.

## Introduction

Molecules experience multiple fates after light absorption.^1^ Among them, fluorescence and photoisomerization acting in tandem have found numerous attractive applications in advanced fluorescence bioimaging^2^ such as superresolution microscopy^3,4,5,6^ or dynamic contrast for multiplexed imaging.^7,8,9,10,11,12,13,14,15^ However, fluorescence emission and photoisomerization are competitive relaxation processes, the former generally being notably slower than the latter.^1^ Hence, to produce molecules that exhibit good performances with respect to both of them is highly demanding.

State-of-the-art reversibly photoisomerizable fluorophores experience a shift of the absorption spectrum and/or a major change of fluorescence emission upon illumination at one or two wavelengths. In reversibly photoswitching fluorophores, the change is restricted to fluorescence brightness but it does not significantly alter the emission spectrum.^16^ In contrast, illumination drives the exchange between two distinctly colored fluorescent states in reversibly photoconverting fluorophores.^17^

The reversibly photoisomerizable fluorophores generally involve a molecule engaged in the light-triggered isomerization of a photochromic motif (e.g. (*Z*)-(*E*) isomerization and/or ring opening-closure reaction) that is possibly followed by a thermally-driven reaction (e.g. protonation or aggregation).^16,17^ In reversibly switchable fluorescent proteins (RSFPs),^18,19,20^ a single chromophore covalently bound to the protein scaffold achieves to combine both photoisomerization and fluorescence emission. In contrast, in synthetic systems, the light-triggered molecule is not necessarily fluorescent. Hence, most realizations necessitate it to be linked with one (or more) fluorophores and to involve photoinduced energy or electron transfer to drive photoswitching or photoconversion of fluorescence. Therefore, the resulting multichromophoric system is rather large and its design and accessible color changes are photo(physical)chemically constrained.

Here, we introduce another strategy. Reversibly photoswitching fluorophores involve a host turning on fluorescence of a photoisomerizable fluorogenic guest, which gets liberated and dark upon illumination. When the medium further contains another fluorogenic guest acting as a competitor, illumination can change the color of fluorescence emission, thereby opening a path towards unprecedented non-covalent reversibly photoconvertible fluorescent tags.

Wash-free chemogenetic fluorescent systems involving non-covalent interaction between a biomolecular tag and a fluorogen dark in water are favorable to implement the latter strategy. Among hybrid systems combining synthetic fluorophores with genetically encoded tags for labeling of proteins^21^ and RNA,^22^ they are increasingly used in view of their labeling reversibility and continuous fluorogen replenishment, which are favorable for quantification and long duration observation.^23^ Originally reported in a few systems,^24,25,26,27,28^ fluorogen photoisomerization has recently benefited from a thorough theoretical and experimental study^29^ in the Fluorescence-activating and Absorption-Shifting Tag (**FAST**) series.^30,31^ In particular, it led us to identify a favorable kinetic regime of “photoejection” at low fluorogen concentration and high light intensity in which the limits of moderate fluorogen photoisomerization in the photostationary state are overcome to reach high fluorescence on/off ratios. Hence, we introduced **RSpFAST** combining **pFAST**^32^ with acidic fluorogens as a series of green non-covalent reversibly photoswitching fluorescent proteins (ncRSFPs) for enlarging the family of reversibly switchable fluorescent proteins (RSFPs).^18,19,20^

Here, we show that photoejection is not restricted to a few acidic fluorogens interacting with **pFAST**, but that it is a general dynamic behavior observed with multiple fluorogens when they interact with **FAST** variants such as **nirFAST**.^33^ Hence, we first deliver a large set of negative reversible photoswitchers, which turn off their fluorescence emission tailorable from the blue to the infrared wavelength range under illumination. We then expand the exploitation of photoejection. Overcoming the combination of the protein scaffold with a single photoisomerizable fluorogen,^32,36^ we harness pairs of fluorogens to turn **FAST** variants into various types of photoswitchers. Thus, we first generate a large palette of non-covalent reversibly photoconvertible fluorescent proteins (ncRCFPs) changing the color of their fluorescence emission upon illumination, a reversible behavior that still remains absent in fluorescent proteins.^18,34,35^ Further introducing a dark photoisomerizable fluorogen, we also produce positive reversible photoswitchers, that increase their brightness under illumination. As an application of ncRCFPs, we eventually use a pair of fluorogens to open the possibility to discriminate six spectrally similar **FAST** variants in living cells by exploiting the distinct time dependence of their fluorescence response to illumination in two spectral channels.

## Results and discussion

### The concept

When a fluorogen in its photoisomerized state **F’** exhibits a much lower affinity for the **FAST** scaffold **P** than in its initial thermodynamically stable state **F**, illumination of the **FAST**-**F** complex denoted **B** activates a photocycle, which involves continuous disruption of the **FAST**-**F’** complex denoted **B’**, and **F** recombination with the photoliberated **FAST** (Figure 1).^29^ Combined with a fluorogen (path a), **FAST** behaves as **RSFAST**, a reversible negative photoswitcher that turns from bright to dark upon illumination and thermally recovers its initial fluorescence. Its fluorescence signal exhibits maximal decay when the exchange between its free and bound states is slower than fluorogen photoisomerization (typically at light intensity higher than 0.1 E⋅m^-2^⋅s^-1^ -a few W.cm^-2^ - and fluorogen concentration lower than 1 μM).^29^ Here, **FAST** is further combined not anymore with *one* but with *two* fluorogens, which considerably expands the repertoire of light-driven fluorescence changes. Hence, thermally recovering reversibly photoconvertible fluorescent proteins (paths b,c) and reversible positive photoswitchers (path d) may become accessible as a consequence of the competition for occupying the tag complexing site (section 1.2 in SI).

**Figure 1.**
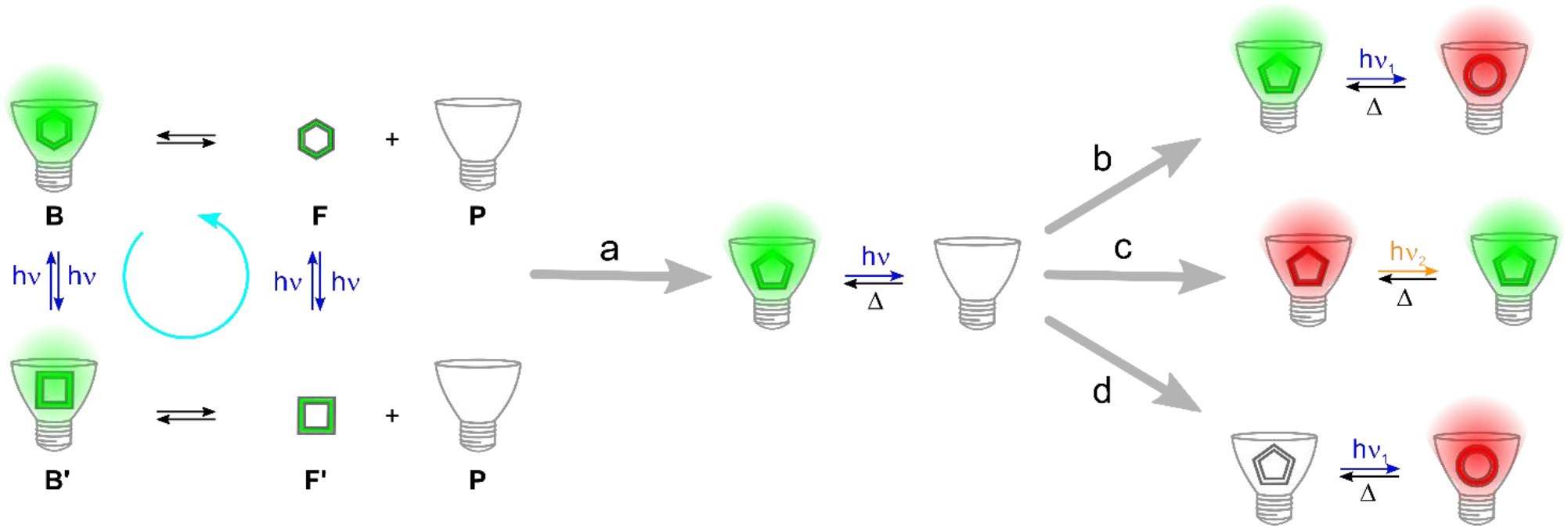
Non-covalent reversibly photoswitchable and photoconvertible fluorescent tags with thermal recovery for wash-free protein labeling. With one fluorogen, the four-state photocycle of **FAST**-based ncRSFPs reduces to a two-state reversible negative photoswitcher in the kinetic regime of photoejection (path a).^29^ Further exploiting photoejection but with two fluorogens, the combination of a reversibly photoisomerizable fluorogen giving a bright complex and either a non-photoisomerizable (path b) or a photoisomerizable (path c) one giving a bright complex of different color provides a reversibly photoconvertible fluorescent protein, whereas the combination of a reversibly photoisomerizable fluorogen giving a dark complex and a non-photoisomerizable one giving a bright complex (path d) yields a reversible positive photoswitcher. See also Figure S1 in SI.

### The screening step

To evaluate our concept, we submitted a large set of 15 reported^30,32,33,36,37^ and 7 newly synthesized fluorogens absorbing light over the whole UV-Visible wavelength range (Figure 2 and section 2.1 in SI) to a preliminary series of experiments: (i) **HBO3M, HBO35DM, HBT35DM**, and **HBR3CN** absorb at the near UV-Visible border and emit in the blue; (ii) **HBR3M, HBR25DM, HBR3Cl, HBR3Cl5F, HBR3F5OM, HBR35DF** absorb in the blue and emit in the green; (iii) **HBT35DOM** and **HBP35DOM** absorb in the cyan and emit in the yellow; (iv) **HBR35DOM** absorb in the green and emit in the orange; (v) **HPAR3OM** and **HPAR35DOM** absorb in the red and emit at the red-near infra-red border. We further produced fluorogens possessing lowered pK for enhanced **FAST** affinity (**HBT3CN, HBP3CN**, and **HBTH3CN** as blue emitters, and **HPAR3Cl, HPAR3F5OM** as red emitters) or low brightness (**HBIR3Cl, HBIR35DOM** as cyan and orange absorbers respectively).^29,32^

**Figure 2.**
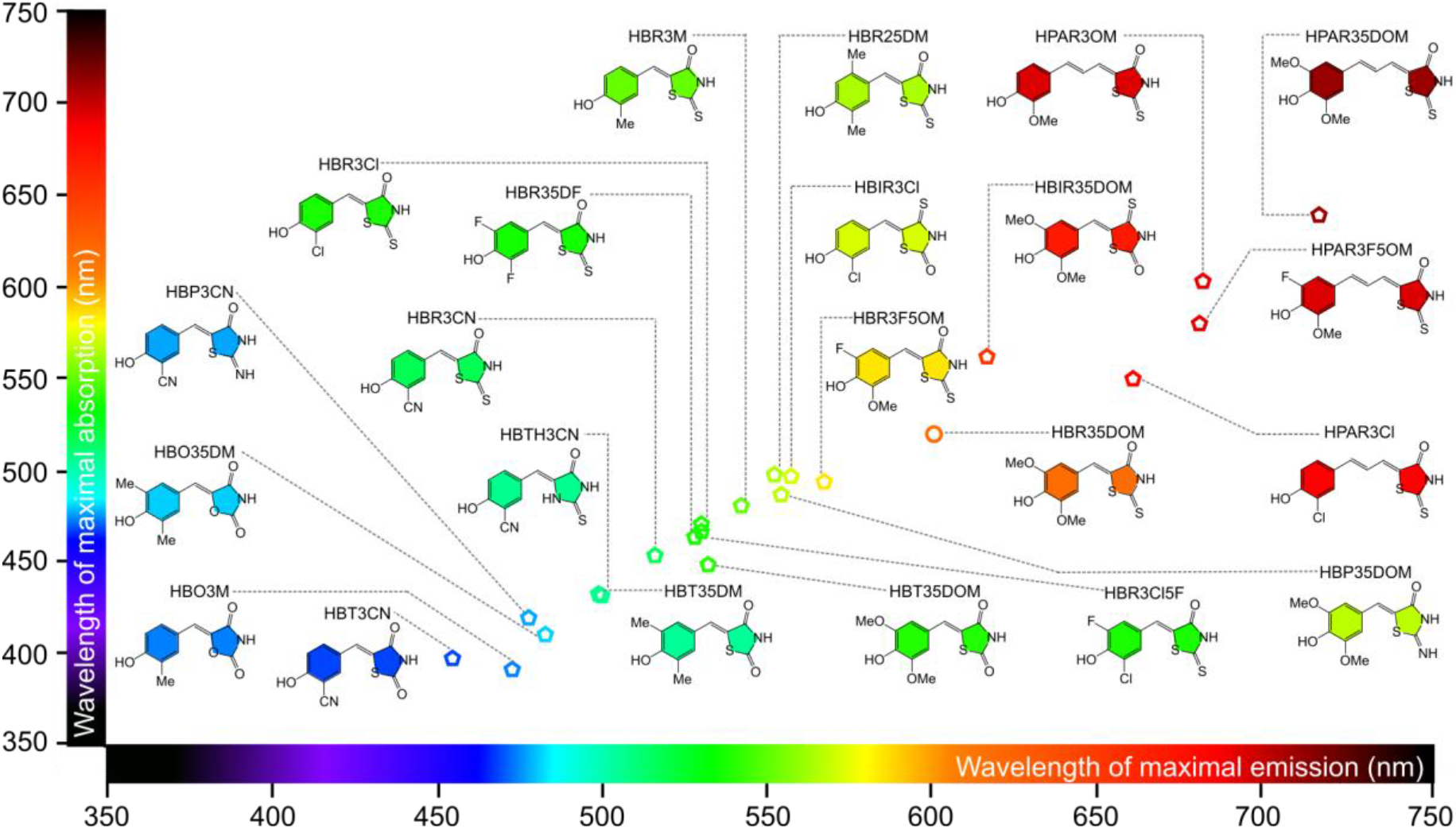
Palette of **FAST** fluorogens screened for reversible photoisomerization.

After recording their absorption and emission spectra (section 3.1 in SI), these 22 fluorogens have been assayed for reversible photoisomerization (section 3.2 in SI), thermal recovery (section 3.3 in SI), and proton exchange (section 3.4 in SI). Thus, we concluded that these visible range-spanning fluorogens photoisomerize (except **HBR35DOM**) with similar effective cross sections (sum of the cross sections for the forward and inverse transitions; 10^3^ m^2^.mol^-1^ range at the absorption maximum), thermally recover at a time scale ranging from less than 1 s to more than 1 h, and exhibit a proton exchange constant between 5 and 9. We then restricted the next experiments to a subset of representative fluorogens giving FAST complexes spanning the whole absorption/emission wavelength range (section 4.1 in SI) and measured 0.1-10 nM for the range of the dissociation constant of their complexes with **pFAST** and **nirFAST** (section 4.2 in SI).

### A series of negative FAST-based fluorescence photoswitchers

Photoejection of the photoisomerized fluorogen from its **pFAST** tag in labeled cells was previously^29^ observed at (i) fluorogen concentration in the 0.1-1 μM range and (ii) light intensity higher than 0.1 Ein.m^-2^.s^-1^ (a few W.m^-2^ range), which is widely encountered in optical microscopies. Anticipating a similar behavior for all the photoisomerizable fluorogens,^29^ we led a first series of experiments devoted to evidence photoejection and delimitate the experimental conditions of its observation. Confocal microscopy was used to image live HeLa cells expressing **H2B-pFAST** or **H2B-nirFAST** at the nucleus that were conditioned with fluorogens at 0.1-10 μM concentration (Table S8 in SI for the acquisition conditions).^29^ As displayed in Figures 3, S32, and S33, the fluorescence level at the nucleus exhibited fast monoexponential decrease (section 1.3 in SI) by up to 80 % upon applying repetitive scans in the presence of 0.1 μM of fluorogen for **HBT35DM, HBR3M**, and **HBR25DM** with **pFAST**, and for **HPAR3Cl, HPAR3OM**, and **HPAR35DOM** with **nirFAST**. As previously observed,^29^ photoejection reversibility was evidenced by essentially recovering initial fluorescence upon introducing a sufficient delay (typically 120 s) before applying a new series of repetitive scans. Moreover, the fluorescence drop originating from photoejection could be abolished by increasing the fluorogen concentration above 1 μM, where we only observed slow photobleaching, rejoining well-established experimental conditions for **FAST** labeling.^30,32^ Under such concentration conditions, the rate of **FAST**-Fluorogen recombination efficiently counterbalances the rate of light-induced disruption and one only observes a fluorescence decay caused by photobleaching.^29^

**Figure 3.**
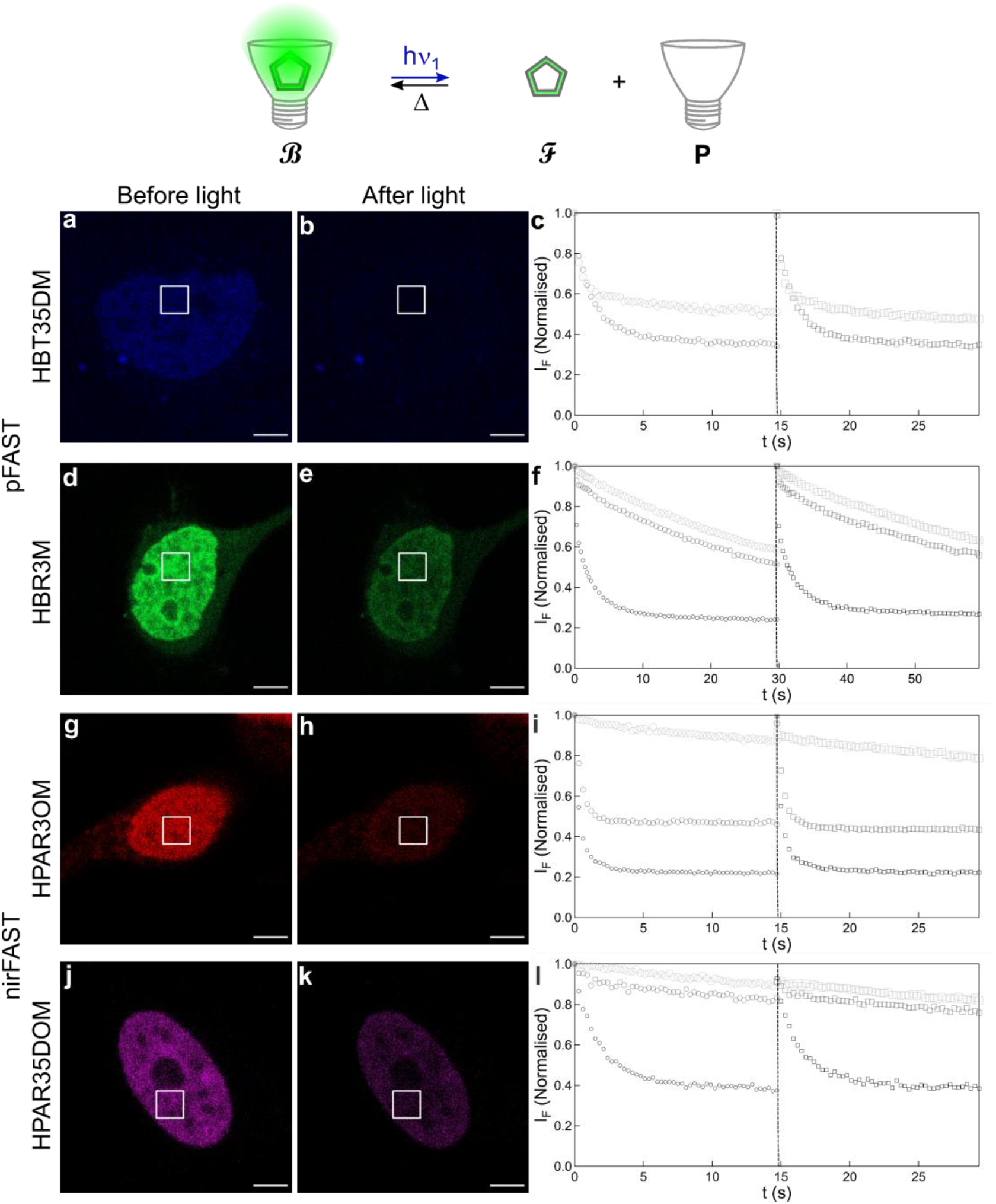
**FAST** to generate a palette of negative ncRSFPs. Time evolution of 445 (**a**-**c**), 488 (**d**-**f**), and 639 (**g**-**l**) nm light-induced fluorescence emission from **HBT35DM** (**a**-**c**), **HBR3M** (**d**-**f**), **HPAR3OM** (**g**-**i**), and **HPAR35DOM** (**j**-**l**) in **H2B-pFAST** (**a**-**f**) or **nirFAST** (**g**-**l**)-labeled live HeLa cells observed in confocal microscopy in the presence of the fluorogen in DMEM pH 7.4 (see also Figures S32 and S33). Photoejection is evidenced at 0.1 μM concentration from the contrast at the nuclei between the initial (**a**,**d**,**g**,**j**) and photosteady state (**b**,**e**,**h**,**k**) images, its reversibility by the similar loss of fluorescence normalized by its initial value after two consecutive series of frame acquisition (circles and squares markers after the 1^st^ and 2^nd^ series respectively) in the square region of interest at nuclei, and its concentration-dependence by the distinct decays observed at 0.1 (small black markers), 1 (medium grey markers), or 10 (large light grey markers) μM fluorogen concentration (**c**,**f**,**i**,**l**). Scale bar: 5 μm. *T* = 298 K. See Table S8 for the conditions of image acquisition.

### Optimizing the light-induced FAST fluorescence change in the presence of two fluorogens

Photoejection can further be exploited when **FAST** interacts with two fluorogens. Indeed, illuminating a first reversibly photoisomerizable fluorogen **ℱ**_***1***_ drives photoconversion of its thermodynamically stable (*Z*) to its (*E*) stereoisomer, which becomes less complexed by the tag in the 0.1-1 μM total fluorogen concentration range.^29^ It results in a backward shift of the extent of **ℱ**_***1***_ complexation, thereby liberating free tag sites now made available for complexation of a second fluorogen **ℱ**_***2***_, reversibly photoisomerizable or not. Hence, depending on the brightness, emission color, and thermokinetic properties of **ℱ**_***1***_ and **ℱ**_***2***_, illumination can generate various time evolutions of the **FAST** fluorescence (Figure 1).

Here, the concentrations of **ℱ**_***1***_ and **ℱ**_***2***_ must be properly fixed to benefit from important fluorescence changes. Rational optimization would primarily require collecting multiple thermokinetic constants for both fluorogens, which is demanding.^29^ In the absence of extensive kinetic information, we then used the respective dissociation constants *K*_d,1_ and *K*_d,2_ of the **FAST-**(*Z*)-Fluorogen complexes as guides for tuning the range of the concentrations of **ℱ**_***1***_ and **ℱ**_***2***_ (Tables S5 and S6). Indeed, fixing the fluorogen concentration to *K*_d_ optimizes both the complexation extent (governing the fluorescence level) and its change upon modifying the total fluorogen concentration (governing fluorescence sensitivity to illumination). Further expecting that endogenous cellular components would interact with the **FAST** fluorogens,^29^ we eventually ended up by exploring fluorogen concentrations ranging between 0.1 and a few μM for optimization.

### A palette of reversibly photoconvertible fluorescent proteins

To get reversibly photoconvertible fluorescent tags, we first used the strategy displayed in path b of Figure 1 and retained **HBR3Cl** (yielding a bright green **pFAST** complex) for the reversibly photoisomerizable fluorogen **ℱ**_***1***_, and **HBR35DOM** (giving a bright red complex with **pFAST**) for the non-photoisomerizable fluorogen **ℱ**_***2***_. Figure 4a1-a10 shows the time-lapse images of the nucleus of a 1:0.5 μM **HBR3Cl**:**HBR35DOM**-conditioned live HeLa cell expressing **H2B-pFAST**, which result from applying repetitive illumination scans in confocal microscopy. One observes a drop and a rise of the fluorescence emission of **ℱ**_***1***_ and **ℱ**_***2***_ respectively, that evidences successful implementation of photoconvertibility in the **FAST** system. As displayed in Figure 4b, the reversibility of photoconvertibility in **pFAST**-**HBR3Cl**:**HBR35DOM** was further evaluated by observing only 4 % and 9 % fluorescence loss in the green and red channels after 4 cycles of photoejection-90 s-long thermal recovery.

**Figure 4.**
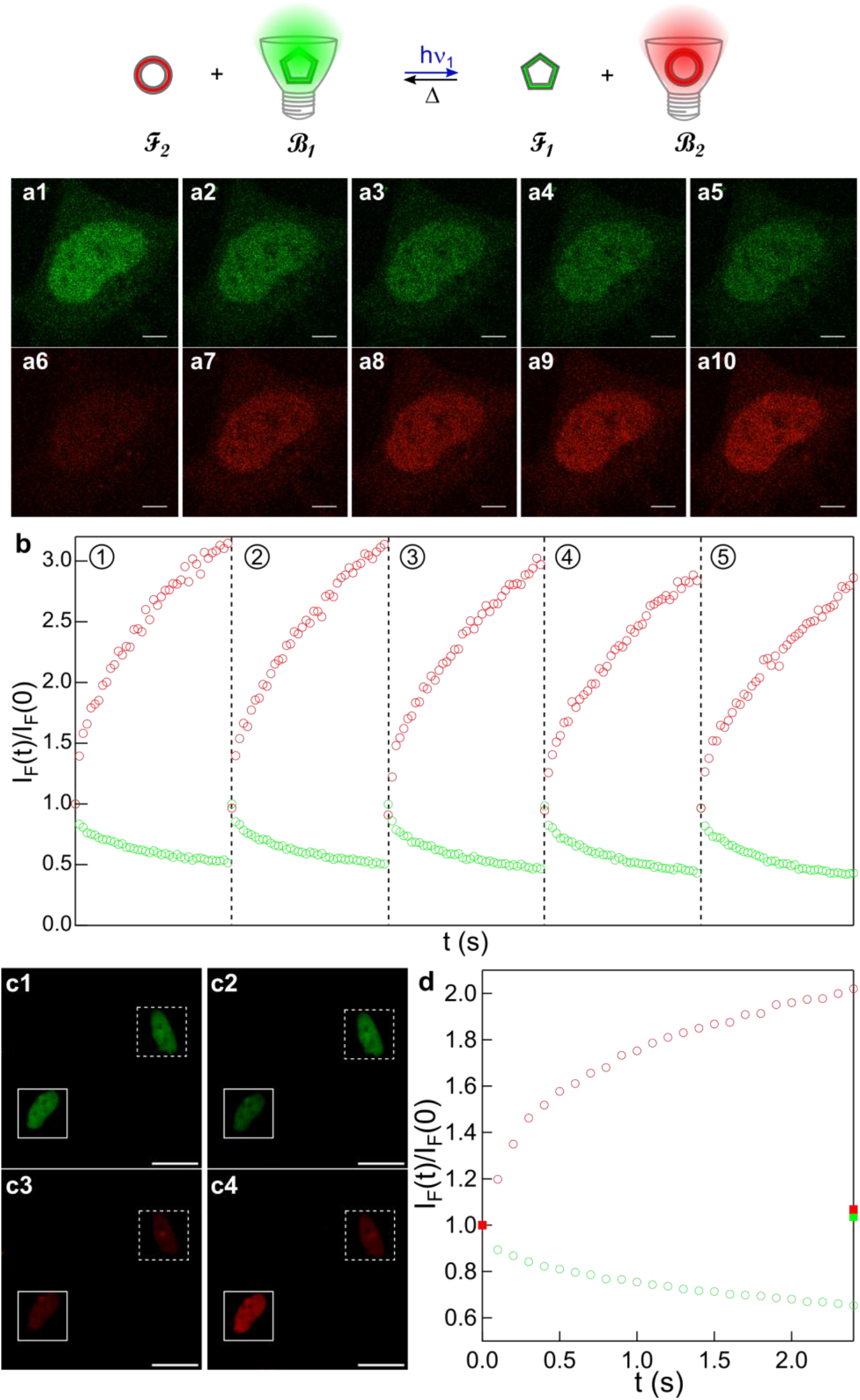
The **pFAST**-**HBR3Cl**:**HBR35DOM**: ncRCFP. **a1**-**a10**: Time-lapse of 488 nm light-induced change of fluorescence emission in the green (**a1-a5**) and red (**a6-a10**) channels of confocal microscopy from **H2B-pFAST**-labeled live HeLa cells conditioned with the **HBR3Cl**:**HBR35DOM** mixture of fluorogens in DMEM pH 7.4; **b**: Benchmarking of photostability of **pFAST**-**HBR3Cl:HBR35DOM.** Evolution of the normalized fluorescence intensity in the labeled nucleus shown in **a** over the course of five photoejection-thermal recovery cycles. Each vertical dashed line represents a 90 s thermal recovery period in the dark (not included in the time axis for compactness). The duration of each illumination period is 5.9 s; **c1**-**c4**: Selective photoconversion at the nucleus of a targeted **H2B-pFAST**-labeled live HeLa cell under local (solid white square) 470 nm illumination within a population. Local photoconversion is evidenced from comparing the initial (**c1**,**c3**) and final (**c2**,**c4**) fluorescence images in the green (**c1**,**c2**) and red (**c3**,**c4**) channels. The local drop and rise of the normalized fluorescence level at the targeted nucleus in the green (green circles) and red (red circles) channels (**d**) contrasts against the constant normalized fluorescence levels at the non-illuminated nucleus (green and red squares). Scale bar: 5 μm (**a1-a10**,**b**) or 28 μm (**c1-c4**). *T* = 298 (**a**,**c**) or 310 (**b**) K. See Table S8 for the conditions of image acquisition.

The 1:2 μM **HBR3Cl**:**HBR35DOM** mix was subsequently implemented in live HeLa cells expressing **H2B-pFAST** to selectively drive photoconversion at a targeted cell within a cell population by using a recently developed home-built microscope enabling us for patterned illumination.^38^ After recording the first image of two cells in the green (Figure 4c1) and red (Figure 4c3) fluorescence channels under uniform 470 nm illumination, we went on to selectively illuminate the nucleus of a single cell. We observed its fluorescence change with a drop and a rise in the green and red channels respectively (Figure 4d), ending up in a final spectral signature contrasting against the unchanged one of the non-illuminated nucleus (Figures 4c2,c4).

The strategy shown in path b of Figure 1 was then extended to three other mixtures involving **HBR35DOM** as the non-photoisomerizable fluorogen **ℱ**_***2***_ (Figure 5). Thus, we retained **HBT35DM** (yielding a blue **pFAST** complex), and **HPAR3OM** and **HPAR3F5OM** (giving both a bright near infra-red **nirFAST** complex) for the reversibly photoisomerizable fluorogen **ℱ**_***1***_. Then, we imaged in confocal microscopy live HeLa cells expressing **H2B-pFAST** that were conditioned with 10:0.25 μM **HBT35DM**:**HBR35DOM**, or **H2B-nirFAST** with 0.5:0.25 μM **HPAR3OM**:**HBR35DOM** and with 0.5:0.5 μM **HPAR3F5OM**:**HBR35DOM**. Similar changes of fluorescence normalized by its initial value were observed for all the investigated fluorogen pairs after consecutive series of frame acquisition separated by 90 s thermal recovery period in the dark, thereby validating generation of four reversibly photoconvertible fluorescent tags (Figures 5a-c5 and Figure S34).

**Figure 5.**
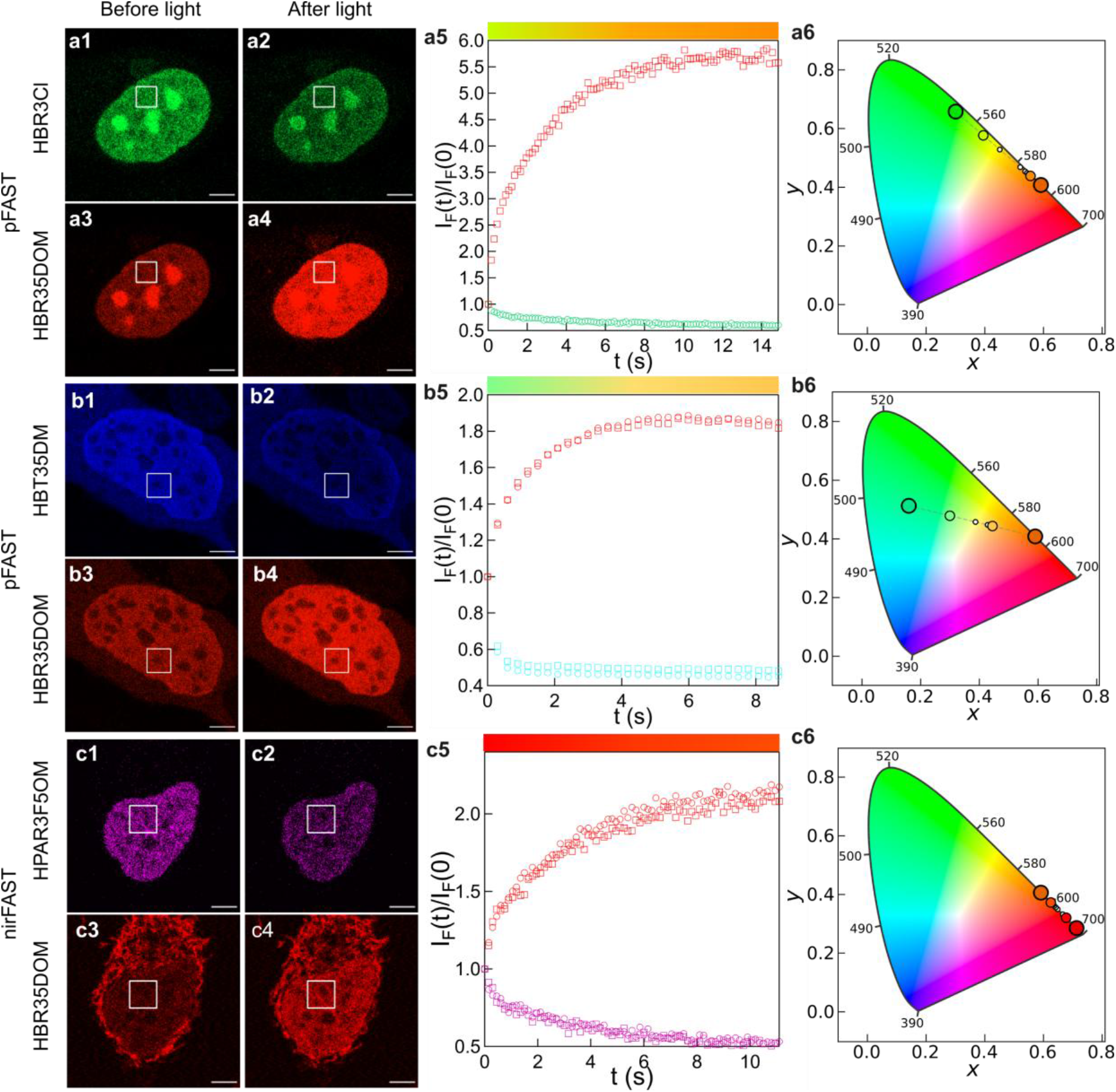
**FAST** to generate a palette of ncRCFPs. **a1**-**c6**: 488 (**a1**-**a6**), 445 (**b1**-**b6**), and 561 (**c1**-**c6**) nm light-induced changes of fluorescence emission in two wavelength channels of confocal microscopy from **H2B-pFAST** (**a1**-**b6**) or **nirFAST** (**c1**-**c6**)-labeled live HeLa cells conditioned with the **HBR3Cl**:**HBR35DOM** (**a1**-**a6**), **HBT35DM**:**HBR35DOM** (**b1**-**b6**), and **HPAR3F5OM**:**HBR35DOM** (**c1**-**c6**) mixtures of fluorogens in DMEM pH 7.4. The fluorescence change is evidenced from the contrast between the initial (**a1**,**a3**,**b1**,**b3**,**c1**,**c3**) and photostationary state (**a2**,**a4**,**b2**,**b4**,**c2**,**c4**) images of the nuclei and the time trace of the fluorescence normalized by its initial value in the region of interest at nuclei (**a5**,**b5**,**c5**), and its reversibility by the similarity of the time traces after two consecutive series of frame acquisition (circles and squares markers after the 1^st^ and 2^nd^ series respectively in **b5**,**c5**) separated by 90 s thermal recovery period in the dark. In the CIE 1931 color spaces (**a6**,**b6**,**c6**), the coordinates (x,y; SI Eq. (S42)) of the fluorescence emission of **HBT35DM, HBR3Cl, HBR35DOM**, and **HPAR3F5OM** are displayed as large circles, whereas the ones of the FAST labels conditioned with the fluorogen mixtures at initial and photosteady states are shown as medium size circles. The light-induced drifts of fluorescence color are linearly sampled with small circles along a dashed line. Scale bar: 5 μm. *T* = 298 K. See Table S8 for the conditions of image acquisition.

In fact, the time evolutions of the fluorescence level in two wavelength channels displayed in Figures 5a-c5 do not easily reflect the real change of fluorescence color at the cell nucleus. Hence, to make objective its light-driven change, we harnessed the CIE 1931 color space linking the visible spectrum and human color vision.^39^ To establish a reference, we first titrated **pFAST-HBT35DM** by **HBR35DOM** and evidenced that the progressive replacement of a fluorogen by another one delivered a linear trajectory in the color space (section 4.3 in SI). We then exploited the time evolution of the fluorescence level in both recorded wavelength channels of Figures 5a-c5 to compute the light-driven color shift at the labeled nucleus (section 1.3.7 in SI). Figures 5a-c6 together with the colored ribbons in Figures 5a-c5 demonstrate that the **FAST**-labeled nuclei encounter considerable changes of their color under their illumination.

To further enlarge the series of ncRCFPs, we then implemented the strategy involving a cocktail of two photoisomerizable fluorogens (path c of Figure 1). We first conditioned live HeLa cells expressing **H2B-nirFAST** with 1 μM **HBR3F5OM** as **ℱ**_***1***_ (yielding a bright green complex) and 1 μM **HPAR3OM** as **ℱ**_***2***_ (giving a bright red/infra-red fluorescent complex) and applied the following sequence of illuminations: (i) recording the initial frame at λ_1_ = 488 nm to image both **ℱ**_***1***_ and **ℱ**_***2***_; (ii) applying a set of repetitive scans at λ_2_ = 639 nm driving decay of **ℱ**_***2***_ fluorescence up to the photostationary state; (iii) recording the final frame at λ_1_ = 488 nm for imaging **ℱ**_***1***_ and **ℱ**_***2***_. In Figure S35, similar changes of the fluorescence color were observed between the initial and photostationary states for two consecutive series of frame acquisition separated by 90 s thermal recovery period in the dark: the **HBR3F5OM** fluorescence signal increased by 10 and 12 % and the **HPAR3OM** one decreased by 24 and 29 % during the first and second series of frame acquisition respectively.

### A series of positive fluorescence photoswitchers

When **ℱ**_***1***_ and **ℱ**_***2***_ give a dark photoisomerizable and a bright non-photoisomerizable **FAST** complex respectively, **ℱ**_***1***_ photoisomerization increases **ℱ**_***2***_ fluorescence, thereby generating a positive reversible photoswitcher (path d in Figure 1). For illustration, we adopted for **ℱ**_***1***_ **HBIR3Cl** (giving a dark **pFAST** complex), and first retained **HBR35DOM** for **ℱ**_***2***_. Figures 6a1-a5 show that applying in confocal microscopy sets of 514 nm light scans targeting **HBIR3Cl** photoisomerization separated by 90 s thermal recovery period in the dark reversibly doubled red fluorescence at the cell nucleus of live **H2B-pFAST** expressing-HeLa cells conditioned with 1:0.25 μM **HBIR3Cl**:**HBR35DOM**. We then again chose **HBIR3Cl** for **ℱ**_***1***_ (giving a dark **nirFAST** complex) and used **HPAR35DOM** for **ℱ**_***2***_ (yielding a bright red **nirFAST** complex with low absorbance at λ_1_ = 488 nm). Here, the sets of 488 nm light scans reversibly tripled the level of red/infra-red fluorescence at the cell nucleus of live **H2B-nirFAST** expressing-HeLa cells conditioned with 1:0.5 μM **HBIR3Cl**:**HPAR35DOM** as concluded from comparing the initial (Figures 6b1,b3) and final (Figures 6b2,b4) images recorded at 639 nm.

**Figure 6.**
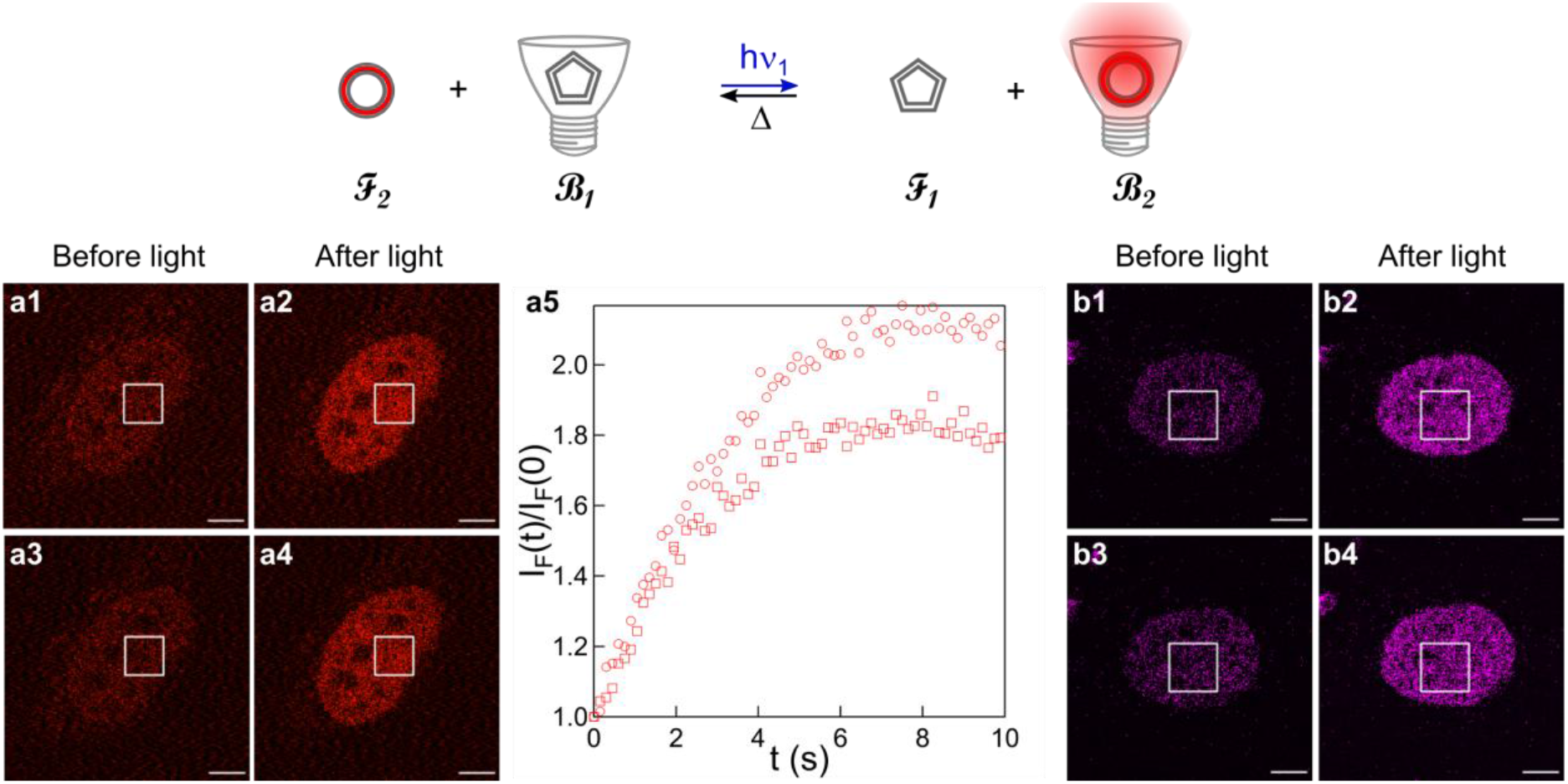
**FAST** to generate positive ncRSFPs. 514 (**a1**-**a5**) and 488 (**b1**-**b4**) nm light-induced changes of fluorescence images in confocal microscopy from **H2B-pFAST-** (**a1**-**a5**) and **H2B-nirFAST-** (**b1**-**b4**) labeled live HeLa cells respectively conditioned with the **HBIR3Cl**:**HBR35DOM** and **HBIR3Cl**:**HPAR35DOM** mixtures of fluorogens in DMEM pH 7.4. The fluorescence change is evidenced from the fluorescence contrast at the nuclei between the initial (**a1**,**a3**,**b1**,**b3**) and photosteady state (**a2**,**a4**,**b2**,**b4**) images, its reversibility by the similar changes of fluorescence normalized by its initial value in the region of interest after two consecutive series of frame acquisition (circles and squares markers after the 1^st^ and 2^nd^ series respectively in **a5**) separated by 90 s thermal recovery period in the dark. Scale bar: 5 μm. *T* = 298 K. See Table S8 for the conditions of image acquisition.

### Application towards multiplexed fluorescence imaging

When conditioned with a cocktail of two differently colored fluorogens, **FAST** variants are prone to deliver specific kinetic responses to illumination as time traces of the fluorescence level in the spectral channels used to follow the substitution of a fluorogen by the other within the tag cavity. Indeed, these traces report on a rich set of photophysical (i.e., absorption and emission spectra, quantum yields of fluorescence of the **FAST-Fluorogen** complexes), photochemical (i.e., cross sections of photoisomerization in the free and bound fluorogen states), and thermokinetic parameters (i.e., thermal rate constants of exchange between the four photocycle states) that are anticipated to depend on the **FAST** scaffold.

In order to evaluate the potential of such traces for multiplexed fluorescence imaging, we examined discrimination among a pool of live HeLa cells labeled at the nucleus with nine distinct **H2B-xFAST** (x=Y,^30^ p,^32^ nir,^33^ fr,^40^ t,^32^ green,^41^ i,^42^ o,^32^ Tsi^43^). After conditioning with 1:0.5 μM **HBR3Cl**:**HBR35DOM**, we acquired at 488 nm a sequence of confocal microscopy frames imaging the nuclei of ten to fifteen single cells from each **H2B-xFAST**-labeled sample. The images were then processed to retrieve the time evolution of the fluorescence response at the nucleus in the green and red spectral channels, mostly reporting on the fluorescence emission from **FAST-HBR3Cl** and **FAST-HBR35DOM** respectively. Figure 7a displays the sets of the time traces of the fluorescence levels in both color channels normalized to their first level and stacked sequentially. Except for **nirFAST**, fluorescence monotonously drops in the green channel whereas initial increases followed by slow decays are observed in the red channel. Noteworthily, the traces differ by their amplitude as well as by their time dependence.

**Figure 7.**
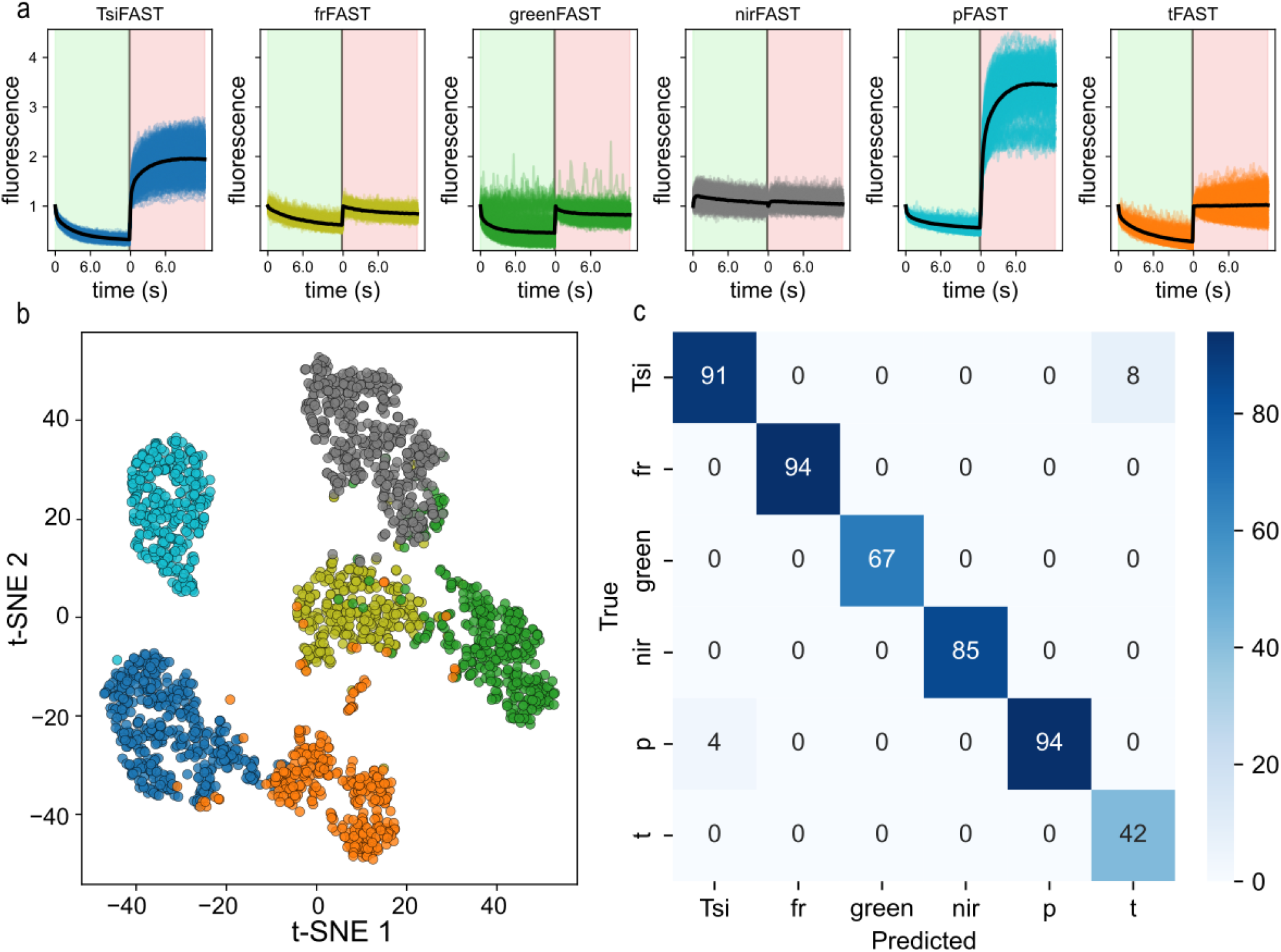
**rcFASTs** for multiplexed imaging. **a**: Combined response to 488 nm light in confocal microscopy of the normalized levels of fluorescence emission from the nuclei of live HeLa cells expressing **H2B-FAST** mutants (**TsiFAST, frFAST, greenFAST, nirFAST, pFAST, tFAST**) conditioned with 1:0.5 μM **HBR3Cl**:**HBR35DOM** in the 499-544 (green) and 588-694 (red) nm spectral channels; **b**: Dimension reduction from 150-dimension to 2-dimension to visualize the separability of the data. A supervised linear reduction of dimension of the whole dataset (Linear Discriminant Analysis, dimension 150 reduced to dimension 5) is followed by a non-linear dimension reduction technique (t-SNE, dimension 5 reduced to dimension 2); **c**: Confusion matrix associated with the 6 discriminated types of **FAST**-labeled. *T* = 298 K. See Table S8 for the conditions of image acquisition.

To refine this appreciation and conclude about its relevance for discriminating the **FAST** mutants, we evaluated the capacity of a trained machine learning model (Linear Discriminant Analysis) to distinguish the mutants (section 2.5.5 in SI). To artificially increase the dataset size, we split the cells into micro-regions and considered each as a data sample (2335 fluorescence traces in total). We divided the dataset into a training set and a test set. The test set corresponds to all the micro-regions retrieved from three cells out of the 10-15 cells for each protein. We tested different number of proteins, different normalization methods and data obtained from the green or red channel only, or combined channels. We concluded that we can discriminate up to six **FAST** mutants with an accuracy of 98% when considering the combined channels normalized at their first fluorescence level respectively. The tests on the normalization method allowed us to evidence that the relative amplitude of the signal evolution and not only the kinetics is a discriminant trait, and that combining the two channels presents an improvement in the classification. Combined channels allow to discriminate six **FAST** mutants whereas using a single channel limits the separability to four different **FAST** mutants for the green channel (accuracy 97 %) and three different **FAST** mutants for the red channel (accuracy 100 %, section 5.3 in the SI). Figure 7b shows a two-dimensional representation of the separability of the labelled proteins, while Figure 7c displays the confusion matrix on a test dataset which highlights predictions proposed by the trained algorithm, as well as errors.

## Conclusion

### From chemogenetic fluorescent labels to reversible negative photoswitchers

We previously evidenced that acidic fluorogens exhibit photoejection from **pFAST** at low concentration of fluorogen and high light intensity.^29^ Here, we showed that the photoejection scope expands to less acidic fluorogens. Indeed, it concerns essentially all the investigated fluorogens and **FAST** representatives. We also showed fluorogen concentration to be a key parameter determining the **FAST** behavior for protein labeling:^29^ in a regime of high light intensity below 1 μM, **FAST** acts as a reversible fluorescent photoswitcher. In contrast, photoejection is essentially abolished above 1 μM and **FAST** then acts as a fluorescent label now well-established.^30,32^ In fact, photoejection and shift of the labeling behavior may manifest themselves beyond **FAST** since other non-covalent chemogenetic systems (e.g., Spinach and Broccoli) exploit photoisomerizable donor-acceptor fluorogens.^44,45,46,47^

### A labeling mix strategy to generate reversibly photoswitchable and photoconvertible fluorescent protein tags

Fluorogen mixtures have never simultaneously been used for labeling a chemogenetic system. Here, we benefited from the **FAST** promiscuity^29,32,33^ and harnessed a mixture of two fluorogens emitting fluorescence in distinct wavelength ranges to tailor the color of their fluorescent complexes by implementing the principles of additive color synthesis. Interestingly, the collection of **FAST-**Fluorogen complexes delimits a large portion of the CIE 1931 color space, thereby establishing high potential to generate a variety of labeling color.

We jumped one step further by exploiting such fluorogen cocktails to expand state-of-the-art of the reversibly photoswitchable fluorescent protein tags. Hence, we illustrated that a proper choice of the fluorogen pair converts at will a same labeling tag into reversible negative or positive photoswitchers, as well as fluorescent proteins reversibly photoconverting from one color to another within a large palette. These reversible photoswitching or photoconversion of the **FAST** fluorescence are noteworthily transient with thermal recovery of the initial state occurring at a few tens of seconds time scale, a range shorter than most regular RSFPs.^11^ Whereas such a short range may be limiting for labeling a target within a population, it may be favorable for other applications (e.g., to apply multiplexed imaging protocols like OPIOM^48^).

### Rich photocycles for dynamic contrast

Any photocycle can be harnessed to kinetically discriminate reversibly photoactivatable fluorophores with protocols of dynamic contrast.^7,8,9,10,11,12,13,14,15^ However, its efficiency depends on the number of its light-driven and thermal steps and the span of its thermokinetic constants. Regular RSFPs exhibit rich photocycles involving multiple steps of light-driven and thermal isomerization, proton exchange, and formation of triplet states.^18,19,20^ Such a network of reactions is further expanded in **RCFASTs** from adding steps of association and dissociation of the fluorogen with the labeling tag where the fluorogen concentration acts as a control parameter for tuning the rate of fluorogen recombination.

### Enriching dynamic contrast with colors

Discrimination of fluorophores is generally achieved either in the spectral or in the temporal domain. Here, we illustrated that both discriminative dimensions can be fruitfully combined. When using two differently colored fluorogens, **rcFASTs** offer multiple control parameters (e.g. fluorogen concentrations, wavelengths of excitation and fluorescence collection) and observables (e.g. fluorescence levels in several wavelength channels) to exploit a rich kinetic set and generate discriminative fingerprints for highly multiplexed imaging.^49,50,51^

## Acknowledgements

This work was supported by the ANR (ANR-22-CE11-0011).

## Author contribution statements

Conceptualization: M. M., Y. S., A. L., I. A., A. G., L. J.; Methodology: M. M., Y. S., A. L., I. A., A. G., L. J.; Software: Y. S., A. L.; Validation: M. M., Y. S., A. L., I. A., T. L. S., A. G., L. J.; Formal analysis: M. M., Y. S., A. L., F. P., I. C., T. L. S., L. J.; Investigation: M. M., Y. S., E. L.; Resources: M. M., Y. S., A. L., F. P., L. E. H., I. C., M.-A. P., I. A.; Data Curation: M. M., Y. S., A. L.; Writing - Original Draft: M. M., Y. S., A. L., L. J.; Writing - Review & Editing: M. M., Y. S., I. A., A. G., L. J.; Visualization: M. M., Y. S., A. L., A. G., L. J.; Supervision: F. P., I. A., A. G., L. J.; Project administration: L. J.; Funding acquisition: L. J.

## Data availability

The datasets generated during and/or analyzed during the current study and that are not in the Supporting Information are available from the corresponding author on request.

## Supplementary information

Supplementary information reports on Theory, Materials and Methods, Investigation of the free fluorogens, Investigation of the **FAST**-fluorogen complexes, and **RSFAST** and **RCFAST** observations in microscopy.

## Table of Contents

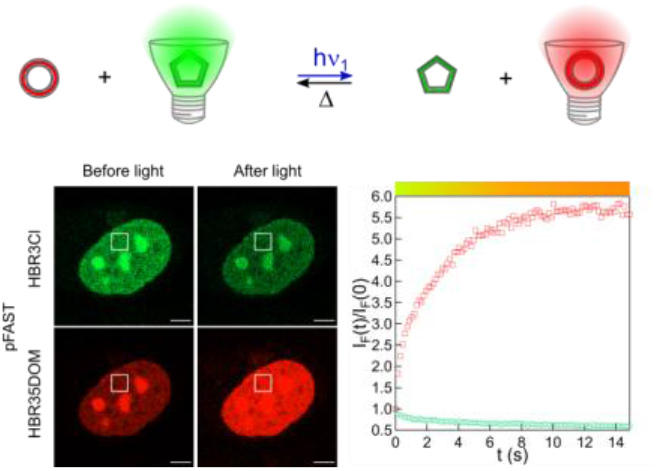

FAST proteins combined with one or two fluorogens undergo reversible fluorescence switching or photoconversion through photoisomerization. At low fluorogen concentration and high light intensity, they act as negative or positive photoswitchers and photoconvertible tags, which are promising to discriminate FAST variants.

