## Supporting Information for "Non-covalent reversibly photoconvertible fluorescent tags for wash-free protein labeling"

April 27, 2026

2.1.2.3 **(Z)-5-(3-cyano-4-hydroxybenzylidene)-2-iminothiazolidin-4-one 3c [HBP3CN]** . . 12
2.1.2.4 **(Z)-5-(3-cyano-4-hydroxybenzylidene)-2-thioxoimidazolidin-4-one 3d [HBTH3CN]** 12
2.5.3 Methods for acquisition of the thermokinetic and photophysical parameters of the free fluorogens 22

### 1 Theory

#### 1.1 Concept for non-covalent reversibly photoswitchable and photoconvertible fluorescent tags

In the following, we address several theoretical issues necessitated for the design and the implementation of our concept for accessing non-covalent reversibly photoswitchable and photoconvertible fluorescent tags with thermal recovery for wash-free protein labeling (Figure S1).

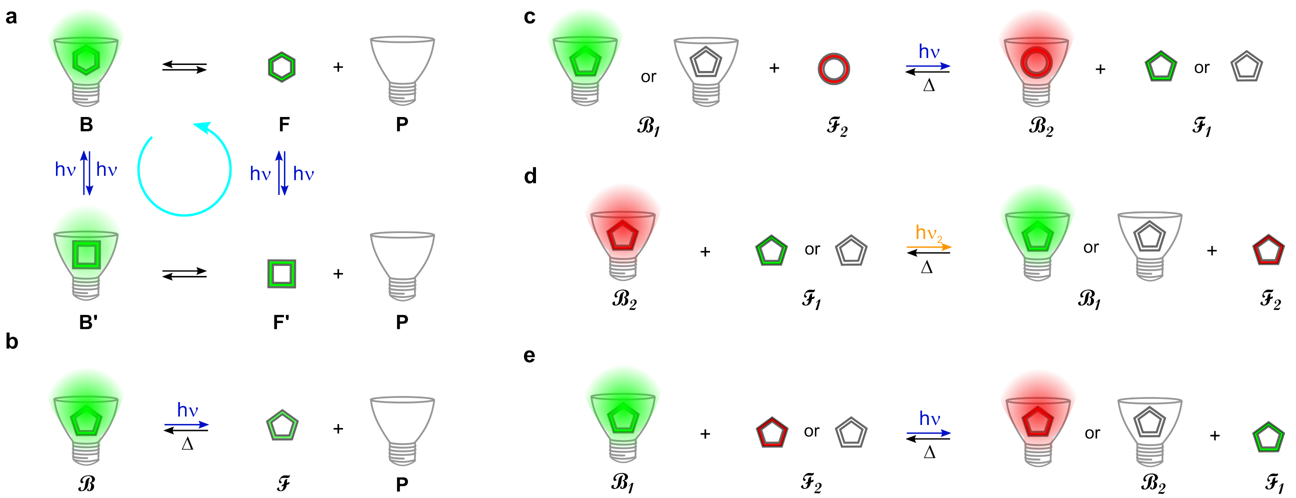

Figure S1: *Principle of non-covalent reversibly photoswitchable and photoconvertible fluorescent tags with thermal recovery for wash-free protein labeling.* **a**,**b**: The four-state photocycle of **FAST**-based ncRSFPs (**a**) yields a two-state reversible negative photoswitcher in the kinetic regime of photoejection (**b**); **c**–**e**: Combining a reversibly photoisomerizable fluorogen giving either a bright or dark complex and a non-photoisomerizable fluorogen (**c**), or a photoisomerizable fluorogen giving a bright complex with another photoisomerizable one giving either a bright or a dark complex of different color (**d**,**e**), provides a reversibly photoconvertible fluorescent protein (**c**,**d**,**e**), a reversible positive photoswitcher (**c**), or an enhanced reversible negative photoswitcher (**d**). In **c**–**e**, h$\nu_1$ and h$\nu_2$ refer to illumination at $\lambda_1$ and $\lambda_2$ in the absorption band of the complexes $\mathcal{B}_1$ and $\mathcal{B}_2$ respectively, where $\lambda_1 \leq \lambda_2$.

#### 1.2 Thermodynamic equilibrium in the absence of illumination

##### 1.2.1 Interaction between the protein scaffold and a single fluorogen

In this subsection, we consider that the fluorogen and the **FAST** protein scaffold interact to provide a fluorescent complex in the absence of illumination. One correspondingly adopts the thermodynamic model displayed in Figure S2 to analyze the thermodynamics of the interaction where the thermodynamic constant of association of **F** and **P** is denoted $K$ and

**F+P=B**

Figure S2: *Thermodynamic model of the* ***FAST-Fluorogen*** *interaction.*

the dissociation constant of the complex **B** is $K_d = 1/K$. Noting $F_{tot}$ and $P_{tot}$ the total concentrations of **F** and **P**, the equilibrium concentrations $F^\Delta(\infty)$, $P^\Delta(\infty)$, and $B^\Delta(\infty)$ of the three species **F**, **P**, and **B** are

$$B^\Delta(\infty) = \frac{[K(F_{tot}+P_{tot})+1] - \sqrt{[K(F_{tot}+P_{tot})+1]^2 - 4K^2F_{tot}P_{tot}}}{2K} \quad (S1)$$

$$F^\Delta(\infty) = F_{tot} - B(\infty) \quad (S2)$$

$$P^\Delta(\infty) = P_{tot} - B(\infty) \quad (S3)$$

The latter expression can be simplified when one component is in excess over the other:

- When $F_{tot} \gg P_{tot}$

$$B^{\Delta}(\infty) = \frac{KF_{tot}}{1+KF_{tot}} P_{tot} \tag{S4}$$

$$F^{\Delta}(\infty) = F_{tot} \tag{S5}$$

$$P^{\Delta}(\infty) = \frac{1}{1+KF_{tot}} P_{tot} \tag{S6}$$

- When $F_{tot} \ll P_{tot}$

$$B^{\Delta}(\infty) = \frac{KP_{tot}}{1+KP_{tot}} F_{tot} \tag{S7}$$

$$F^{\Delta}(\infty) = \frac{1}{1+KP_{tot}} F_{tot} \tag{S8}$$

$$P^{\Delta}(\infty) = P_{tot} \tag{S9}$$

Neglecting the brightness of the empty protein scaffold **P**, fluorescence intensity $I_F$ given in Eq.(S10) results from the contributions of the fluorogen **F** and its protein complex **B** with respective brightnesses $Q_F$ and $Q_B$

$$I_F = Q_F F^{\Delta}(\infty) + Q_B B^{\Delta}(\infty) \tag{S10}$$

###### 1.2.2 Interaction between the protein scaffold and two competing fluorogens

We now consider a mixture that involves a protein scaffold interacting with two competing fluorogens. We correspondingly adopted the thermodynamic model displayed in Figure S3 to analyze the thermodynamics of the interaction where the thermodynamic constant of association of $\mathbf{F}_i$ and **P** is denoted $K_i$ and the dissociation constant of the complex **B** is $K_{d,i} = 1/K_i$.

$\mathbf{F_1}$**+P=**$\mathbf{B_1}$
$\mathbf{F_2}$**+P=**$\mathbf{B_2}$

Figure S3: *Thermodynamic model of the interaction of* ***FAST*** *with two competing fluorogens.*

In the most general case, there is no analytic expression of the concentrations $F_1(\infty)$, $B_1(\infty)$, $F_2(\infty)$, and $B_2(\infty)$ at thermodynamic equilibrium. In order to overcome this limitation, we hypothesize that the experimental conditions are chosen such that $P \ll B_1 + B_2$. Then the system displayed in Figure S3 can be simplified to the thermodynamic model shown in Figure S4.

$\mathbf{F_2}$**+**$\mathbf{B_1}$**=**$\mathbf{B_2}$**+**$\mathbf{F_1}$

Figure S4: *Simplified thermodynamic model of the interaction of* ***pFAST*** *with two competing fluorogens.*

Denoting $F_{1,tot}$, $F_{2,tot}$, and $P_{tot}$ the total concentrations of $\mathbf{F}_1$, $\mathbf{F}_2$, and **P**, one then obtains the expressions (S11–S14) of the concentrations $F_1^{\Delta}(\infty)$, $B_1^{\Delta}(\infty)$, $F_2^{\Delta}(\infty)$, and $B_2^{\Delta}(\infty)$ at thermodynamic equilibrium.

$$B_1^{\Delta}(\infty) = \frac{-[(1-K)P_{tot} + KF_{2,tot} + F_{1,tot}] + \sqrt{[(1-K)P_{tot} + KF_{2,tot} + F_{1,tot}]^2 + 4(K-1)P_{tot}F_{1,tot}}}{2(K-1)} \tag{S11}$$

$$F_1^{\Delta}(\infty) = F_{1,tot} - B_1^{\Delta}(\infty) \tag{S12}$$

$$B_2^{\Delta}(\infty) = P_{tot} - B_1^{\Delta}(\infty) \tag{S13}$$

$$F_2^{\Delta}(\infty) = F_{2,tot} - P_{tot} + B_1^{\Delta}(\infty) \tag{S14}$$

where $K = \frac{K_2}{K_1}$.
Neglecting the brightness of the free fluorogens with respect to the brightness of their complexes with the protein, the fluorescence intensity from the system is given in Eq.(S15)

$$I_F(\infty) = Q_{B_1}B_1^{\Delta}(\infty) + Q_{B_2}B_2^{\Delta}(\infty) = Q_{B_2}P_{tot} + (Q_{B_1} - Q_{B_2})B_1^{\Delta}(\infty) \tag{S15}$$

#### 1.3 Response of a non-covalent reversibly photoswitchable fluorescent protein to light scanning in confocal fluorescence microscopy

Confocal fluorescence microscopy works by rapidly scanning a focused laser beam across the imaging area. Therefore, the illumination can no longer be considered constant and homogeneous. Then, the scanning parameters need to be considered for an accurate model of the evolution of the fluorescence signal from a non-covalent reversibly photoswitchable fluorescent protein at each pixel as a function of the number of scans/illumination time.

##### 1.3.1 Point spread function

In an inverted fluorescence microscope with a laser light source, the excitation and emission point spread functions (PSF) have non-trivial spatial distributions, but near the focus they can be approximated by Gaussian functions $\phi(r, w)$ with radial widths $w(z)$ dependent on the axial distance $z$ from the beam waist.

$$\begin{aligned} PSF_{ex}(r,z) &= \frac{w_{ex}^2(0)}{w_{ex}^2(z)}\,\phi(r, w_{ex}(z)) \\ PSF_{em}(r,z) &= \frac{w_{em}^2(0)}{w_{em}^2(z)}\,\phi(r, w_{em}(z)) \\ \phi(r,w) &= e^{-2r^2/w^2} \\ w(z) &= w_0\sqrt{1+\left(\frac{\lambda z}{\pi w_0^2}\right)} \end{aligned} \tag{S16}$$

where $\lambda$ is the light wavelength, $r$ is the radial distance from the axis of illumination in a given $z$ plane and $w_0$ is the radial beam waist (minimal beam radius obtained at the focus, defined at the radial distance at which light intensity is $1/e^2$ of the peak). The PSF as defined here are normalized by the peak light intensity $I_0$ at the focal point such that $I(r, z) = I_0 PSF(r, z)$.

In a confocal microscope, the emitted light is additionally passed through a pinhole before reaching the detector. Therefore, when a pinhole is employed, the detection PSF is narrower than the emission PSF, especially in the axial direction. The axial intensity follows a sinc function, but for small pinhole diameters ($\leq$ 1 Airy), it can be roughly approximated by a Gaussian with beam radius $w_{ax}$.

$$\begin{aligned} PSF_{det}(r,z) &\approx \phi(r, w_0)\phi(z, w_{ax}) \\ w_{ax} &\approx \frac{3.47n}{NA} w_0 \end{aligned} \tag{S17}$$

where $n$ is the refractive index of the imaging medium and $NA$ is the numerical aperture of the objective.

The final confocal point spread function $PSF_{conf}(r, z)$ is the product of the excitation and detection PSF. However, if the pinhole is not used ("widefield" conditions), the point spread function is $PSF_{wf}(r, z)$

$$\begin{aligned} PSF_{conf}(r,z) &= PSF_{ex}(r,z)\cdot PSF_{det}(r,z) \\ PSF_{wf}(r,z) &= PSF_{ex}(r,z)\cdot PSF_{em}(r,z) \end{aligned} \tag{S18}$$

##### 1.3.2 Photon exposure in scanning microscopy

In order to image an area, the laser beam is scanned across the sample row by row at a constant speed $v$, which can be calculated from the dwell time $t_d$ and the horizontal pixel size $\Delta x$ given by the instrument.

$$v = \Delta x / t_d \tag{S19}$$

We first consider the illumination from a single horizontal scanline at $y = 0$, with the focal plane at $z = 0$. To calculate the total photon exposure $H_0$ (in E·m$^{-2}$)[a] received at a given position $(x, y, z)$, we need to integrate the illumination PSF as it travels across the width $l$ of the sample. The total width is much larger than the width of the PSF, so we can consider it infinite if we discard the pixels at the edge of the image.

$$\begin{aligned} H_0(y,z) &= \int_{-l/2}^{l/2} \frac{I_0 PSF_{ex}(r,z)dx}{v} \approx \frac{I_0}{v}\int_{-\infty}^{+\infty} \frac{w_{ex}^2(0)\phi\left(\sqrt{x^2+y^2}, w_{ex}(z)\right)dx}{w_{ex}^2(z)} \\ &= \frac{I_0 w_{ex}^2(0)}{w_{ex}^2(z)v}\int_{-\infty}^{+\infty} e^{-2(x^2+y^2)/w_{ex}^2(z)}dx = \frac{\sqrt{\pi}I_0 w_{ex}^2(0)e^{-2y^2/w_{ex}^2(z)}}{\sqrt{2}w_{ex}(z)v} \\ &= \frac{\sqrt{\pi}w_{ex}^2(0)}{\sqrt{2}\Delta x w_{ex}(z)} I_0 t_d \phi(y, w_{ex}(z)) = \frac{\sqrt{\pi}w_{ex}(z)}{\sqrt{2}\Delta x} I_0 t_d PSF_{ex}(y,z) \end{aligned} \tag{S20}$$

The resulting equation shows that the amount of light received is homogeneous along the $x$ axis. Moreover a scaling factor $\frac{\sqrt{\pi}w_{ex}(z)}{\sqrt{2}\Delta x}$ dependent on the beam width and pixel size appears in addition to the "naive" expression of photon exposure $H_0 = I_0 t_d PSF_{ex}$.

An important parameter is the vertical inter-line distance $\Delta y$. Small values of $\Delta y$ cause the illumination PSF to partially overlap, exposing again the same area of the sample to the laser.

**1.3.2.1 Non-overlapping scan lines** We consider that the scan lines do not overlap, when $\Delta y \geq 3w_{ex}$. In that case, the total photon exposure $H$ is equal to $H_0$.

**1.3.2.2 Overlapping scan lines** On the other hand, if $\Delta y < w_{ex}$, the scan lines lie close to one another and the cumulative photon exposure $H$ can be calculated by adding up the contributions of all $N_y$ scan lines. Since $N_y \gg 1$, we can approximate the number of lines as being infinite if the lines near the edges of the image are excluded.

$$H = \sum_{n=1}^{N_y} H_0(y - n\Delta y, z) \approx \sum_{n\in\mathbb{Z}} H_0(y + n\Delta y, z) \tag{S21}$$

If $\Delta y \ll w_{ex}$, we can approximate further by considering that the overlapping lines form a continuum with a line density of $1/\Delta y$.

$$\begin{aligned} H &\approx \int_{-\infty}^{+\infty} \frac{H_0(y,z)}{\Delta y}dy = \frac{\sqrt{\pi}I_0 w_{ex}^2(0)}{\sqrt{2}\Delta x \Delta y w_{ex}(z)}\int_{-\infty}^{+\infty} e^{-2y^2/w_{ex}^2(z)}dy \\ &= \frac{\pi w_{ex}^2(0)}{2\Delta x \Delta y} I_0 t_d \end{aligned} \tag{S22}$$

In this case, the photon exposure is homogeneous in space within the limits of the image area.

##### 1.3.3 Photoisomerization in the protein cavity

The photoswitching and complexation of a fluorogen in the presence of **FAST** can be described with the 4-state model shown in Figure S5, which includes thermal and light-driven processes.
Illumination drives activation of a photocycle: (i) since **FAST P** has been selectively evolved to yield a complex **B** with the thermodynamically stable fluorogen configuration **F**, the resulting less stable complex **B'** between the photoisomerized fluorogen **F'** and **FAST P** would disrupt, thereby liberating free **FAST P** and the photoisomerized fluorogen **F'**; (ii) the latter would subsequently experience back photoisomerization to its thermodynamically stable state **F** and (iii) eventually regenerate the initial **FAST**-fluorogen complex **B**.

[a]In this manuscript, we provide the values of the light intensities in E.m$^{-2}$.s$^{-1}$ (or mol. of photons.m$^{-2}$.s$^{-1}$). This unit is currently used in actinometry. However, it is not often used in other fields such as optical microscopy, in which the researchers prefer to adopt W.m$^{-2}$. We provide the conversion in Eq.(S43).

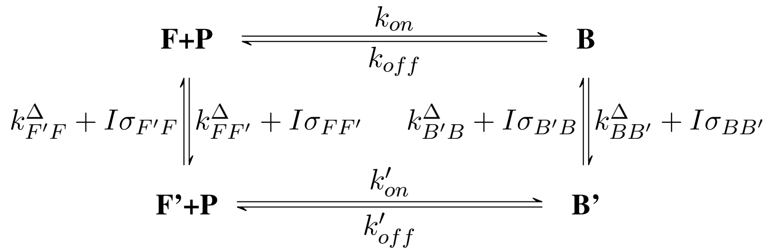

Figure S5: *Four-state model of the* ***FAST-Fluorogen*** *photocycle.* **F**, **F'**, **P**, **B** and **B'** represent free (*Z*)-**fluorogen**, free (*E*)-**fluorogen**, free **FAST** protein, (*Z*)-**fluorogen**:**FAST** complex, and (*E*)-**fluorogen**:**FAST** complex respectively. The thermal rate constants are denoted as $k^{\triangle}$ while $\sigma$ denotes a cross-section of photoswitching. $k_{on}$, $k_{off}$, $k'_{on}$ and $k'_{off}$ represent the on- and off-rate constants associated with the formation of the complexes **B** and **B'** respectively. Light intensity is denoted as $I$.

Considering the dwell time in confocal microscopy to be on the order of the microsecond, photoisomerization is the only reaction that can take place at this timescale in the **FAST**:**Fluorogen** system. The concentrations of **B** and **B'** at a given pixel after they were illuminated by the beam, $B_{1a}$ and $B'_{1a}$ respectively, can be calculated upon considering that only the thermodynamically stable isomer **B** is initially present. Then

$$\begin{aligned} B_{1a} &= B_0(a + (1-a)\alpha) = B_0 A \\ B'_{1a} &= B_0((1-a) - (1-a)\alpha) = B_0(1-A) \end{aligned} \tag{S23}$$

where $B_0$ is the initial concentration of **B** and

$$\begin{aligned} \alpha &= e^{-H\sigma} \\ \sigma &= \sigma_{BB'} + \sigma_{B'B} \\ a &= \frac{\sigma_{B'B}}{\sigma_{BB'} + \sigma_{B'B}} \\ A &= a + (1-a)\alpha \end{aligned} \tag{S24}$$

##### 1.3.4 Photoejection and recombination

After one full image is scanned, the process is repeated. During the time $t_f$ for frame acquisition, we chose experimental conditions such that photoejection and recombination of the fluorogen have enough time to take place. Then the concentrations of **B** and **B'** at a given pixel after this delay in the dark, $B_1$ and $B'_1$ respectively, can be calculated

$$\begin{aligned} B_1 &= b + (B_{1a} - b)\beta = B_{1a}\beta + b(1-\beta) \\ B'_1 &= b' + (B'_{1a} - b')\beta' \approx 0 \end{aligned} \tag{S25}$$

with

$$\begin{aligned} \beta &= e^{-(k_{on}F + k_{off})t_f} \\ \beta' &= e^{-(k'_{on}F' + k'_{off})t_f} \approx 0 \\ b &= \frac{FP_{tot}}{F + K_D} \approx P_{tot} = B_0 \\ b' &= \frac{F'P_{tot}}{F' + K'_D} \approx 0 \end{aligned} \tag{S26}$$

where $K_D$ and $K'_D$ refer to the dissociation constant of **B** and **B'**, and $P_{tot}$ designates the total concentration in **FAST**. Considering that $t_f$ is on the timescale of hundred of milliseconds for all the investigated fluorogens, $k'_{off}t_f \gg 1$, and consequently $\beta' \approx 0$. Moreover, under the concentration conditions for photoejection ($F' \ll K'_D$ and $F \gg K_D$), $b' \approx 0$ and $b \approx P_{tot} = B_0$. In other words, **B'** is completely ejected during the time between frames. Meanwhile $k_{on}F + k_{off}$ is small and **B** cannot reach its equilibrium concentration at the same time scale under those conditions.

###### 1.3.5 Repetition of the illumination sequence

The sequence of illumination and dark delay is repeated until a stationary state is reached, which can be described using a recursive definition of $B_n$, the concentration of **B** after the $n$th cycle.

$$\begin{aligned} B_n &\approx B_{n,a}\beta + B_0(1-\beta) = B_{n-1}A\beta + B_0(1-\beta) \\ &= B_0\left[(A\beta)^n + (1-\beta)\sum_{k=0}^{n-1}(A\beta)^n\right] \\ &= B_0\left[(A\beta)^n + \frac{(1-\beta)(1-(A\beta)^n)}{1-A\beta}\right] \\ &= B_0[B^* + (1-B^*)(A\beta)^n] \end{aligned} \tag{S27}$$

$$B^* = \frac{1-\beta}{1-A\beta}$$

The concentration of **B** at the photostationary state $B^*$ depends on light intensity and a first-order approximation can be made if $H\sigma \ll 1$.

$$\begin{aligned} B^* &= \frac{1-\beta}{1-\beta(a+(1-a)\alpha)} = \frac{1-\beta}{(1-a\beta)-\beta(1-a)e^{-H\sigma}} \\ &\approx \frac{1-\beta}{(1-a\beta)-\beta(1-a)(1-H\sigma)} = \frac{1}{1+\beta\frac{1-a}{1-\beta}\,H\sigma} \\ &\approx 1-\beta\frac{1-a}{1-\beta}\,H\sigma \end{aligned} \tag{S28}$$

At first order, the amplitude of photoejection is proportional to the photon exposure.
Transforming the equation for $B_n$, it becomes apparent that the function $B_n$ obeys an exponential decay law with apparent kinetic rate constant of photoejection $k_{PE}$.

$$B_n = B_0[B^* + (1-B^*)(A\beta)^n] = B_0[B^* + (1-B^*)e^{-k_{PE}n}] \tag{S29}$$

with

$$k_{PE} = -\ln A\beta > 0 \tag{S30}$$

The rate of photoejection depends on photon exposure according to the relationship:

$$k_{PE} = -\ln A\beta = -\ln[\beta(a+(1-a)\alpha)] = -\ln\left[\beta(a+(1-a)e^{-H\sigma})\right] \tag{S31}$$

If $H\sigma \ll 1$, or in other words if the extent of photoisomerization during each illumination cycle is small, we can perform a first-order approximation of $k_{PE}$.

$$\begin{aligned} k_{PE} &= -\ln\beta - \ln A \\ &\approx -\ln\beta - \ln(a+(1-a)(1-H\sigma)) \\ &= -\ln\beta - \ln(1+(a-1)H\sigma) \\ &\approx (1-a)H\sigma - \ln\beta \\ &= (1-a)H\sigma + (k_{on}F + k_{off})t_f \end{aligned} \tag{S32}$$

It becomes apparent that the rate of photoejection depends at first order on the photon exposure (and thus on light intensity, dwell time and pixel size) as well as concentration of free fluorogen.

##### 1.3.6 Fluorescence signal

Finally, we express the fluorescence signal $I_F$ originating from a point at $(x, y, z)$ and collected at a pixel with coordinates $(0, 0, 0)$ at frame $n$. The concentration of **B** and **B'** changes as the laser beam moves across the scanned line. However, if the light intensity and dwell time are such that that the extent of photoisomerization during one scan is small, $B_n(t)$ and $B'_n(t)$ remain approximately constant during the line scan. Furthermore, at the start of the scan **B** is the only fluorescent species present, so any contribution from **B'** can be ignored.

$$\begin{aligned} I_F &\approx I_0 Q_B B_n(t) PSF_{conf}(r, z) \\ &= I_0 Q_B B_0 \left[B^* + (1 - B^*)e^{-k_{PE}n}\right] PSF_{conf}(r, z) \end{aligned} \tag{S33}$$

##### 1.3.7 Retrieving information on the fluorescence hue from the detector signals

**1.3.7.1 Detector signals** We consider two fluorescent **FAST** complexes $\mathbf{B}_1$ and $\mathbf{B}_2$ with brightnesses $b_1(\lambda)$ and $b_2(\lambda)$ and concentrations $C_1$ and $C_2$ respectively, where $\lambda$ is the emission wavelength and the brightnesses are taken with respect to the same excitation wavelength $\lambda_{exc}$. Upon excitation with light intensity $I$, the total fluorescence spectral intensity $I_F(\lambda)$ (in units of Ein·m$^{-2}$·s$^{-1}$·nm$^{-1}$) is the sum of the contribution from $\mathbf{B}_1$ and $\mathbf{B}_2$:

$$I_F(\lambda) = (C_1 b_1(\lambda) + C_2 b_2(\lambda))I \tag{S34}$$

We consider the case when the fluorescence from the sample is detected in two channels. The detected fluorescence signals $S_1$ and $S_2$ depend on the fluorescence spectral intensity $I_F(\lambda)$ as well as the respective spectral sensitivities $s_1(\lambda)$ and $s_2(\lambda)$ of the two detection channels:

$$\begin{aligned} S_1 &= \int_\lambda I_F(\lambda) s_1(\lambda) d\lambda = I \int_\lambda (C_1 b_1(\lambda) + C_2 b_2(\lambda)) s_1(\lambda) d\lambda \\ S_2 &= \int_\lambda I_F(\lambda) s_2(\lambda) d\lambda = I \int_\lambda (C_1 b_1(\lambda) + C_2 b_2(\lambda)) s_2(\lambda) d\lambda \end{aligned} \tag{S35}$$

Rewriting in matrix form using the inner product notation $\langle \cdot, \cdot \rangle$

$$\langle f, g \rangle := \int_\lambda f(\lambda) g(\lambda) d\lambda \tag{S36}$$

$$\begin{bmatrix} S_1 \\ S_2 \end{bmatrix} = I \begin{bmatrix} \langle b_1, s_1 \rangle & \langle b_2, s_1 \rangle \\ \langle b_1, s_2 \rangle & \langle b_2, s_2 \rangle \end{bmatrix} \cdot \begin{bmatrix} C_1 \\ C_2 \end{bmatrix} \tag{S37}$$

**1.3.7.2 CIE coordinates** The tristimulus coordinates $X, Y, Z$ of a fluorescent (emissive) source with spectral power intensity E($\lambda$) (in units of W·m$^{-2}$·nm$^{-1}$) in CIE 1983 space are:

$$\begin{bmatrix} X \\ Y \\ Z \end{bmatrix} = \begin{bmatrix} \langle E, \bar{x} \rangle \\ \langle E, \bar{y} \rangle \\ \langle E, \bar{z} \rangle \end{bmatrix} \tag{S38}$$

Where $\bar{x}(\lambda), \bar{y}(\lambda), \bar{z}(\lambda)$ are the standard 2° color matching functions. Since this definition expects units of spectral power, we have to adjust it to use the spectral intensity $I_F(\lambda)$:

$$\begin{gathered} \begin{bmatrix} X \\ Y \\ Z \end{bmatrix} = \begin{bmatrix} \langle I_F, \bar{x}' \rangle \\ \langle I_F, \bar{y}' \rangle \\ \langle I_F, \bar{z}' \rangle \end{bmatrix} \\ \\ \bar{x}'(\lambda) = \bar{x}(\lambda) hcN_A/\lambda \\ \bar{y}'(\lambda) = \bar{y}(\lambda) hcN_A/\lambda \\ \bar{z}'(\lambda) = \bar{z}(\lambda) hcN_A/\lambda \end{gathered} \tag{S39}$$

Expanding $I_F(\lambda)$ and rearranging:

$$\begin{bmatrix} X \\ Y \\ Z \end{bmatrix} = \begin{bmatrix} \langle b_1, \bar{x}' \rangle & \langle b_2, \bar{x}' \rangle \\ \langle b_1, \bar{y}' \rangle & \langle b_2, \bar{y}' \rangle \\ \langle b_1, \bar{z}' \rangle & \langle b_2, \bar{z}' \rangle \end{bmatrix} \cdot \begin{bmatrix} C_1 \\ C_2 \end{bmatrix} \tag{S40}$$

Combining with Eq.(S37), we derive an equation for the CIE $X, Y, Z$ coordinates as a function of the observed detector signals $S_1, S_2$:

$$\begin{bmatrix} X \\ Y \\ Z \end{bmatrix} = \begin{bmatrix} \langle b_1, \bar{x}' \rangle & \langle b_2, \bar{x}' \rangle \\ \langle b_1, \bar{y}' \rangle & \langle b_2, \bar{y}' \rangle \\ \langle b_1, \bar{z}' \rangle & \langle b_2, \bar{z}' \rangle \end{bmatrix} \cdot \begin{bmatrix} \langle b_1, s_1 \rangle & \langle b_2, s_1 \rangle \\ \langle b_1, s_2 \rangle & \langle b_2, s_2 \rangle \end{bmatrix}^{-1} \cdot \begin{bmatrix} S_1 \\ S_2 \end{bmatrix} \tag{S41}$$

Finally, the CIE chromacity coordinates $x, y$ can be calculated from the $X, Y, Z$ coordinates:

$$\begin{bmatrix} x \\ y \end{bmatrix} = \frac{1}{X + Y + Z} \begin{bmatrix} X \\ Y \end{bmatrix} \tag{S42}$$

### 2 Materials and Methods

#### 2.1 Syntheses of the fluorogens

##### 2.1.1 General information

3-Chloro-4-hydroxybenzaldehyde, pseudothiohydantoin was purchased from Sigma Aldrich. 5-formyl-2- hydroxybenzonitrile was purchased from BLDpharm. 3-fluoro-4-hydroxy-5-methoxybenzaldehyde were purchased from abcr GmbH. Rhodanine, isorhodanine, 2,4-thiazolidinedione and thiohydantoin were purchased from Acros Organics. Commercially available reagents were used as obtained. $^1$H and $^{13}$C NMR spectra were recorded at 300 K on a Bruker AM 300 spectrometer; chemical shifts are reported in ppm with protonated solvent as internal reference ($^1$H, $CHD_2SOCD_3$ in $CD_3SOCD_3$ 2.50 ppm; $^{13}$C, $^{13}CD_3SOCD_3$ in $CD_3SOCD_3$ 39.5 ppm, $^{19}$F, $C_7H_5F_3$ in $CHD_2SOCD_3$ -63.7 ppm). Mass spectra high resolution were performed by the Service de Spectrométrie de Masse de Chimie ParisTech and the Service de Spectrométrie de Masse de l'Institut de Chimie Organique et Analytique (Orléans). Analytical thin-layer chromatography (TLC) was conducted on Merck silica gel 60 F254 pre-coated plates.

##### 2.1.2 Synthesis protocols

The synthesis pathways are displayed in Scheme 1.

**2.1.2.1 (E)-5-(3-chloro-4-hydroxybenzylidene)-4-thioxothiazolidin-2-one 3a [HBIR3Cl]** 4-Hydroxy-3-chloro benzaldehyde **1a** (140 mg, 0.9 mmol, 1.2 eq) and isorhodanine **2a** (100 mg, 0.75 mmol) were heated together at 170 °C. The mixture first fused and then solidified. After being cooled to 50–60 °C, the solid was stirred in ethanol at reflux until complete dissolution. Then an excess of water ($V_{H_2O} = 2 \times V_{EtOH}$) was slowly added and a precipitate appeared. It was filtered, washed with water, and dried over $P_2O_5$. **HBIR3Cl** was obtained as an orange powder (58 %, 118 mg). **$^1$H NMR (300 MHz, DMSO-d$_6$, $\delta$)**: 13.82 (br, 1H), 11.33 (br, 1H), 8.00 (s, 1H), 7.72 (d, J=3 Hz, 1H), 7.51(dd, J= 3, 9 Hz, 1H), 7.12 (d, J=9 Hz, 1H). **$^{13}$C NMR (75 MHz, DMSO-d$_6$, $\delta$):** 195.1, 170.5, 155.9, 135.1, 133.0, 130.7, 127.5, 125.7, 120.9, 117.4. **HRMS (ESI):** m/z calc. for $C_{10}H_7ClNO_2S_2$: 271.9601, found: 271.9601 $[M+H]^+$.

**2.1.2.2 (Z)-5-(3-cyano-4-hydroxybenzylidene)thiazolidine-2,4-one 3b [HBT3CN]** Same as **HBIR3Cl** using 5-formyl-2-hydroxybenzonitrile **1b** (50 mg, 0.34 mmol) and 2,4-thiazolidinedione **2b** (47 mg, 0.40 mmol, 1.2 eq). **HBT3CN** was obtained as a dark red powder (65%, 54 mg). **$^1$H NMR (300 MHz, DMSO-d$_6$, $\delta$):** 12.30 (s, br, 2H), 7.91 (d, J= 2.0 Hz, 1H), 7.72 (s, 1H), 7.69 (dd, J = 2.0, 8.8, 1H), 7.16 (d, J = 8.8 Hz, 1H). **$^{13}$C NMR (75 MHz, DMSO-d$_6$, $\delta$):** 167.8, 167.4, 161.6, 136.3, 135.3, 130.1, 124.7, 121.9, 117.3, 116.2, 100.1. **HRMS (ESI):** m/z calc. for $C_{11}H_5N_2O_3S$: 245.0023, found: 245.0031 $[M-H]^-$.

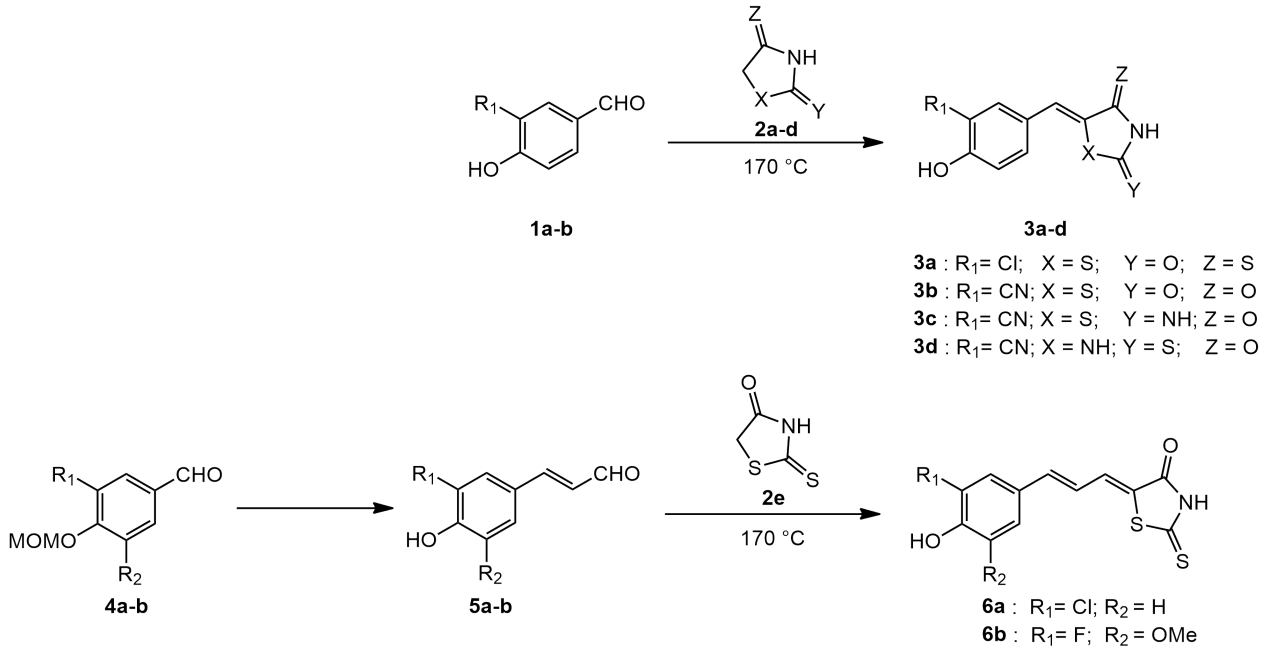

Scheme 1: Syntheses of the new fluorogens **3a-d** and **6a-b**.

**2.1.2.3 (Z)-5-(3-cyano-4-hydroxybenzylidene)-2-iminothiazolidin-4-one 3c [HBP3CN]** Same as **HBIR3Cl** using 5-formyl-2-hydroxybenzonitrile **1b** (50 mg, 0.34 mmol) and pseudothiohydantoin **2c** (46 mg, 0.40 mmol, 1.2 eq). **HBP3CN** was obtained as a dark red powder (36%, 30 mg). **$^1$H NMR (300 MHz, DMSO-d$_6$, *δ*):** 7.47 (d, J= 2.0, 1H), 7.36 (s, 1H), 7.31 (dd, J= 2.0, 9.0 Hz, 1H), 6.46 (d, J = 9.0 Hz, 1H). **$^{13}$C NMR (75 MHz, DMSO-d$_6$, *δ*):** 181.0, 175.0, 171.3, 136.3, 134.7, 129.7, 121.1, 120.8, 119.9, 117.1, 100.4. **HRMS (ESI):** m/z calc. for $C_{11}H_6N_3O_2S$: 244.0184, found: 244.0191 [M-H]$^-$.

**2.1.2.4 (Z)-5-(3-cyano-4-hydroxybenzylidene)-2-thioxoimidazolidin-4-one 3d [HBTH3CN]** Same as **HBIR3Cl** using 5-formyl-2-hydroxybenzonitrile **1b** (50 mg, 0.34 mmol) and thiohydantoin **2d** (46 mg, 0.40 mmol, 1.2 eq). **HBTH3CN** was obtained as a dark red powder (30%, 25 mg). **$^1$H NMR (300 MHz, DMSO-d$_6$, *δ*):** 12.3 (s, br, 3H), 8.1 (d, J = 2.0 Hz, 1H), 7.80 (dd, J= 2.0, 8.7 Hz, 1H), 7.02 (d, J = 8.7 Hz, 1H), 6.42 (s, 1H). **$^{13}$C NMR (75 MHz, DMSO-d$_6$, *δ*):** 178.9, 165.7, 161.2, 137.2, 134.5, 126.7, 123.8, 116.7, 116.6, 110.1, 99.8. **HRMS (ESI):** m/z calc. for $C_{11}H_6N_3O_2S$: 244.0181, found: 244.0188 [M-H]$^-$.

**2.1.2.5 ((Z)-5-((E)-3-(4-hydroxy-3-chlorophenyl)allylidene)-2 thioxothiazolidin-4-one 6a [HPAR3Cl]. 3-Chloro-4-(methoxymethoxy)benzaldehyde 4a** To a stirred solution of 3-chloro-4-hydroxybenzaldehyde (500 mg, 3.2 mmol) in anhydrous dichloromethane (5 mL) at 0 °C was added diisopropylethylamine (1.1 mL, 6.4 mmol, 2 eq) and chloromethyl methyl ether (0.36 mL, 4.8 mmol, 1.5 eq). The mixture was stirred overnight at room temperature. The reaction was quenched with a saturated aqueous solution of ammonium chloride, and the aqueous layer was extracted with dichloromethane. The combined organic layers were washed with brine, dried over magnesium sulfate and concentrated in vacuo. The desired compound **4a** was obtained quantitatively as a pale yellow solid and used without further purification for the next step. **$^1$H NMR (300 MHz, acetone-d$_6$, *δ*):** 9.92 (s, 1H), 7.95 (d, J = 3 Hz, 1H), 7.87 (dd, J = 3 and 9 Hz, 1H), 7.43 (d, J = 9 Hz, 1H), 5.44 (s, 2H), 3.50 (s, 3H).

**3-Chloro-4-hydroxycinnamaldehyde 5a** To a stirred solution of 3-chloro-4-(methoxymethoxy)benzaldehyde **4a** (250 mg, 1.2 mmol) and tributyl (1,3-dioxolan-2-ylmethyl) phosphonium bromide (915 mg, 2.4 mmol, 2 eq) in anhydrous tetrahydrofurane (5 mL) was added sodium hydride (60% dispersion in mineral oil) (100 mg, 2.4 mmol, 2 eq) and a catalytic amount of 18-crown-6 (64 mg, 0.24 mml, 0.2 eq). The reaction mixture was stirred at room temperature for the night. After addition of water, the mixture was extracted with ethyl acetate. The combined organic layers were concentrated in vacuo and the resulting residue was dissolved in THF/$H_2O$ (v/v : 5/1). TFA was added (2 mL) and

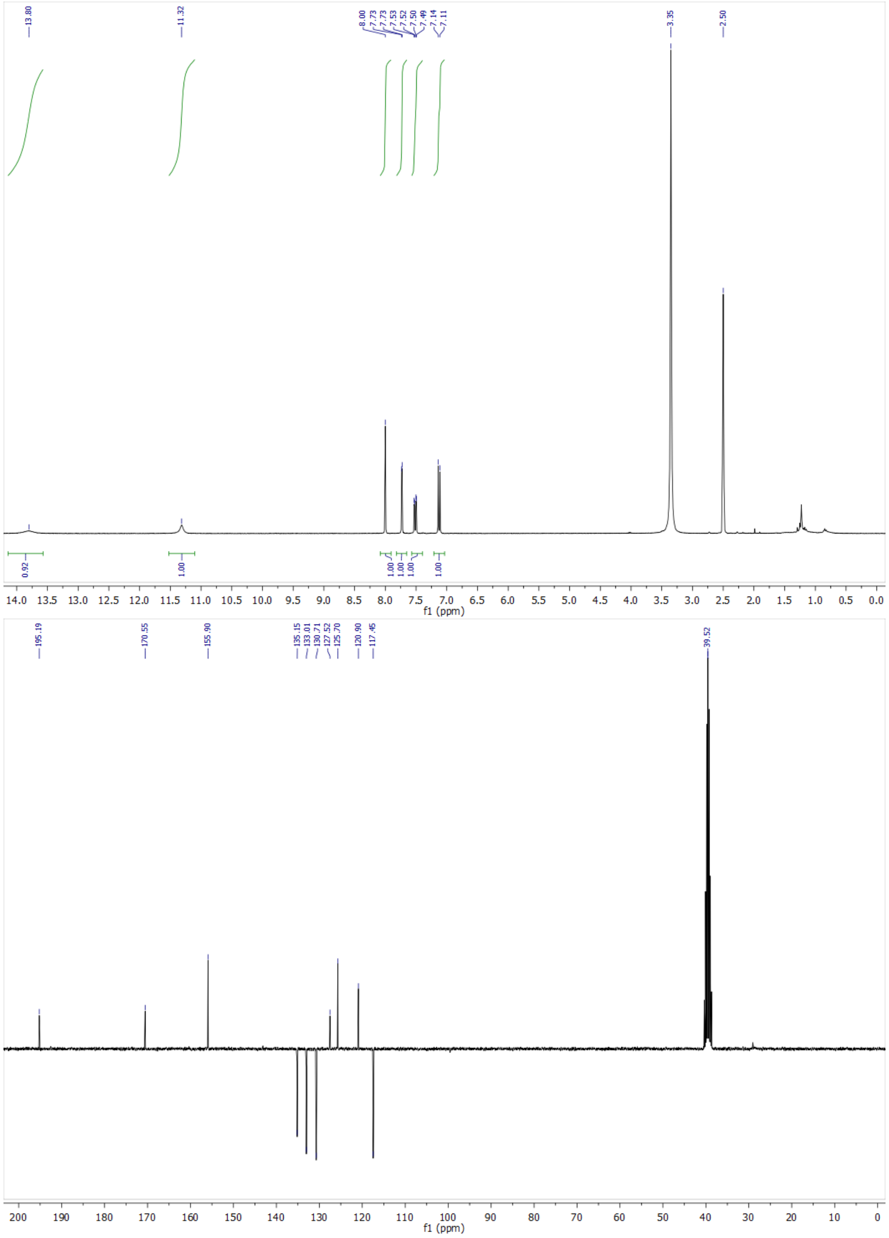

Figure S6: $^{1}$H-NMR and $^{13}$C-NMR spectra of **HBIR3Cl**.

the solution was stirred at room temperature until complete hydrolysis. The solution was extracted with ethyl acetate. The combined organic layers were dried over magnesium sulfate and concentrated. The crude was purified by flash chromatography on silica gel with cyclohexane/ethyl acetate (v/v : 8/2) as eluant. **5a** was obtained as a pale yellow solid (175 mg, 77% yield). **$^{1}$H NMR (300 MHz, acetone-d$_6$, $\delta$):** 9.67 (d, J = 9Hz, 1H), 7.78 (d, J = 3 Hz, 1H), 7.58 (d, J = 15 Hz, 1H), 7.58 (dd, J = 3 and 9 Hz, 1H), 7.11 (d, J = 9 Hz, 1H), 6.68 (dd, J = 9 and 15 Hz, 1H) ; **$^{13}$C NMR (75 MHz, DMSO-d$_6$, $\delta$):** 193.9, 155.8, 152.1, 130.4, 129.1, 126.6, 126.4, 120.5, 116.9. **HRMS (ESI):** m/z calc. for $C_9H_6ClO_2$: 181.0136, found: 181.0064 [M-H]$^-$.

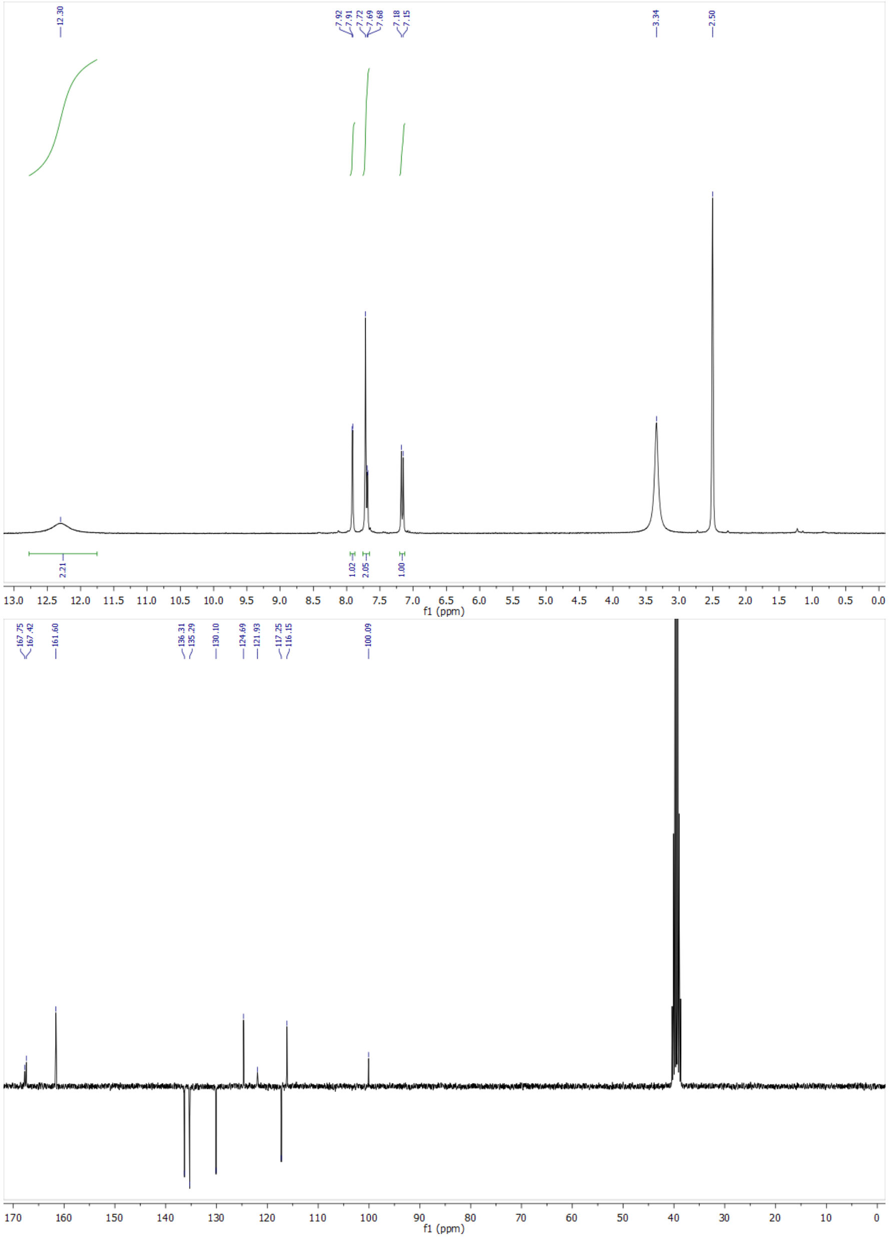

Figure S7: $^{1}$H-NMR and $^{13}$C-NMR spectra of **HBT3CN**.

To a stirred solution of rhodanine **2e** (150 mg, 1.1 mmol, 1.5 eq) and 4-hydroxy 3-chlorocinnamaldehyde (135 mg, 0.74 mmol) in ethanol (8 mL) was added 4-dimethylaminopyridine (14 mg, 0.11 mmol, 0.15 eq). The solution was stirred at reflux for 24 h. The mixture was neutralized by adding 1 M HCl (16 mL). After cooling to 4 °C and standing overnight, the resulting precipitate was filtered, washed with a cold solution EtOH/$H_2O$ (v/v : 1/5). After vacuum drying over $P_2O_5$, **HPAR3Cl** was obtained as a red powder (195 mg, 88% yield). **$^{1}$H NMR (300 MHz, acetone-d$_6$, $\delta$):** 7.77 (d, J = 3 Hz, 1H), 7.52 (dd, J = 3 and 9 Hz, 1H), 7.34 (dd, J = 1 and 12 Hz, 1H), 7.23 (bd, J = 15 Hz, 1H), 7.07 (d, J = 9 Hz, 1H), 6.93 (dd, J = 12 and 15 Hz, 1H)) ; **$^{13}$C NMR (75 MHz, DMSO-d$_6$, $\delta$):** 195.2, 168.6, 154.8, 143.8, 132.5, 129.3, 128.7, 128.1, 125.7, 121.9, 120.5, 116.8. **HRMS (ESI):** m/z calc. for $C_{12}H_7ClNO_2S_2$: 295.9687, found: 295.9615 [M-H]$^-$.

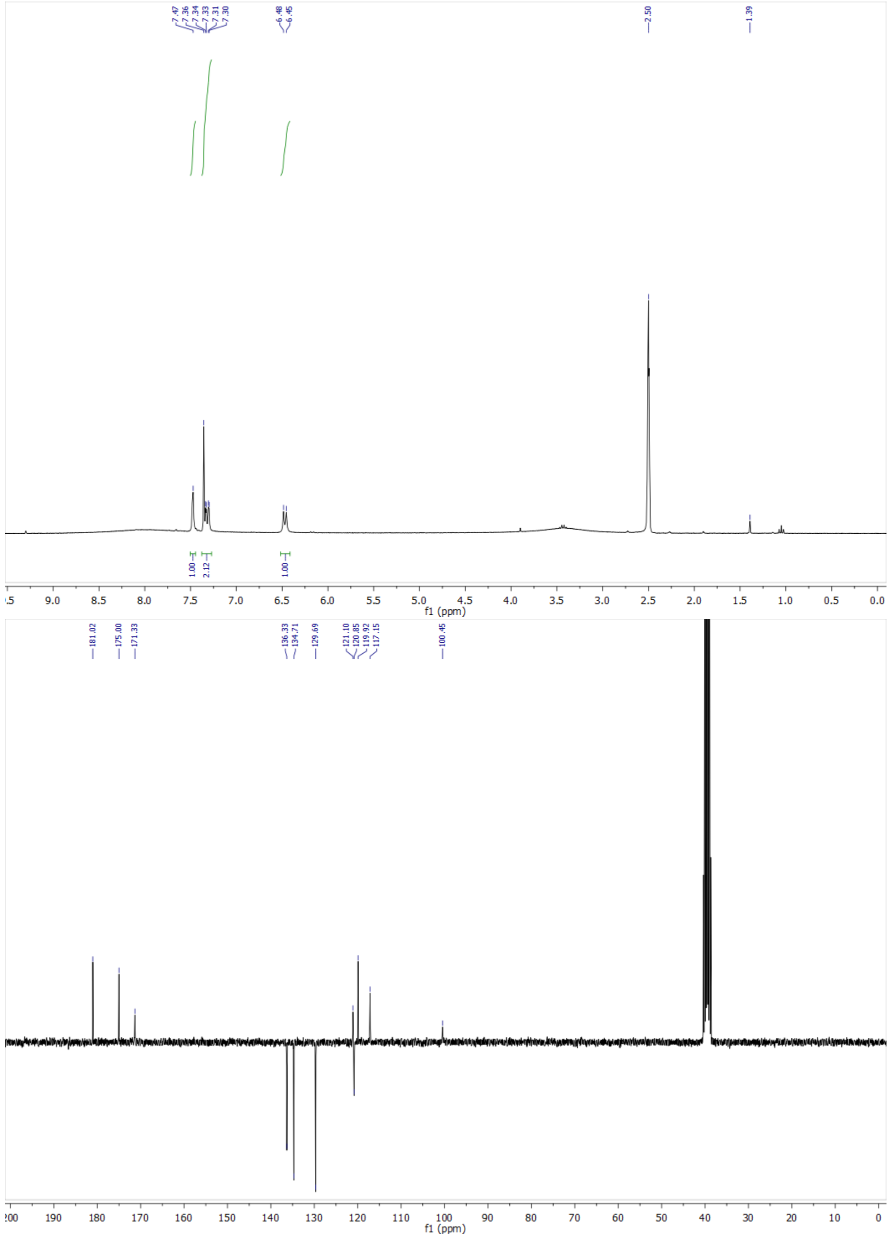

Figure S8: $^{1}$H-NMR and $^{13}$C-NMR spectra of **HBP3CN**.

###### 2.1.2.6 ((Z)-5-((E)-3-(4-hydroxy-3-fluoro-5-methoxyphenyl)allylidene)-2 thioxothiazolidin-4-one 6b [HPAR3F5OM].

**3-Fluoro-4-(methoxymethoxy)-5-methoxybenzaldehyde 4b** Same as **4a** using 3-fluoro-4-hydroxy-5-methoxybenzaldehyde (544 mg, 3.2 mmol). The desired compound **4b** was obtained quantitatively as a pale yellow viscous oil and used without further purification for the next step. **$^{1}$H NMR (300 MHz, acetone-d$_6$, $\delta$):** 9.92 (d, 1H, J= 3 Hz), 7.45 (dd, J = 3 Hz, 1H), 7.38 (dd, J = 3 and 12 Hz, 1H), 7.43 (d, J = 9 Hz, 1H), 5.22 (s, 2H), 3.99 (s, 3H), 3.54 (s, 3H).

**3-Fluoro-4-hydroxy-5-methoxycinnamaldehyde 5b** Same as **5a** using 3-Fluoro-4-(methoxymethoxy)-5-methoxybenz aldehyde **4b** (600 mg, 2.8 mmol), tributyl (1,3-dioxolan-2-ylmethyl) phosphonium bromide (2 g, 5.6 mmol, 2 eq), sodium hydride (60% dispersion in mineral oil) (224 mg, 5.6 mmol, 2 eq) and 18-crown-6 (148 mg, 0.56 mml, 0.2 eq) in dry tetrahydrofurane (15 mL). After purification by flash chromatography on silica gel with cyclohexane/ethyl acetate (v/v : 8/2) as eluant. **5b** was obtained as a pale yellow solid (350 mg, 64% yield). **$^{1}$H NMR (300 MHz, acetone-d$_6$, $\delta$):** 9.65

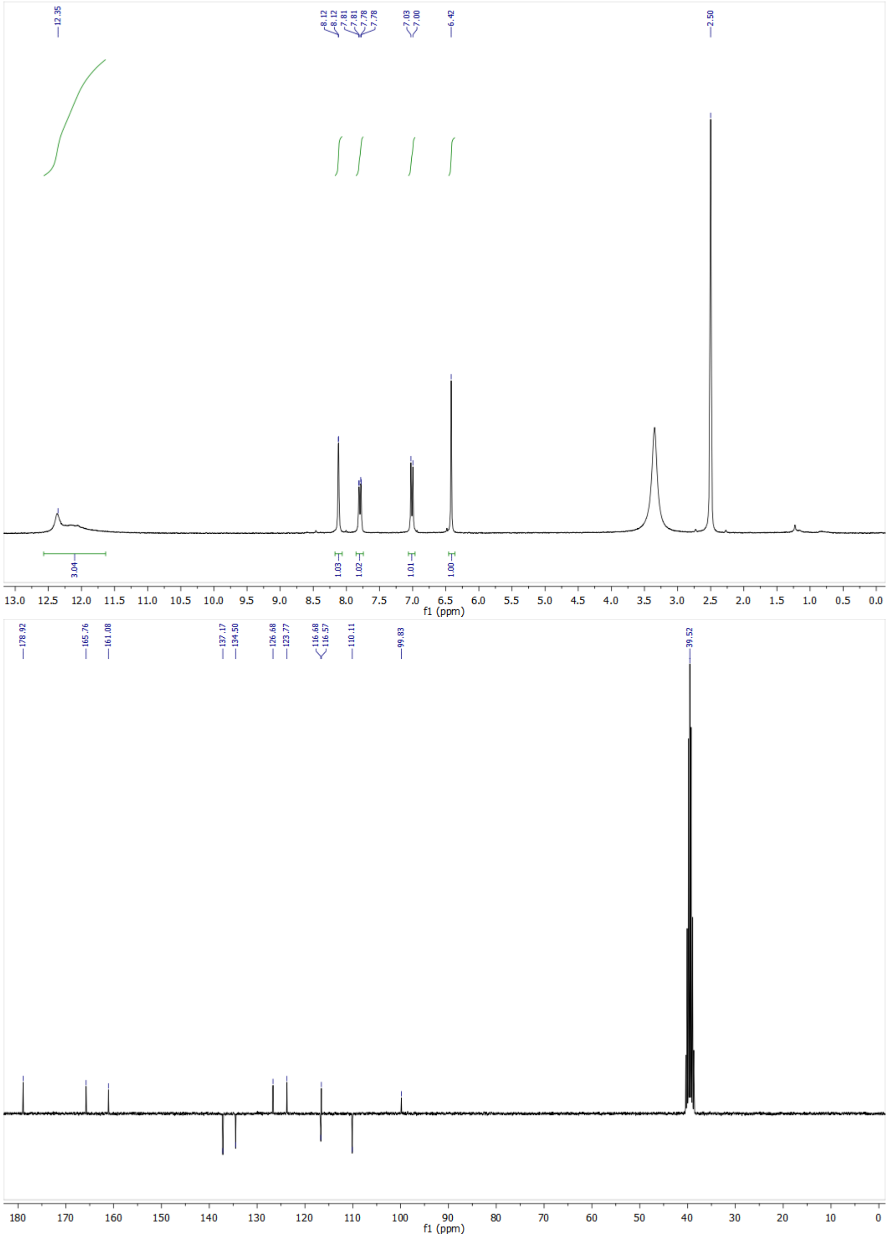

Figure S9: $^{1}$H-NMR and $^{13}$C-NMR spectra of **HBTH3CN**.

(d, J = 9Hz, 1H), 7.56 (d, J = 15 Hz, 1H), 7.58 (d, J = 15 Hz, 1H), 7.58 (dd, J = 3 and 9 Hz, 1H), 7.11 (d, J = 9 Hz, 1H), 6.68 (dd, J = 9 and 15 Hz, 1H). **$^{13}$C NMR (75 MHz, DMSO-d$_6$, $\delta$):** 194.0, 152.8 (d, JCF = 2 Hz), 151.2 (d, JCF = 237 Hz), 149.6 (d, JCF = 7 Hz), 137.5 (d, JCF = 14 Hz), 127.1, 124.7 (d, JCF = 9 Hz), 109.8 (d, JCF = 19 Hz), 108.2, 56.4. **$^{19}$F NMR (282 MHz, DMSO-d$_6$, $\delta$):** -137.95 (s, 1F). **HRMS (ESI):** m/z calc. for $C_{10}H_8FO_3$: 195.0537, found: 195.0464 [M-H]$^-$.

Same as **HPAR3Cl** using rhodanine **2e** (180 mg, 1.33 mmol, 1.5 eq), 3-Fluoro-4-hydroxy-5-methoxycinnamaldehyde (175 mg, 0.89 mmol) and 4-dimethylaminopyridine (16 mg, 0.13 mmol, 0.15 eq) in ethanol (8 mL). **HPAR3F5OM** was obtained as a red powder (240 mg, 88% yield). **$^{1}$H NMR (300 MHz, DMSO-d$_6$, $\delta$):** 13.56 (br, 1H), 9.84 (s, 1H), 7.30 (d, J = 12 Hz, 1H), 7.21 (dd, J = 3Hz and JHF = 12 Hz, 1 H), 7.19 (d, J = 15Hz, 1H), 7.14 bs, 1H), 6.93 (dd, J = 12 and 15 Hz, 1H), 3.87 (s, 3H) ; **$^{13}$C NMR (75 MHz, DMSO-d$_6$, $\delta$):** 195.1, 168.6, 151.2 (d, JCF = 248 Hz), 149.7 (d, JCF = 5 Hz), 144.5 (d, JCF = 2 Hz), 136.7 (d, JCF =14 Hz), 132.4, 126.4 (d, JCF =9 Hz), 125.7, 122.3, 108.9 (d, JCF = 19 Hz), 107.6, 56.4 ; **$^{19}$F NMR (282 MHz, DMSO-d$_6$, $\delta$):** -137.97 (s, 1F). **HRMS (ESI):** m/z calc. for $C_{13}H_9FNO_3S_2$: 310.0087, found: 310.0015 [M-H]$^-$.

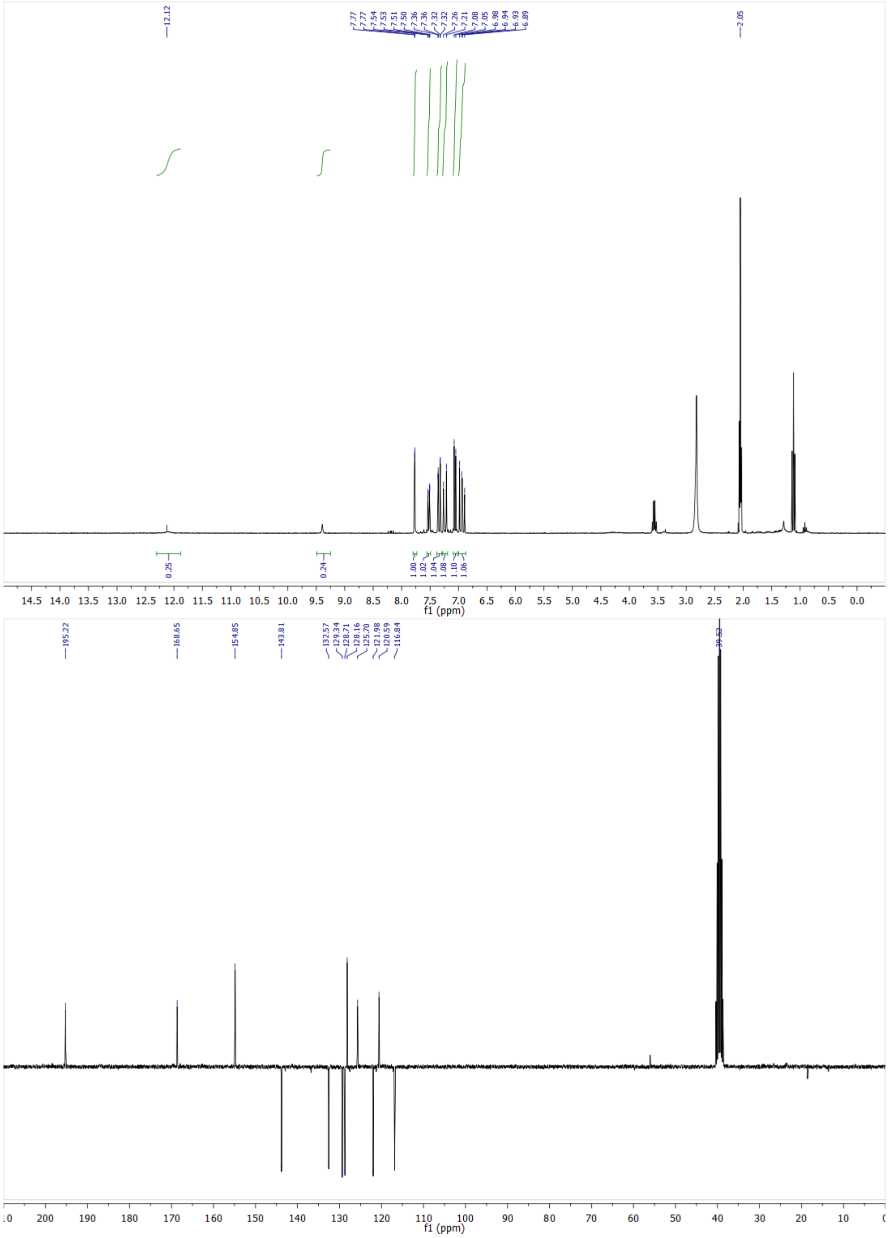

Figure S10: $^{1}$H-NMR and $^{13}$C-NMR spectra of **HPAR3Cl**.

#### 2.2 Production and purification of fluorescent proteins

##### 2.2.1 Plasmids

The plasmids for bacterial expression of **Dronpa-2**[1] and mammalian expression of **H2B-Dronpa-2**[2] were previously described. The plasmids for mamalian expression of H2B-xFAST (x=Y,[3] p,[4] nir,[5] fr,[6] t,[4] green,[7] i,[8] o,[4] Tsi[9] ) have been reported in the original articles.

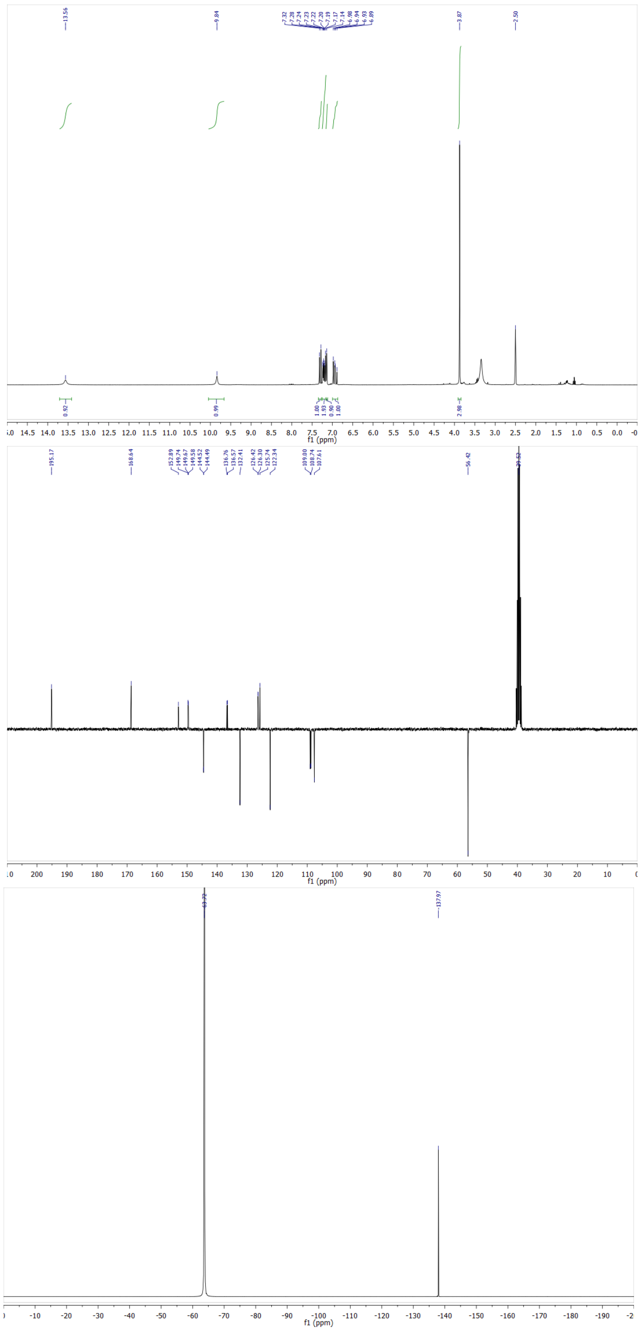

Figure S11: $^{1}$H-NMR, $^{13}$C-NMR, and $^{19}$F-NMR spectra of **HPAR3F5OM**.

##### 2.2.2 Production and purification of Dronpa-2

The **Dronpa-2** plasmid with an N-terminal hexahistidine tag was transformed in E. coli TOP10 strain. Cells were grown in Terrific Broth at 37 °C. The expression was induced at 16 °C by addition of IPTG to a final concentration of 1 mM at optical density at 600 nm of 0.6. The cells were collected after 16 h of expression and lysed by sonication in lysis buffer (50 mM PBS with 150 mM NaCl at pH 7.4, 1 mg.ml$^{-1}$ DNAse, 5 mM $MgCl_2$ and 1 mM phenyl-methylsulfonyl fluoride, and a cocktail of protease inhibitors (Roche, ref 04693132001). After lysis, the mixture was incubated on ice for 2 h for DNA digestion. The insoluble material was removed by centrifugation and the supernatant was incubated overnight with

Ni-NTA agarose beads (ThermoFisher) at 4 °C in a rotator-mixer. The protein loaded Ni-NTA column was washed with 20 column volumes of N1 buffer (50 mM PBS, 150 mM NaCl, 20 mM imidazole, pH 7.4) and 5 column volumes of N2 buffer (50 mM PBS, 150 mM NaCl, 40 mM imidazole, pH 7.4). The bound protein was eluted with N3 buffer (50 mM PBS, 150 mM NaCl, 0,5 M imidazole, pH 7.4). Imidazole was removed from protein fractions using an PD-10 column (Amersham -Cytiva) and replaced with 50 mM PBS, 150 mM NaCl pH 7.4 buffer.

##### 2.2.3 Production and purification of pFAST

**Expression.** Plasmids were transformed in Rosetta (DE3) pLysS E. coli (Merck). Cells were grown at 37 °C in a lysogen broth (LB) medium supplemented with 50 $\mu$g/mL kanamycin and 34 $\mu$g/mL of chloramphenicol to $OD_{600nm}$ 0.6. Expression was induced overnight at 16 °C by adding isopropyl-$\beta$-D-1-thiogalactopyranoside (IPTG) to a final concentration of 1 mM. Cells were collected by centrifugation (4300×g for 20 min at 4 °C) and frozen.
**Purification.** The cell pellet was resuspended in lysis buffer (PBS supplemented with 2.5 mM $MgCl_2$, 1 mM of protease inhibitor phenylmethanesulfonyl fluoride PMSF, and 0.025 mg/mL DNAse, pH 7.4) and sonicated (5 min, 20% of amplitude) on ice. The lysate was incubated for 2 h on ice to allow DNA digestion by DNAse. Cellular fragments were removed by centrifugation (9000×g for 1 h at 4 °C). The supernatant was incubated overnight at 4 °C by gentle agitation with prewashed Ni-NTA agarose beads in PBS buffer complemented with 20 mM of imidazole. Beads were washed with 10 volumes of PBS complemented with 20mM of imidazole and with 5 volumes of PBS complemented with 40 mM of imidazole. His-tagged proteins were eluted with 5 volumes of PBS complemented with 0.5M of imidazole. The buffer was exchanged to PBS (0.05 M phosphate buffer and 0.150 M NaCl) using PD-10 desalting columns or Midi-Trap G-25 (GE Healthcare). The purity of the proteins was evaluated using SDS–PAGE electrophoresis stained with Coomassie blue.

##### 2.2.4 nirFAST expression in bacteria and purification

Plasmid encoding nirFAST fused to a His-tag was transformed in competent *Escherichia coli* expression strain (BL21(DE3) Competent E. coli / ref C2527H New England Biolabs). Cells were grown at 37 °C in lysogen broth medium supplemented with 50 $\mu$g.mL$^{-1}$ kanamycin (and 34 $\mu$g.mL$^{-1}$ of chloramphenicol for Rosetta) to OD(600 nm) 0.6. Expression was induced overnight at 16 °C by adding isopropyl $\beta$-D-1-thiogalactopyranoside (IPTG) to a final concentration of 1 mM. Cells were collected by centrifugation (4,300 × g for 20 min at 4 °C) and frozen. For purification, the cell pellet was resuspended in lysis buffer (PBS supplemented with 2.5 mM $MgCl_2$, 1 mM of protease inhibitor phenylmethanesulfonyl fluoride and 0.025 mg.mL$^{-1}$ DNase, pH 7.4) and sonicated (5 min, 20 % of amplitude) on ice. The lysate was incubated for 2 h on ice to allow DNA digestion by DNase. Cellular fragments were removed by centrifugation (9000×g for 1 h at 4 °C). The supernatant was incubated overnight at 4 °C by gentle agitation with pre-washed Ni-NTA agarose beads in PBS buffer complemented with 20 mM of imidazole. Beads were washed with ten volumes of PBS complemented with 20 mM of imidazole and with five volumes of PBS complemented with 40 mM of imidazole. His-tagged proteins were eluted with five volumes of PBS complemented with 0.5 M of imidazole. The buffer was exchanged to PBS (0.05 M phosphate buffer and 0.15 M NaCl) using PD-10 desalting columns or Midi-Trap G-25 (GE Healthcare). The purity of the proteins was evaluated using SDS–PAGE electrophoresis stained with Coomassie blue.

##### 2.2.5 Preparation and storage of the solutions

The stock solutions of the free fluorogens were first prepared at 10 mM concentration in spectroscopy grade dimethyl sulfoxide (DMSO) and stored in a glass vial covered with aluminum foil at – 20°C.
The **pFAST** and **nirFAST** stock solutions in 1x pH = 7.4 PBS buffer (140 mM NaCl, 10 mM $Na_2HPO_4$) were also stored at – 20°C. They were subsequently diluted to the final concentrations in 1x pH = 7.4 PBS buffer just prior to use. The resulting solutions have been covered with aluminum foil and preserved in the dark for the duration of experiment.
All fluorogens and **pFAST** solutions were prepared always in a glass vial to avoid non-specific adsorption.

#### 2.3 Production of labeled mammalian cells

HeLa cells (ATCC CCL2) were cultured in Dulbecco's Modified Eagle Medium (DMEM High glucose, Hyclone, Cytiva) supplemented with phenol red and 10% (vol/vol) fetal calf serum at 37°C in a 5% $CO_2$ atmosphere. The cells were then seeded ($1.4 \times 10^5$ cells/mL) in $\mu$-Dish 35 mm Imaging Chamber IBIDI (Biovalley, Clinisciences) coated with poly-L-lysine (ref P4832-Sigma-Merck). Cells were transiently transfected using Genejuice (Merck) according to the manufacturer's protocols for 24–48 h prior to imaging. For imaging, live cells were washed with DPBS (Dulbecco's Phosphate-Buffered Saline), and treated with DMEM media (without serum and phenol red) containing the fluorogens at the indicated concentration. The cells were imaged directly without washing. Fixation of cells was performed using formaldehyde solution at 3.7% for 25 min. Fixed cells were then washed three times with DPBS and treated with DPBS containing the fluorogens at the indicated concentration prior to imaging.

#### 2.4 Instruments

##### 2.4.1 UV/Vis absorption and fluorescence spectrometers

UV/Vis absorption spectra were recorded on a UV/Vis spectrophotometer (Cary 300 UV-Vis, Agilent Technologies, Santa Clara, CA) at 293 K equipped with a Peltier 1×1 thermostatic cell holder (Agilent Technologies). Samples were contained either in 1 cm × 1 cm (3 mL; the cuvette content was stirred) or in 0.3 cm × 0.3 cm (54 µL; the cuvette content was not stirred) quartz cuvettes (Hellma Optics, Jena, Germany). Fluorescence measurements were acquired on a LPS 220 spectrofluorometer (PTI, Monmouth Junction, NJ), equipped with a TLC50 cuvette holder (Quantum Northwest, Liberty Lake, WA) thermoregulated at 293 K.

##### 2.4.2 Light sources

In order to photoconvert the reactants and track the evolution of fluorescent products via absorbance and fluorescence under time constant illumination, we used: a (i) 365, (ii) 405 or (iii) 480 nm Light Emitting Diode (NCSU033B, Nichia Corp., Anan, Japan; LHUV-405, LXZ1-PB01, Lumileds, NL) monochromatically filtered (BP ZET 365-20, Chroma technology, Bellows Falls, VT, BP ET405/20x; Chroma Technology, VT, BP FF01-479/40-25, Semrock, NY) to generate a spatially homogeneous source of illumination.

##### 2.4.3 pH measurements

By assimilating activity and concentration, the proton concentration was directly measured after calibration of the pH meter (PHM210 standard pH meter from MeterLab, Radiometer Analytical) equipped with a combined pH electrode (BNC plug).

##### 2.4.4 Confocal microscopes

The confocal micrographs of the cells were acquired either on a Zeiss LSM 710 Laser Scanning Microscope equipped with continuous laser lines at 405 and 488 nm and a Plan NeoFluar 20×/0.5 objective, or on a Zeiss LSM 980 confocal Laser Scanning Microscope equipped with a 63 ×/1.4 NA oil immersion objective, continuous laser lines delivering light at 405, 488, 561 and 639 nm and equipped with photomultiplier modules. ZEN black software was used to collect the data. The images were analyzed with Fiji (Image J) or ZEN blue.

##### 2.4.5 Home-built microscope for patterned illumination

The images of Figures 4c,d of the Main Text have been recorded with a highly versatile dual–digital micromirror device (DMD) microscope that enables independent and simultaneous spatial and temporal modulation of illumination at two wavelengths, accommodates a wide range of spatial scales through objective interchangeability, and supports diverse excitation and emission wavelength combinations.[10]

#### 2.5 Methods

##### 2.5.1 Measurement of light intensity

The light intensities were measured either through exploiting fluorescent actinometers[11] or by using a power meter.
In this manuscript, we provide the values of the light intensities in E.m$^{-2}$.s$^{-1}$ (or mol. of photons.m$^{-2}$.s$^{-1}$). This unit is currently used in actinometry. However, it is not often used in other fields such as optical microscopy, in which the researchers prefer to adopt W.m$^{-2}$. We provide below the conversion between both units.
We consider a monochromatic light of wavelength $\lambda_{\rm exc}$. Its values in E.m$^{-2}$.s$^{-1}$ and W.m$^{-2}$ are respectively denoted as $I(\lambda_{\rm exc}, {\rm E.m^{-2}.s^{-1}})$ and $I(\lambda_{\rm exc}, {\rm W.m^{-2}})$. The relation between $I(\lambda_{\rm exc}, {\rm E.m^{-2}.s^{-1}})$ and $I(\lambda_{\rm exc}, {\rm W.m^{-2}})$ is given in Eq.(S43)

$$I(\lambda_{\rm exc}, {\rm W.m^{-2}}) \quad = \quad \frac{hcN_A}{\lambda_{\rm exc}} \times I(\lambda_{\rm exc}, {\rm E.m^{-2}.s^{-1}}) \approx 0.12 \times \frac{{\rm I}(\lambda_{\rm exc}, {\rm E.m^{-2}.s^{-1}})}{\lambda_{\rm exc}\,({\rm m})} \qquad \text{(S43)}$$

with the Planck constant $h = 6.63\ 10^{-34}$ m$^2$.kg.s$^{-1}$, speed of light in a vacuum $c = 3.00\ 10^8$ m.s$^{-1}$, the Avogadro number $N_A = 6.02\ 10^{23}$ mol$^{-1}$, and where $\lambda_{\rm exc}$ is in m.

###### 2.5.1.1 Cuvette experiments

The light intensity delivered from the light sources (i) reported in subsection 2.4.2 has been calibrated by using $\alpha$-(4-diethylamino)phenyl)-N-phenylnitrone as an actinometer and rhodamine B as a fluorescent reporter. The light intensity delivered from light sources (ii) and (iii) reported in subsection 2.4.2 has been calibrated by using 1 $\mu$M Dronpa-2 solution in 1x pH = 7.4 PBS buffer.[11]

###### 2.5.1.2 Confocal microscopes

The light intensity applied at 405 and 488 nm laser in the cell experiments was measured from analyzing the time evolution of the Dronpa-2 fluorescence at the nucleus of Dronpa-2-labeled fixed Hela cells recorded at different light powers over area of 40 × 40 pixels$^2$ (frame: 512 × 512 pixel$^2$; zoom factor: 1; dwell time: 1.02 $\mu$s).[11]
The light intensity applied at 445, 514, 543, 561, and 639 nm with the lasers during the cell experiments in confocal microscopy was measured by analyzing the fluorescence intensity of a 10 $\mu$M DDAO solution in aqueous HEPES buffer (pH 7.9; 100 mM NaCl, 5 mM NaOH, 10 mM HEPES), recorded at different light powers over an area of 40 × 40 pixels$^2$ (frame: 512 × 512 pixels$^2$; zoom factor: 4; dwell time: 0.49 $\mu$s), following the protocols described in a previous report.[11]

##### 2.5.2 Calibration of the detector sensitivity in confocal microscopy

In order to retrieve the hue of the fluorescence emission, it is first necessary to measure the relative sensitivities of the detection channels, which depend on the detector type, the spectral range and the voltage gain. In this work, we used the GaAsP (for the visible range) and multialkali (for NIR) photomultipliers available on the LSM 980 microscope. For calibration, we imaged live **H2B-pFAST** HeLa cells in DMEM buffer in the presence of 1 $\mu$M **HBR3F5OM** under 488 nm illumination. This fluorogen was chosen as a convenient reference since its emission covers the 500-750 nm spectral region of interest.
First, the dependence of sensitivity on wavelength was investigated by imaging the same cell with both detector types and varying the channel spectral range, the width of which was kept at 50 nm, from 500-550 to 700-750 nm. We verified that under these conditions, photobleaching was negligible at the end of the sequence. An ROI in the labeled zone was chosen and its average signal intensity $S(\lambda)$ measured, where $\lambda$ is the center of the detection spectral range. For each detector type, we define an approximate relative wavelength sensitivity $s_\lambda(\lambda)$:

$$s_\lambda(\lambda) \approx \frac{S(\lambda)}{S(\lambda_{ref})}\frac{b(\lambda_{ref})}{b(\lambda)} \qquad \text{(S44)}$$
$$\lambda_{ref} = 500\ {\rm nm}$$

where $b(\lambda)$ is the brightness of the fluorescent complex. Since $S(\lambda)$ is not continuous, linear interpolation and extrapolation was needed to extend its domain to the entire 500-750 nm region. The relative sensitivities of the multialkali (MA) and GaAsP detectors were defined at the same reference wavelength:

$$\begin{aligned} s_{GaAsP} &= 1 \\ s_{MA} &= \frac{S_{MA}(\lambda)}{S_{GaAsP}(\lambda)} \\ \lambda_{ref} &= 500 \text{ nm} \end{aligned} \tag{S45}$$

Second, the dependence of sensitivity on gain voltage was investigated by imaging the same cell with both detector types and varying the voltage $U_g$ from 650 to 800 V in 50 V increments. The detection range was set to 500-700 nm for both detector types. As before, the ROI average signal $S(U_g)$ was measured. For each detector type, we define a gain sensitivity $s_g(U_g)$:

$$\begin{aligned} s_g(U_g) &= \frac{S(U_g)}{S(U_{g,ref})} \\ U_{g,ref} &= 650\ V \end{aligned} \tag{S46}$$

Finally, we can define the sensitivity $s(\lambda)$ for an arbitrary detection channel with detection range $\lambda_1 - \lambda_2$, gain voltage $U_g$ and detector type $det$:

$$s(\lambda) = \begin{cases} s_{g,det}(U_g) s_{det} s_{\lambda,det}(\lambda) & \text{if } \lambda \in [\lambda_1, \lambda_2] \\ 0 & \text{otherwise} \end{cases} \tag{S47}$$

###### 2.5.3 Methods for acquisition of the thermokinetic and photophysical parameters of the free fluorogens

**2.5.3.1 Measurement of $K_{\mathbf{FH}}$** The present fluorogens contain an ionizable phenol group yielding an acid and a basic state denoted **FH** and **F**$^-$. At chemical equilibrium, one can write Eqs.(S48–S49)

$$FH = \frac{H}{H + K_{\mathbf{FH}}} F_{tot} \tag{S48}$$

$$F^- = \frac{K_{\mathbf{FH}}}{H + K_{\mathbf{FH}}} F_{tot} \tag{S49}$$

where $H$ designates the proton concentration and $K_{\mathbf{FH}}$ is the proton exchange thermodynamic constant associated to **FH**. In the present study, we retrieved $K_{\mathbf{FH}}$ by analyzing the evolutions of the absorbance $A(\lambda)$ of the free fluorogens as a function of pH were analyzed with Eq.(S50)

$$A(\lambda) = (\varepsilon_{F^-} F^- + \varepsilon_{FH} FH)\ell \tag{S50}$$

where $\varepsilon_{F-}$, $\varepsilon_{FH}$, and $\ell$ respectively designate the molar absorption coefficients of **F**$^-$ and **FH**, and the optical path length, and where $F^-$ and $FH$ are given in Eqs.(S48,S49).

**2.5.3.2 Measurement of $k^{\Delta}_{FF'} + k^{\Delta}_{F'F}$** The sum of the thermal rate constants $k^{\Delta}_{FF'} + k^{\Delta}_{F'F}$ associated with thermal relaxation of the fluorogen after photoisomerization was determined by performing a thermal return experiment. A solution of free fluorogen was illuminated with light intensity $I$ until it reached the steady state. Then the illumination was turned off for a period of time $\Delta t$, before illuminating the sample again up to its steady state. This sequence was repeated for a range of $\Delta t$ values. The recovery of fluorescence $\Delta I^{\Delta}_F$ during the dark period as a function of $\Delta t$ is given in Eq.(S51).

$$\Delta I^{\Delta}_F(\Delta t) = I_{F,\infty} + \Delta I_F e^{-(k^{\Delta}_{FF'} + k^{\Delta}_{F'F})\Delta t} \tag{S51}$$

Fitting the experimental data with Eq.(S51) yielded $k^{\Delta}_{FF'} + k^{\Delta}_{F'F}$.

**2.5.3.3 Measurement of $\sigma_{FF'} + \sigma_{F'F}$** The effective cross section associated with photoisomerization of the free fluorogen, $\sigma_{FF'} + \sigma_{F'F}$, was determined by fitting the time evolution of fluorescence intensity of a solution of free fluorogen under illumination at constant light intensity $I$ as reported in reference.[12] The dependence of the inverse of the characteristic time $\tau$ retrieved at different $I$ values was then linearly fitted with Eq.(S52).

$$\frac{1}{\tau(I)} = k^{\Delta}_{F'F} + k^{\Delta}_{FF'} + I(\sigma_{FF'} + \sigma_{F'F}) \tag{S52}$$

$\sigma_{FF'} + \sigma_{F'F}$ was eventually determined from the slope of the fit.

##### 2.5.4 Methods for acquisition of the thermokinetic and photophysical parameters of the complexes between pFAST or nirFAST and the fluorogens

**2.5.4.1 Measurement of the quantum yield of fluorescence emission** The overall emission quantum yield after one-photon excitation, $\Phi$, was calculated from Eq. (S5)

$$\Phi_F(\lambda_{exc}) = \Phi_R \times \frac{I_S}{A_S} \times \frac{A_R}{I_R} \times \frac{n_S^2}{n_R^2} \tag{S5}$$

where the subscripts S and R stand for samples and standard reference respectively, $A$ is the absorbance at the excitation wavelength $\lambda_{exc}$, $I$ is the integrated area of the emission spectrum, and $n$ is the refractive index of the solvent. The standard reference for the quantum yield measurements was either rhodamine 6G in ethanol or fluorescein-sodium salt in 0.1 M NaOH with $\Phi_{ref}$ = 0.95 (rhodamine 6G) and 0.98 (fluorescein-sodium salt). The absorption and emission spectra were recorded at five different concentrations of fluorogen in presence of excess **pFAST** or **nirFAST** and 3 different concentrations of standard fluorophore. To record emission spectra, $\lambda_{exc}$ was kept fixed for both the fluorogen and standard reference. The slopes were calculated from integrated area vs absorbance plot for the fluorogen and standard reference and used to calculate the fluorescence quantum yield, $\Phi$ ($\lambda_{exc}$).

**2.5.4.2 Measurement of the thermodynamic dissociation constant of the complex between pFAST or nirFAST and the thermodynamically stable state of the fluorogen ($K_d$)** The thermodynamic dissociation constant of the complex between **pFAST** and the thermodynamically stable state of the fluorogen $K_d$ was determined by fitting the dependence of the fluorescence intensity $I_F$ on the total concentration of the fluorogen $F_{tot}$ at constant concentration of the protein $P_{tot}$ by using Eq.(S53), which was derived from Eq.(S1,S10). Since the complex between **pFAST** and the thermodynamically stable state of the fluorogen is photoactive, this dependence has been retrieved from extrapolating to time zero the decay of the fluorescence signal upon constant illumination.

$$I_F = IQ_B \frac{(F_{tot} + P_{tot} + K_d) - \sqrt{(F_{tot} + P_{tot} + K_d)^2 - 4F_{tot}P_{tot}}}{2} \tag{S53}$$

where $IQ_B$ is a constant scaling factor.

**2.5.4.3 Measurement of $k_{on}, k_{off}$** The rate constants associated with the formation of the complex between **pFAST** and the thermodynamically stable (*Z*) stereoisomer of the fluorogen can be measured using a stop-flow experiment by rapidly mixing the fluorogen and protein solutions and following the fluorescence signal over time.
In the investigated series of fluorogens, we only observed the (*Z*) stereoisomer at chemical equilibrium. Hence, $K_F \approx 0$. In order to eliminate the impact of photoisomerization, this experiment has further been performed at low light intensity and high enough concentrations so that the time evolution of the fluorescence signal at short time is governed by the binding step involving the , which is much faster than the photoisomerization step ($k_{on}F_{tot} + k_{off} \gg k^{\Delta}_{FF'} + k^{\Delta}_{F'F} + I(\sigma_{FF'} + \sigma_{F'F}), k^{\Delta}_{BB'} + k^{\Delta}_{B'B} + I(\sigma_{BB'} + \sigma_{B'B})$). Then, the system obeys the two-state model shown in Figure (S12).

$$\mathbf{F+P} \underset{k_{off}}{\overset{k_{on}}{\rightleftharpoons}} \mathbf{B}$$

Figure S12: Kinetic model relevant under the experimental conditions of our stop-flow binding experiment.

We consider that the two feeding syringes respectively contain **F** and **P** such that their respective initial concentrations in the cuvette after fast mixing are $F_{tot}$ and $P_{tot}$. The instantaneous concentrations of **F**, **P**, and **B** in the cuvette, $F$, $P$, and $B$, monotonously evolve towards the equilibrium concentrations $F(\infty)$, $P(\infty)$, and $B(\infty)$ which are given by the expressions (S1–S3). The time evolution of the concentrations is driven by the system of differential equations (S54,S55),

$$\dot{P} = -k_{on}FP + k_{off}B \tag{S54}$$
$$\dot{B} = k_{on}FP - k_{off}B \tag{S55}$$

with solution (S56–S59).[13]

$$B(t) = B(\infty)\left\{1 - \frac{e^{-\frac{t}{\tau}}}{1 + k_{on}\tau B(\infty)\left[1 - e^{-\frac{t}{\tau}}\right]}\right\} \tag{S56}$$
$$F(t) = F_{tot} - B(t) \tag{S57}$$
$$P(t) = P_{tot} - B(t) \tag{S58}$$
$$\tau = \frac{1}{k_{on}(F_{tot} + P_{tot}) + k_{off}} \tag{S59}$$

and time evolution of the fluorescence intensity of this system governed by Eq.(S60)

$$I_F(t) = Q_B I B(\infty)\left\{1 - \frac{e^{-\frac{t}{\tau}}}{1 + k_{on}\tau B(\infty)\left[1 - e^{-\frac{t}{\tau}}\right]}\right\} \tag{S60}$$

by neglecting the brightness of the free fluorogen.
with

$$\frac{1}{\tau(F_{tot})} = k_{on}F_{tot} + k_{off} \tag{S61}$$
$$I_{F,\infty} = F_{tot}I(Q_F + \frac{Q_B k_{on} P_{tot}}{k_{on}F_{tot} + k_{off}}) \tag{S62}$$
$$\Delta I_F = I F_{tot} Q_F - I_{F,\infty} \tag{S63}$$

The time evolution of the fluorescence signal at different $F_{tot}$ values is processed to retrieve the characteristic time $\tau(F_{tot})$ with Eq.(S60). $k_{on}$ is then retrieved from the slope of the dependence of $1/\tau$ on $F_{tot}$ with Eq.(S59). Once $k_{on}$ is extracted, $k_{off}$ is computed by using Eq.(S64)

$$k_{off} = K_d k_{on} \tag{S64}$$

**2.5.4.4 Measurement of $k^{\Delta}_{BB'} + k^{\Delta}_{B'B}$** The sum of the thermal rate constants $k^{\Delta}_{BB'} + k^{\Delta}_{B'B}$ was determined by performing a thermal return experiment in the presence of protein in excess such that $P_{tot} \gg K'_d \gg K_d$. The solution was illuminated with light intensity $I$ until it reached the steady state. Then the illumination was turned off for a period of time $\Delta t$, before illuminating the sample again up to its steady state. This sequence was repeated for a range of $\Delta t$ values. The fluorescence recovery associated with the thermal return follows a mono-exponential decay given in Eq.(S65).

$$\Delta I^{\Delta}_F(\Delta t) = I_{F,\infty} + \Delta I_F e^{-\Delta t/\tau} \tag{S65}$$

with

$$\frac{1}{\tau} = \langle k_+ \rangle + \langle k_- \rangle \tag{S66}$$

$$\langle k_+ \rangle = \frac{K_d k^{\Delta}_{FF'} + P_{tot} k^{\Delta}_{BB'}}{K_d + P_{tot}} \approx k^{\Delta}_{BB'} \tag{S67}$$

$$\langle k_- \rangle = \frac{K'_d k^{\Delta}_{F'F} + P_{tot} k^{\Delta}_{B'B}}{K'_d + P_{tot}} \approx k^{\Delta}_{B'B} \tag{S68}$$

Fitting the data with Eq.(S65) yields $\tau$ and from it, $k^{\Delta}_{BB'} + k^{\Delta}_{B'B}$.
Note that in practice $k^{\Delta}_{BB'} + k^{\Delta}_{B'B}$ was found to be extremely slow ($< 0.0001 s^{-1}$) for all the studied fluorogens.

**2.5.4.5 Measurement of $k^{\Delta}_{BB'}$ and $k^{\Delta}_{B'B}$** At large total concentration in protein scaffold such that $P_{tot} \gg K'_d \gg K_d$, a measurement of the $B'/B$ ratio can be achieved by NMR spectroscopy in order to determine $k^{\Delta}_{BB'}$ and $k^{\Delta}_{B'B}$. Indeed

$$F(\infty) \approx 0 \tag{S69}$$

$$F'(\infty) \approx 0 \tag{S70}$$

$$B(\infty) = \frac{F_{tot}}{K^{\Delta}_B + 1} \tag{S71}$$

$$B'(\infty) = \frac{F_{tot} K^{\Delta}_B}{K^{\Delta}_B + 1} \tag{S72}$$

which yields

$$\frac{B'(\infty)}{B(\infty)} = K^{\Delta}_B = \frac{k^{\Delta}_{BB'}}{k^{\Delta}_{B'B}} \tag{S73}$$

Since $K^{\Delta}_B$ and $k^{\Delta}_{BB'} + k^{\Delta}_{B'B}$ are known, $k^{\Delta}_{BB'}$ and $k^{\Delta}_{B'B}$ can be calculated as

$$k^{\Delta}_{BB'} = \frac{(k^{\Delta}_{BB'} + k^{\Delta}_{B'B}) K^{\Delta}_B}{1 + K^{\Delta}_B} \tag{S74}$$

$$k^{\Delta}_{B'B} = \frac{k^{\Delta}_{BB'} + k^{\Delta}_{B'B}}{1 + K^{\Delta}_B} \tag{S75}$$

Note that $k^{\Delta}_{BB'}$ was found to be negligible for all studied fluorogens.

**2.5.4.6 Measurement of $\sigma_{BB'} + \sigma_{B'B}$** $\sigma_{BB'} + \sigma_{B'B}$ was measured by analyzing the time evolution of the fluorescence signal from a solution containing both the protein scaffold and the fluorogen in the presence of an excess of protein so that $P_{tot} \gg K'_d \gg K_d$ and under experimental conditions when fluorogen photoisomerization is rate limiting. Under such conditions,

$$\langle k_+ \rangle = \frac{K_d (k^{\Delta}_{FF'} + I\sigma_{FF'}) + P_{tot} (k^{\Delta}_{BB'} + I\sigma_{BB'})}{K_d + P_{tot}} \approx k^{\Delta}_{BB'} + I\sigma_{BB'} \tag{S76}$$

$$\langle k_- \rangle = \frac{K'_d (k^{\Delta}_{F'F} + I\sigma_{F'F}) + P_{tot} (k^{\Delta}_{B'B} + I\sigma_{B'B})}{K'_d + P_{tot}} \approx k^{\Delta}_{B'B} + I\sigma_{B'B} \tag{S77}$$

the time evolution of the fluorescence intensity follows a mono-exponential decay given in Eq.(S78)

$$I_F(t) = I_{F,\infty} + \Delta I_F e^{-t/\tau} \tag{S78}$$

with

$$\frac{1}{\tau(I)} = k^{\Delta}_{BB'} + k^{\Delta}_{B'B} + I(\sigma_{BB'} + \sigma_{B'B}). \tag{S79}$$

The characteristic time $\tau$ was retrieved from fitting the time evolution of the fluorescence intensity with Eq.(S78) for a range of light intensities $I$. $\sigma_{BB'} + \sigma_{B'B}$ was then extracted from the slope of a linear fit of $1/\tau$ against $I$.

**2.5.4.7 Measurement of $Q'_B/Q_{B'}, \sigma_{BB'}$, and $\sigma_{B'B}$** Direct measurement of the concentrations of the proportions of the states **B** and **B'** by NMR spectroscopy under illumination to retrieve $\sigma_{BB'}/\sigma_{B'B}$ as reported above proved to be difficult. Therefore, an indirect approach leveraging knowledge on the photochemical properties of the free fluorogen was used.

This approach involves starting with a mixture of the fluorogen states **F** and **F'** at total concentration $F_{tot}$ with a known $F'/F$ ratio. This ratio can be controlled by applying a calibrated jump of light intensity right before the next step. Typically either no (**a**) or high (**b**) light intensity is used, fixing the $F'/F$ ratio to $K_F^\Delta$ or $\Sigma_F$ respectively. Then an excess of protein such that $P_{tot} > F_{tot}$ and $P_{tot} \gg K'_d > K_d$ is rapidly added and mixed. It results in an initial state with the concentrations of **B** and **B'** in the same ratio as the ones of **F** and **F'**.[b]

Upon illumination at light intensity $I$, the fluorescence signal evolves as per the model in Eq.(S78). From the fit, we extracted the initial fluorescence intensities with expressions given in Eqs. (S80,S81)

$$I_{F,a}(0) = \frac{F_{tot}I(Q_B + Q_{B'}K_F)}{1+K_F} \tag{S80}$$

$$I_{F,b}(0) = \frac{F_{tot}I(Q_B + Q_{B'}\Sigma_F)}{1+\Sigma_F} \tag{S81}$$

Then, we could compute the ratio of brightnesses of the complexes $y = Q'_B/Q_{B'}$ by using Eq.(S82)

$$\frac{Q_{B'}}{Q_B} = y = \frac{1+K_F - x(1+\Sigma_F)}{xK_F(1+\Sigma_F) - \Sigma_F(1+K_F)} \tag{S82}$$

with

$$x = \frac{I_{F,b}(0)}{I_{F,a}(0)} \tag{S83}$$

In both cases (**a**) and (**b**), the fluorescence intensity tends towards $I_F(\infty)$ at steady state given in Eq.(S84), which is governed by the light-driven kinetics.

$$I_F(\infty) = \frac{F_{tot}I(1+y\Sigma_B)}{1+\Sigma_B} \tag{S84}$$

Then, $\Sigma_B$ can be calculated using Eq.(S85).

$$\Sigma_B = \frac{1-z+K_F(1-yz)}{z(1+yK_F) - y(1+K_F)} \tag{S85}$$

with

$$z = \frac{I_F(\infty)}{I_{F,a}(0)} \tag{S86}$$

Finally, $\sigma_{BB'}$ and $\sigma_{B'B}$ can be determined from $\sigma_{BB'} + \sigma_{B'B}$ and $\Sigma_B$ using Eqs. (S87,S88).

$$\sigma_{BB'} = \frac{(\sigma_{BB'} + \sigma_{B'B})\Sigma_B}{1+\Sigma_B} \tag{S87}$$

$$\sigma_{B'B} = \frac{\sigma_{BB'} + \sigma_{B'B}}{1+\Sigma_B} \tag{S88}$$

**2.5.4.8 Measurement of $K'_d$** **F'** is kinetically unstable. Hence, a direct measurement of $K'_d$ by titration such as reported above is impossible. To overcome this limitation, we used an indirect method exploiting the difference of the rate constants associated with the thermal recovery after photoisomerization for the free and bound fluorogen. The kinetics of thermal return of a solution of photoisomerized fluorogen with protein concentrations such as $P_{tot} \gg F_{tot}$ and $P_{tot} \gg K_d$ was measured as in subsection 2.5.3.2 at different total protein concentrations $P_{tot}$. It was fitted with Eq.(S51) to retrieve $\tau$. $K'_d$ was eventually extracted from fitting the dependence of $1/\tau$ on $P_{tot}$ with Eq.(S89)

$$\frac{1}{\tau(P_{tot})} = k^\Delta_{BB'} + \frac{K'_d k^\Delta_{F'F} + P_{tot}k^\Delta_{B'B}}{K'_d + P_{tot}} \tag{S89}$$

[b]The addition must be much faster than the thermal kinetics $k^\Delta_{FF'} + k^\Delta_{F'F}$ so that the ratio remains unchanged.

##### 2.5.5 Image segmentation and extraction of single-cell kinetic profiles

**2.5.5.1 Primary cell segmentation** Each video contained between one and two cells with labelled nuclei. To segment the images, we selected a reference frame consiting in the absolute difference between the fifth and the first frame of the green channel to capture the pixels showing significant dynamics. We retrieved the combined nucleus probability flow map computed with Cellpose (CellposeModel default settings, version 4.0.7). We applied a threshold (30) on that map and selected the largest object in the binary mask as our labeled nucleus, with an additional safeguard for cases where nuclei occupies a larger area than the surrounding image content. For a few cells with high expression level, this threshold included the cytoplasm in the segmentation. Those cells were adressed manually by changing the threshold to 100. A few remaining cell errors were corrected with a more robust segmentation: fluorescence overintensity spots were suppressed using percentile-based masking, and nuclei with low contrast relative to cytoplasmic autofluorescence were enhanced on the average or maximum intensity projections. Cellpose standard model is trained on cells with a mean diameter of 30 pixels, to account for large nucleus size and shape variability, increased downscaling and relaxed flow thresholds were applied.

**2.5.5.2 Sub-segmentation of cells into spatial micro-regions** We collected between 10 and 15 video for each of the FAST variants. To artificially augment our dataset, we subdivided each individual mask into multiple micro-regions. A regular grid of seed points was placed within the cell boundary (mesh spacing: 2.6 $\mu m$ - 20/256 pixels), and a watershed transform was applied under the constraint of the cell mask. We removed from the dataset samples with an area below 10 pixels to limit noise-dominated measurements. We obtained a total dataset of 2335 micro-regions for the labeling proteins.

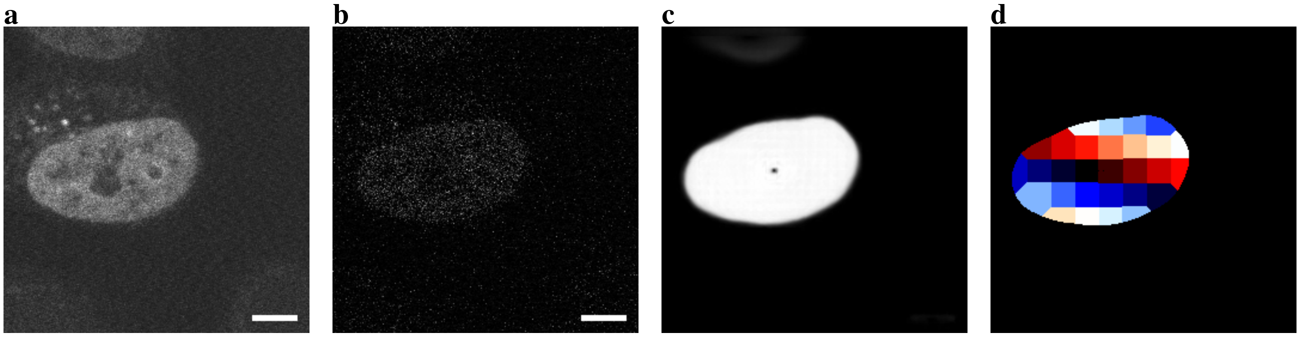

Figure S13: **Segmentation and sub-segmentation workflow.** **a**: Mean of fluorescence frames over the movie; **b**: Reference image (absolute difference between frame 5 and frame 1) used for segmentation; **c**: Cellpose probability map; **d**: Final micro-region partition used for data extraction. Scale bar: 5 $\mu$m.

**2.5.5.3 Extraction and normalization of fluorescence kinetics** For every micro-region, the mean fluorescence intensity was extracted as a function of time for the red and green channels. Each time-series was normalized to its first time point. Normalized traces from the two channels were then concatenated to form a single kinetic trace for each micro-region as displayed in Figure 7 of the Main Text. Each time trace contains 150 datapoints.

**2.5.5.4 Train–test partitioning based on whole cells** To prevent information leakage, the train-test split was performed at the level of whole cells and not individual micro-regions. This ensures that the classifier is evaluated on fluorescence trajectories from entirely unseen cells rather than on neighbouring regions of the same object. We selected 3 cells per protein for the test set and the remaining cells were used for the training. All training procedures were performed with fixed random seeds to enforce reproducibility.

**2.5.5.5 Cross-validation and model selection** We tested several classification models (Linear Discriminant Analysis, Gaussian Naive Bayes and Random Forest) with cross-validation (15-folds). The training set was split into training and validation subsets at the cell level, with 3 cells kept for validation. Since the time traces are relatively long (150 points) compared to the dataset size (1850 samples), we applied a dimensionality reduction to the sub-training set using Principal Component Analysis (PCA). The number of components was treated as a hyperparameter and optimized during cross-validation. The best performing model (Linear Discriminant Analysis) and number of PCA components (8) were selected based on average and standard deviation of cross-validated balanced accuracy.

**2.5.5.6 Final model training and evaluation** PCA (8 components) was fitted on the training set (all training cells and their micro-regions) and the transform was applied to the test set (3 cells per protein). The LDA classifier was trained on the full training dataset. Model performance was assessed on the unseen test set using balanced accuracy and confusion matrices. Micro-region wise, we obtain an accuracy score of 98% with 12 missclassifications out of the 485 micro-region samples. Further, with a voting strategy per cell, we can reach 100% accuracy to correctly classify the 18 test cells.

### 3 Investigation of the free fluorogens

#### 3.1 Absorption and emission spectra of the free fluorogens

Figure S14a–f displays the absorption and emission spectra of the free fluorogens in their thermodynamically stable ($Z$) configuration. Table S1 sums up the results.

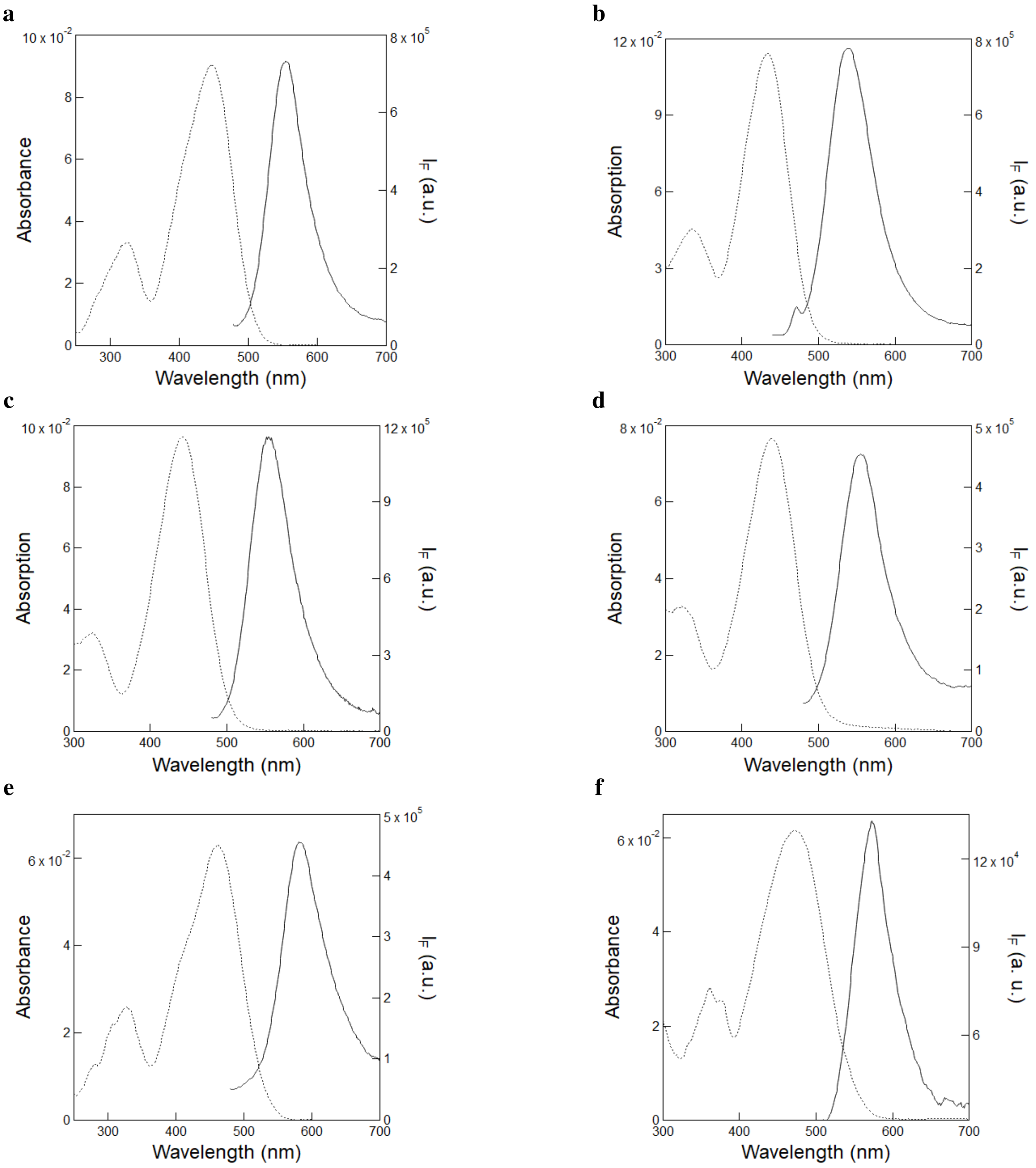

Figure S14: *UV-Vis absorption and emission spectra of the free fluorogens*. The absorption (dotted line) and emission (solid line; $\lambda_{exc}$ = 405 nm) spectra of free fluorogens **HBR3Cl (a)**, **HBR3CN (b)**, **HBR3Cl5F (c)**, **HBR35DF5 (d)**, **HBR3F5OM (e)** and **HBIR3Cl (f)**. The spectra were recorded using 10 $\mu$M fluorogen solution in 1× pH = 7.4 PBS buffer contained in 54 $\mu$L quartz cuvette (3 mm optical pathlength). $T$ = 293 K.

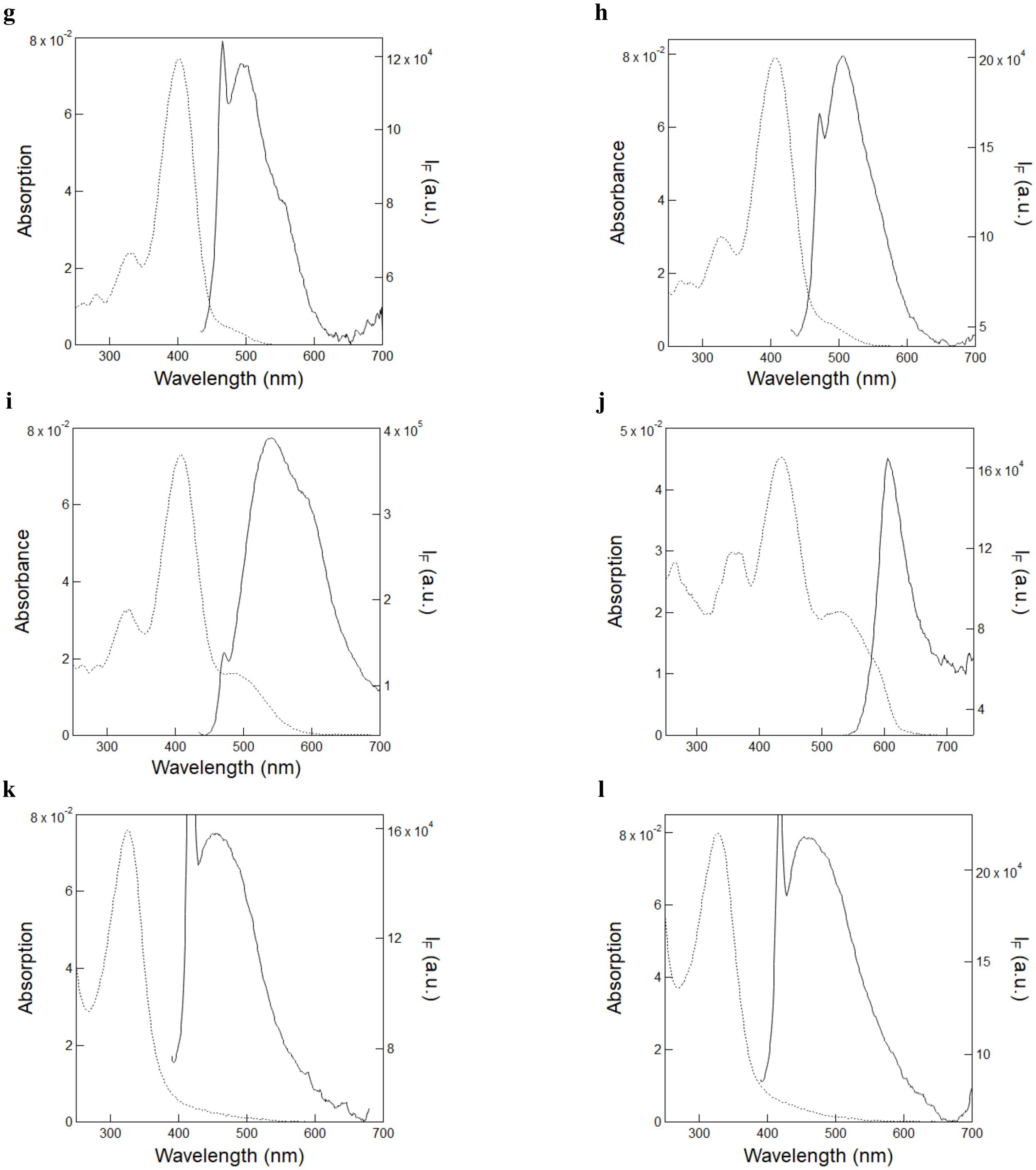

Figure S14: *UV-Vis absorption and emission spectra of the free fluorogens*. The absorption (dotted line) and emission (solid line; $\lambda_{exc}$ = 365 nm (**HBO3M** and **HBO35DM**); $\lambda_{exc}$ = 405 nm (**HBR3M**, **HBR25DM** and **HBR35DOM**); $\lambda_{exc}$ = 480 nm (**HBIR 35DOM**)) spectra of free fluorogens **HBR3M (g)**, **HBR25DM (h)**, **HBR35DOM (i)**, **HBIR35DOM (j)**, **HBO3M (k)** and **HBO35DM (l)**. The spectra were recorded using 10 $\mu$M **HBR3M**, **HBR25DM**, **HBR35DOM**, **HBIR35DOM**, 20 $\mu$M **HBO3M** and 50 $\mu$M **HBO35DM** solution in 1$\times$ pH = 7.4 PBS buffer contained in 54 $\mu$L quartz cuvette (3 mm optical pathlength). $T = 293$ K.

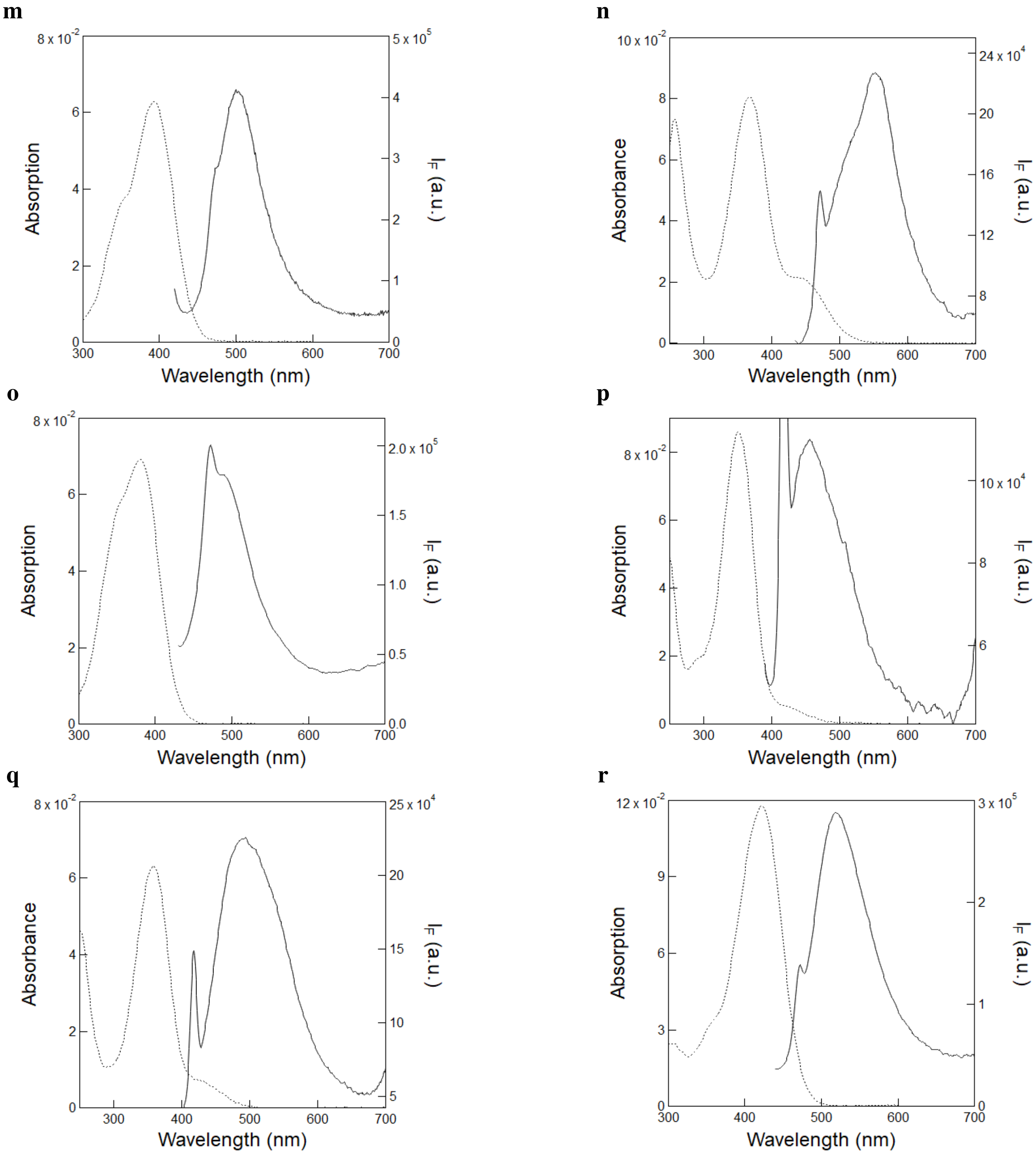

Figure S14: *UV-Vis absorption and emission spectra of the free fluorogens*. The absorption (dotted line) and emission (solid line; $\lambda_{exc}$ = 365 nm (**HBP35DOM**, **HBT35DM** and **HBT35DOM**); $\lambda_{exc}$ = 405 nm (**HBP3CN**, **HBT3CN** and **HBTH3CN**)) spectra of free fluorogens **HBP3CN (m)**, **HBP35DOM (n)**, **HBT3CN (o)**, **HBT35DM (p)**, **HBT35DOM (q)** and **HBTH3CN (r)**. The spectra were recorded using 10 $\mu$M fluorogen solution in $1\times$ pH = 7.4 PBS buffer contained in 54 $\mu$L quartz cuvette (3 mm optical pathlength). $T = 293$ K.

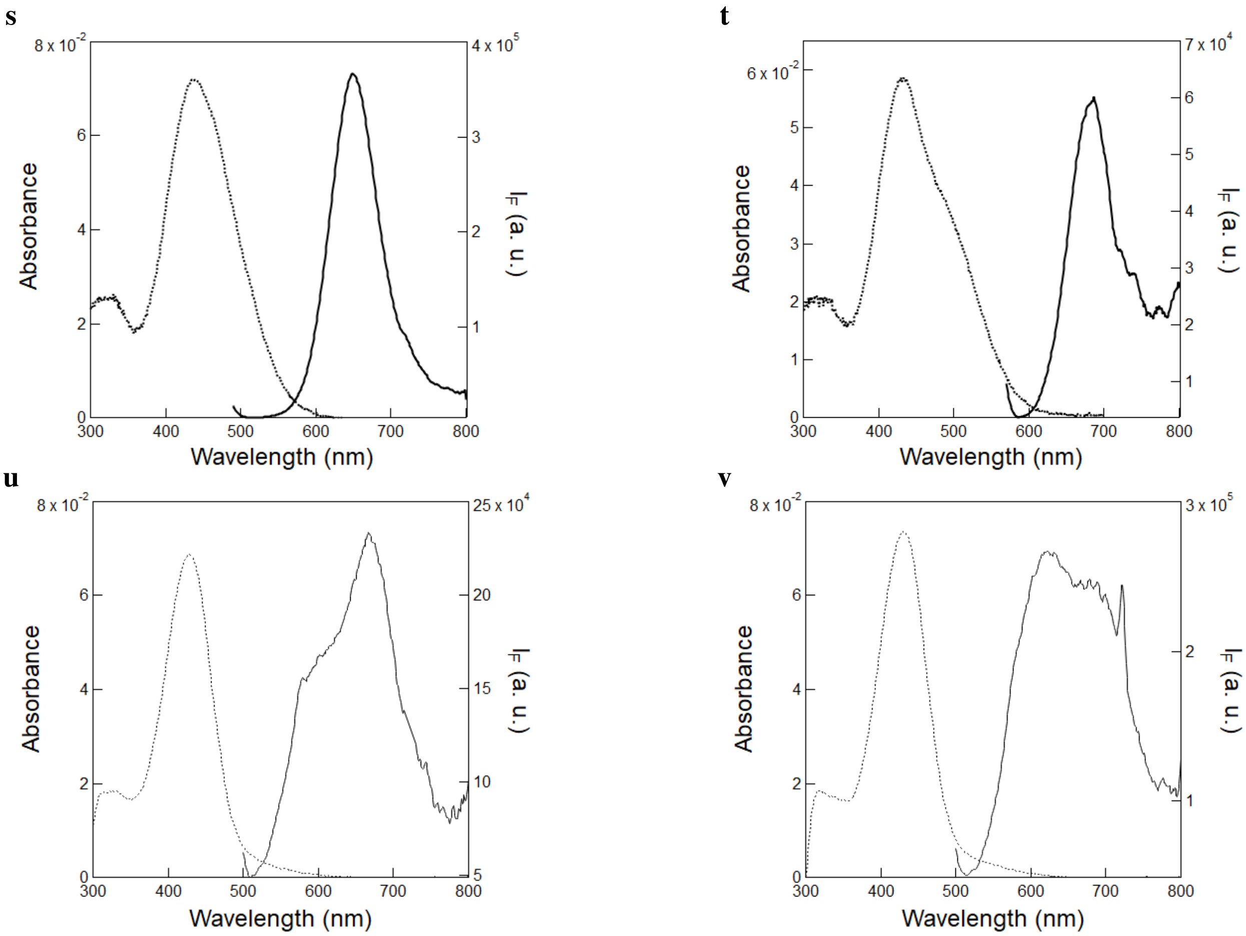

Figure S14: *UV-Vis absorption and emission spectra of the free fluorogens*. The absorption (dotted line) and emission (solid line; $\lambda_{exc}$ = 480 nm) spectra of free fluorogens **HPAR3Cl (s)**, **HPAR3F5OM (t)**, **HPAR3OM (u)** and **HPAR35DOM (v)**. The spectra were recorded using 10 $\mu$M fluorogen solution in 1$\times$ pH = 7.4 PBS buffer contained in 54 $\mu$L quartz cuvette (3 mm optical pathlength). $T = 293$ K.

#### 3.2 Measurement of the effective photoisomerization cross section of the free fluorogens

To measure the effective cross section of photoisomerization of the free fluorogens, 54 $\mu$L of 10 or 15 $\mu$M fluorogen solution in 1$\times$ pH = 7.4 PBS buffer were subjected to a sequence of light pulses of increasing constant intensity $I$ from 365, 405, or 480 nm LEDs separated by long enough periods of darkness to allow for thermal return after photoisomerization to occur. Each time evolution of the fluorescence signal recorded at its maximal emission wavelength was then satisfactorily fitted with a mono-exponential fitting function to retrieve the characteristic time of photoisomerization $\tau$.[12] The effective photoisomerization cross section of the free fluorogen was eventually extracted from the satisfactory linear fit of the inverse of $\tau$ vs light intensity $I$ using Eq. (S52) at 365, 405, or 480 nm.

##### 3.2.1 Measurements at 365 nm

Figure S15a–h displays the measurement of the effective photoisomerization cross section for **HBO3M** (Figure S15a,b), **HBO35DM** (Figure S15c,d), **HBT35DM** (Figure S15e,f), and **HBT35DOM** (Figure S15g,h) at 365 nm. From the linear fit of the inverse of $\tau$ vs light intensity $I$, we retrieved 172 $\pm$ 6, 287 $\pm$ 17, 675 $\pm$ 18, and 605 $\pm$ 34 m$^2$.mol$^{-1}$ for the effective cross section of photoisomerization of **HBO3M**, **HBO35DM**, **HBT35DM**, and **HBT35DOM** at 365 nm.

Table S1: *Photophysical properties of the free fluorogens in their thermodynamically stable (Z) configuration.* Solvent: $1\times$ pH = 7.4 PBS buffer. $T = 293$ K.

| Fluorogens | $\lambda_{abs}^{max\dagger}$ (nm) | $\lambda_{em}^{max\ddagger}$ (nm) | $\epsilon^{*}$ ($mM^{-1}cm^{-1}$) |
|---|---|---|---|
| **HBO3M** | 325 | 452 | 11 |
| **HBT35DM** | 350 | 454 | 29 |
| **HBO35DM** | 327 | 455 | 5 |
| **HBT3CN** | 380 | 483 | 23 |
| **HBT35DOM** | 358 | 491 | 21 |
| **HBR3M** | 402 | 500 | 25 |
| **HBP3CN** | 393 | 502 | 21 |
| **HBR25DM** | 407 | 505 | 26 |
| **HBTH3CN** | 422 | 519 | 39 |
| **HBR3CN** | 433 | 538 | 38 |
| **HBR35DOM** | 408 | 540 | 24 |
| **HBP35DOM** | 367 | 552 | 13 |
| **HBR3Cl5F** | 442 | 554 | 32 |
| **HBR35DF** | 439 | 555 | 34 |
| **HBR3Cl** | 446 | 555 | 30 |
| **HBIR3Cl** | 472 | 573 | 20 |
| **HBR3F5OM** | 462 | 582 | 21 |
| **HBIR35DOM** | 435 | 605 | 15 |
| **HPAR35DOM** | 430 | 620 | 25 |
| **HPAR3Cl** | 437 | 650 | 24 |
| **HPAR3OM** | 428 | 666 | 23 |
| **HPAR3F5OM** | 434 | 685 | 20 |

† $\lambda_{abs}^{max}$ wavelength of maximum absorption.
‡ $\lambda_{em}^{max}$ wavelength of maximum emission.
* $\epsilon$ molar absorption coefficient at $\lambda_{abs}^{max}$.

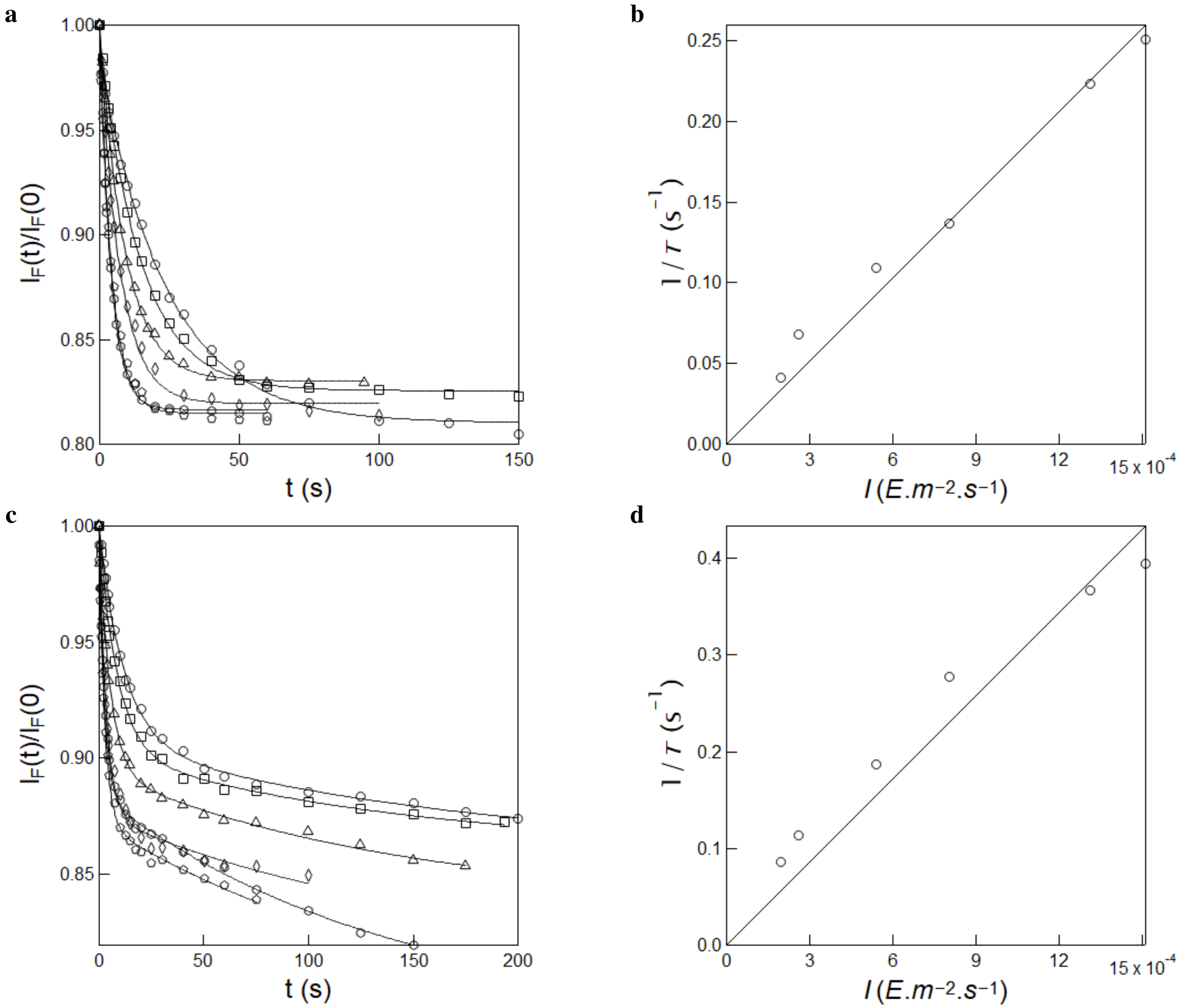

Figure S15: *Measurement of the effective cross section of photoisomerization of **HBO3M** and **HBO35DM** under 365 nm illumination.* **a,c**: Time evolution of the normalized fluorescence emission at 500 nm from 20 $\mu$M **HBO3M (a)** and 50 $\mu$M **HBO35DM (c)** in $1\times$ pH = 7.4 PBS buffer in a 54 $\mu$L cuvette (3 mm optical pathlength) upon irradiation at 365 nm at various constant light intensities (in $10^{-4}$ E.m$^{-2}$.s$^{-1}$): 2.0 (circles), 2.6 (squares), 5.4 (triangles), 8.0 (diamonds), 13.1 (pentagons) and 15.1 (hexagons). Markers: experimental data; solid lines: Monoexponential fit with linear drift delivering $\tau$ (s): 24.1, 14.7, 9.1, 7.3, 4.5, and 3.9 (for **HBO3M**) and 11.6, 8.8, 5.3, 3.6, 2.7, and 2.5 (for **HBO35DM**); **b,d**: Extraction of the effective cross section of photoisomerization for **HBO3M (b)** and **HBO35DM (d)** from the $\tau$ values retrieved in **a,c**. Markers: experimental data; solid line: linear fit with Eq.(S52). $T = 293$ K.

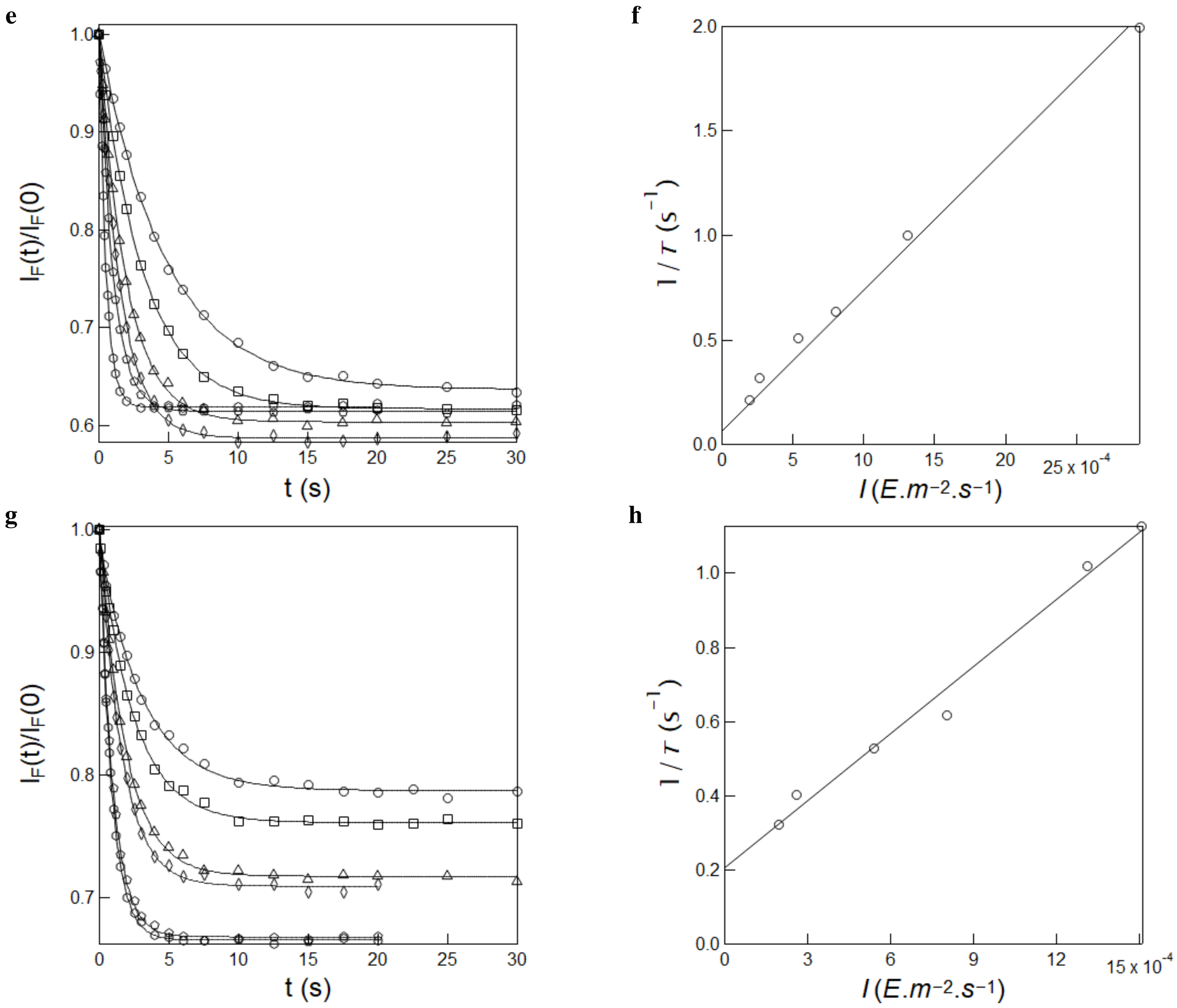

Figure S15: *Measurement of the effective cross section of photoisomerization of* ***HBT35DM*** *and* ***HBT35DOM*** *under 365 nm illumination.* **e,g**: Time evolution of the normalized fluorescence emission at 500 nm from 10 $\mu$M **HBT35DM (e)** and **HBT35DOM (g)** in 1× pH = 7.4 PBS buffer in a 54 $\mu$L cuvette (3 mm optical pathlength) upon irradiation at 365 nm at various constant light intensities (in $10^{-4}$ $E.m^{-2}.s^{-1}$): 2.0 (circles), 2.6 (squares), 5.4 (triangles), 8.0 (diamonds), 13.1 (pentagons) and 29.5 (hexagons) with **HBT35DM** and 2.0 (circles), 2.6 (squares), 5.4 (triangles), 8.0 (diamonds), 13.1 (pentagons) and 15.1 (hexagons) with **HBT35DOM**. Markers: experimental data; solid lines: Monoexponential fit delivering $\tau$ (s): 4.8, 3.2, 2.0, 1.6, 1.0, and 0.5 (for **HBT35DM**) and 3.1, 2.5, 1.9, 1.6, 1.0, and 0.9 (for **HBT35DOM**); **f,h**: Extraction of the effective cross section of photoisomerization for **HBT35DM (f)** and **HBT35DOM (h)** from the $\tau$ values retrieved in **e,g**. Markers: experimental data; solid line: linear fit with Eq.(S52). $T = 293$ K.

##### 3.2.2 Measurements at 405 nm

Figure S16a–n displays the measurement of the effective photoisomerization cross section for **HBR3Cl** (Figure S16a,b), **HBR3CN** (Figure S16c,d), **HBR3Cl5F** (Figure S16e,f), **HBR35DF** (Figure S16g,h), **HBT3CN** (Figure S16i,j), **HBP3CN** (Figure S16k,l) and **HBTH3CN** (Figure S16m,n) at 405 nm. From the linear fit of the inverse of $\tau$ vs light intensity $I$, we retrieved $1472 \pm 18$, $1691 \pm 11$, $728 \pm 11$, $962 \pm 13$. $1090 \pm 40$, $1502 \pm 57$ and $1435 \pm 35$ $m^2.mol^{-1}$ for the effective cross section of photoisomerization of **HBR3Cl**, **HBR3CN**, **HBR3Cl5F**, **HBR35DF**, **HBT3CN**, **HBP3CN** and **HBTH3CN** at 405 nm.

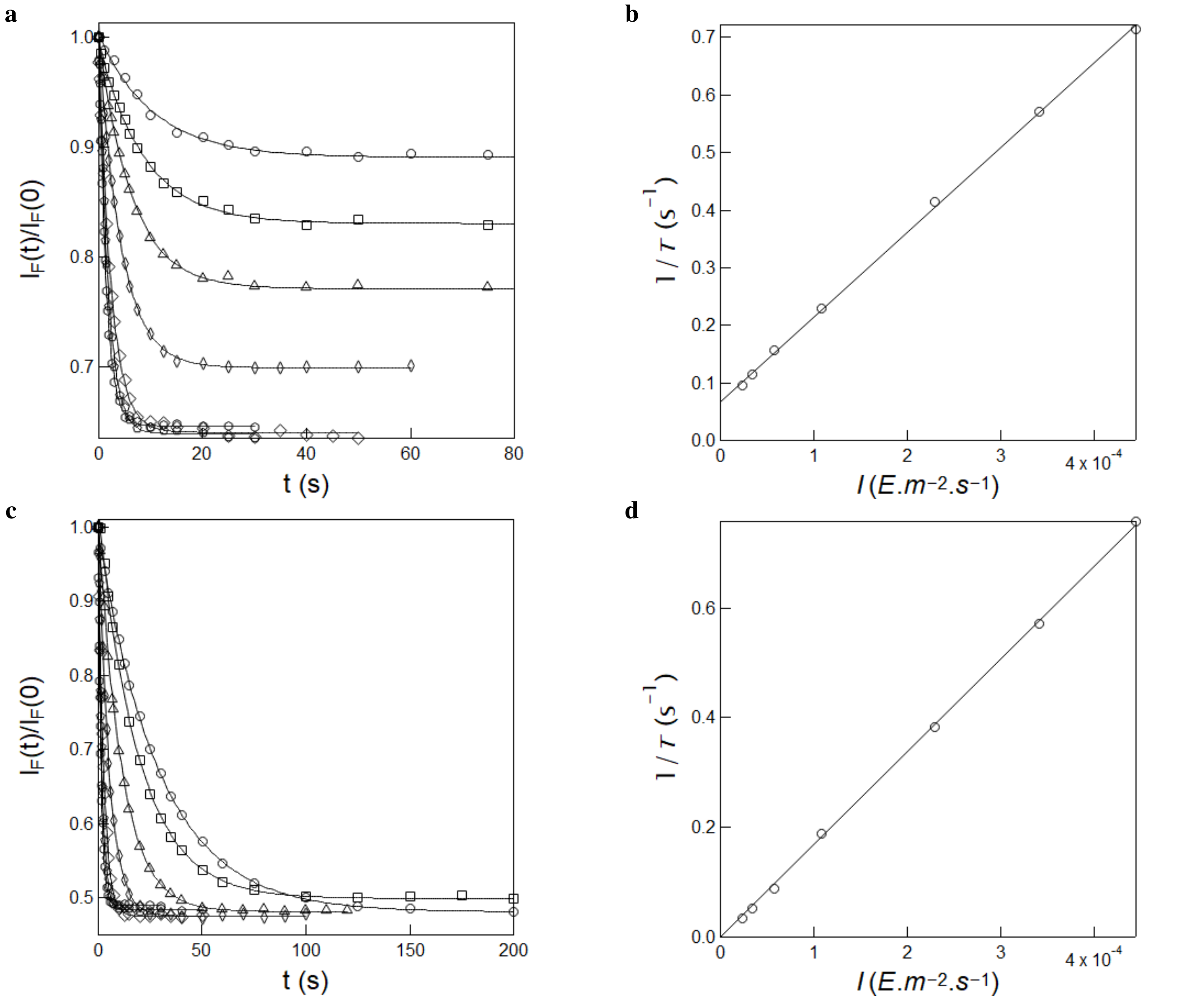

Figure S16: *Measurement of the effective cross section of photoisomerization of **HBR3Cl** and **HBR3CN*** under 405 nm illumination. **a,c**: Time evolution of the normalized fluorescence emission at 550 nm from 10 $\mu$M **HBR3Cl (a)** and **HBR3CN (c)** in 1$\times$ pH = 7.4 PBS buffer in a 54 $\mu$L cuvette (3 mm optical pathlength) upon irradiation at 405 nm at various constant light intensities (in $10^{-5}$ E.m$^{-2}$.s$^{-1}$): 2.4 (circles), 3.5 (squares), 5.7 (triangles), 10.7 (diamonds), 22.9 (discs), 34.1 (pentagons) and 44.4 (hexagons). Markers: experimental data; solid lines: Monoexponential fit delivering $\tau$ (s): 10.5, 8.7, 6.4, 4.3, 2.4, 1.7, and 1.4 (for **HBR3Cl**) and 29.5, 19.5, 11.3, 5.3, 2.6, 1.7, and 1.3 (for **HBR3CN**); **b,d**: Extraction of the effective cross section of photoisomerization for **HBR3Cl (b)** and **HBR3CN (d)** from the $\tau$ values retrieved in **a,c**. Markers: experimental data; solid line: linear fit with Eq.(S52). $T$ = 293 K.

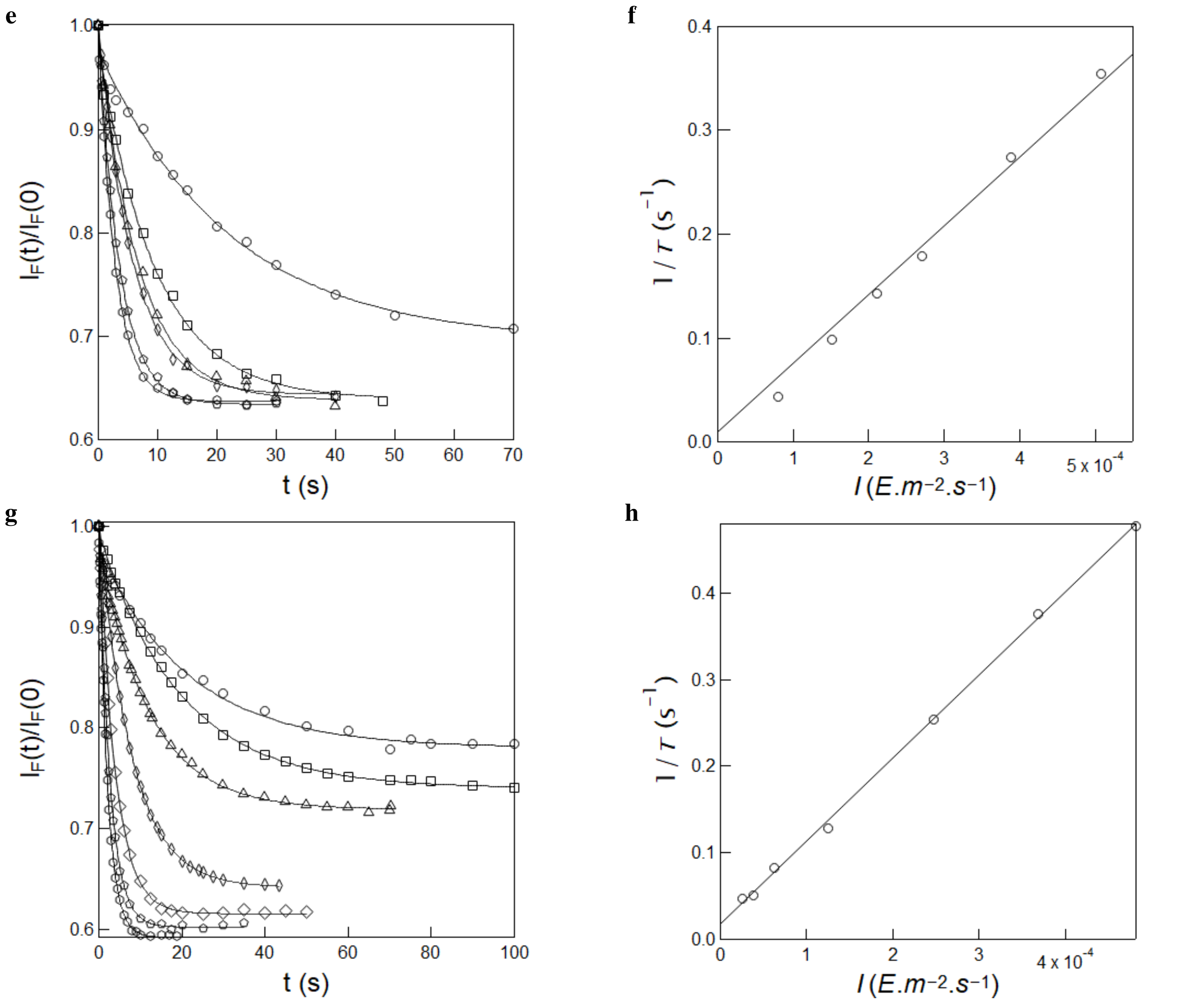

Figure S16: *Measurement of the effective cross section of photoisomerization of* ***HBR3Cl5F*** *and* ***HBR35DF*** *under 405 nm illumination.* **e,g**: Time evolution of the normalized fluorescence emission at 550 nm from 10 $\mu$M **HBR3Cl5F (e)** and **HBR35DF (g)** in 1× pH = 7.4 PBS buffer in a 54 $\mu$L cuvette (3 mm optical pathlength) upon irradiation at 405 nm at various constant light intensities (in $10^{-5}$ E.m$^{-2}$.s$^{-1}$): 7.9 (circles), 15.1 (squares), 21.0 (triangles), 27.0 (diamonds), 38.9 (pentagons) and 50.8 (hexagons) with **HBR3Cl5F**, and 2.6 (circles), 3.8 (squares), 6.2 (triangles), 12.5 (diamonds), 24.8 (disks), 36.8 (pentagons) and 48.2 (hexagons) with **HBR35DF**. Markers: experimental data; solid lines: Monoexponential fit delivering $\tau$ (s): 22.0, 9.6, 6.7, 5.7, 3.6, and 2.8 (for **HBR3Cl5F**) and 21.5, 19.7, 12.2, 7.8, 3.9, 2.6 and 2.1 (for **HBR35DF**); **f,h**: Extraction of the effective cross section of photoisomerization for **HBR3Cl5F (f)** and **HBR35DF (h)** from the $\tau$ values retrieved in **e,g**. Markers: experimental data; solid line: linear fit with Eq.(S52). $T = 293$ K.

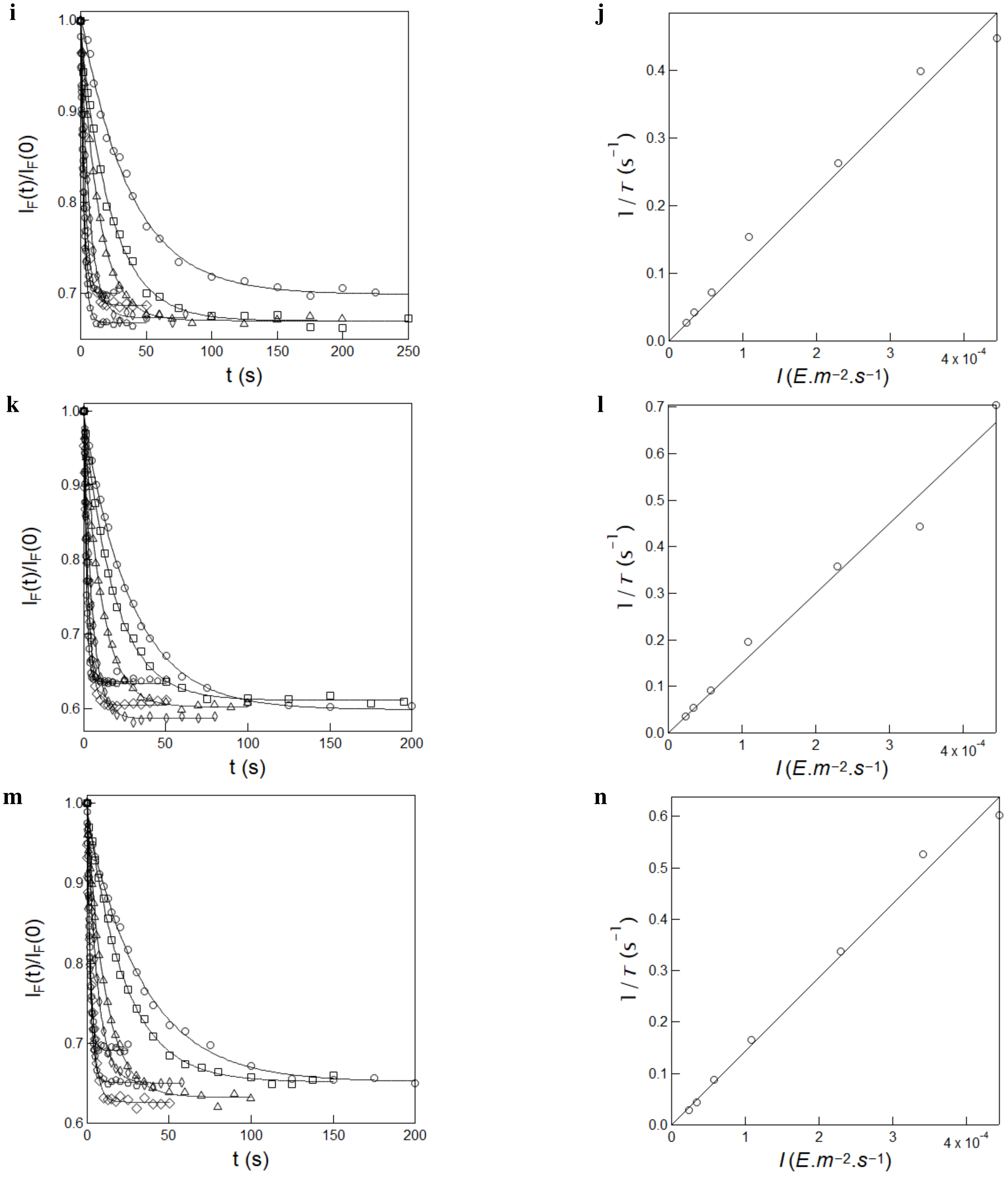

Figure S16: *Measurement of the effective cross section of photoisomerization of* ***HBT3CN****,* ***HBP3CN*** *and* ***HBTH3CN*** *under 405 nm illumination.* **i,k,m**: Time evolution of the normalized fluorescence emission at 500 nm from 10 $\mu$M **HBT3CN (i)**, **HBP3CN (k)** and **HBTH3CN (m)** in 1$\times$ pH = 7.4 PBS buffer in a 54 $\mu$L cuvette (3 mm optical pathlength) upon irradiation at 405 nm at various constant light intensities (in $10^{-5}$ E.m$^{-2}$.s$^{-1}$): 2.4 (circles), 3.5 (squares), 5.7 (triangles), 10.7 (diamonds), 22.9 (discs), 34.1 (pentagons) and 44.4 (hexagons). Markers: experimental data; solid lines: Monoexponential fit delivering $\tau$ (s): 37.7, 23.8, 13.9, 6.5, 3.8, 2.5 and 2.2 (for **HBT3CN**); 28.9, 18.7, 10.9, 5.1, 2.8, 2.2 and 1.4 (for **HBP3CN**) and 34.5, 22.7, 11.4, 6.1, 2.9, 1.9 and 1.7 (for **HBTH3CN**); **j,l,n**: Extraction of the effective cross section of photoisomerization for **HBT3CN (j)**, **HBP3CN (l)** and **HBTH3CN (n)** from the $\tau$ values retrieved in **i,k,m**. Markers: experimental data; solid line: linear fit with Eq.(S52). $T = 293$ K.

##### 3.2.3 Measurements at 480 nm

The effective cross section of photoisomerization has been similarly measured at $\lambda_{exc}$ = 480 nm. Figure S17a–l displays the results for **HBR3Cl** (Figure S17a,b), **HBR3CN** (Figure S17c,d), **HBR3Cl5F** (Figure S17e,f), **HBR35DF** (Figure S17g,h), **HPAR3Cl** (Figure S17i,j) and **HPAR3F5OM** (Figure S17k,l). Hence, we extracted 1040 ± 47, 1561 ± 31, 1367 ± 43, 1155 ± 62, 1296 ± 34 and 928 ± 29 $m^2.mol^{-1}$ at $\lambda_{exc}$ = 480 nm for **HBR3Cl**, **HBR3CN**, **HBR3Cl5F**, **HBR35DF**, **HPAR3Cl** and **HPAR3F5OM** respectively.

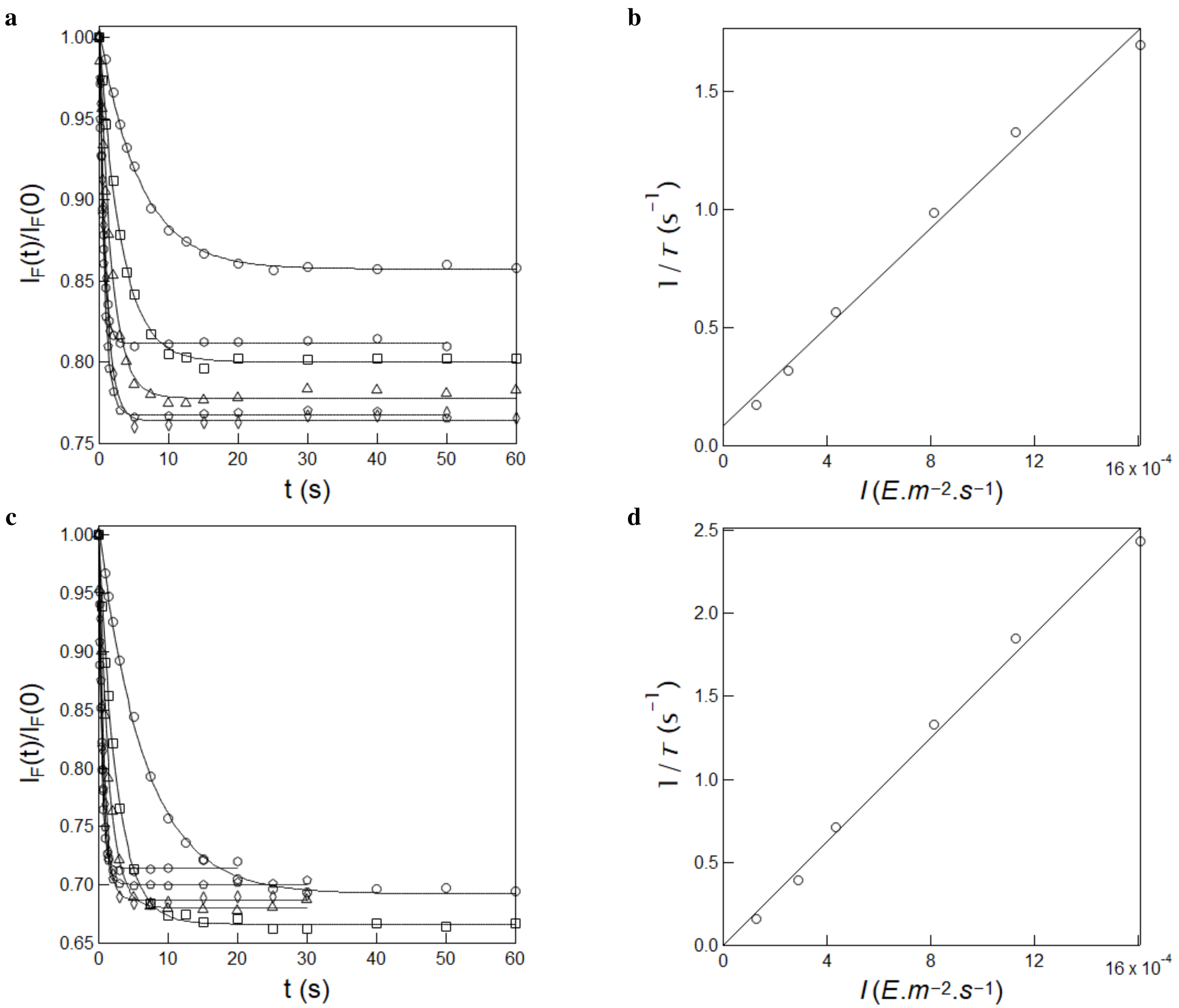

Figure S17: *Measurement of the effective cross section of photoisomerization of* ***HBR3Cl*** *and* ***HBR3CN*** under 480 nm illumination. **a,c**: Time evolution of the normalized fluorescence emission at 550 nm from 15 $\mu$M **HBR3Cl (a)** and **HBR3CN (c)** in 1× pH = 7.4 PBS buffer in a 54 $\mu$L cuvette (3 mm optical pathlength) upon irradiation at 480 nm at constant various light intensities (in $10^{-4}$ $E.m^{-2}.s^{-1}$): 1.3 (circles), 2.5 (squares), 4.4 (triangles), 8.2 (diamonds), 11.3 (pentagons) and 16.1 (hexagons). Markers: experimental data; solid lines: Monoexponential fit delivering $\tau$ (s): 5.8, 3.1, 1.8, 1.0, 0.7, and 0.6 (for **HBR3Cl**) and 6.5, 2.6, 1.5, 0.8, 0.5, and 0.4 (for **HBR3CN**); **b,d**: Extraction of the effective cross section of photoisomerization for **HBR3Cl (b)** and **HBR3CN (d)** from the $\tau$ values retrieved in **a,c**. Markers: experimental data; solid line: linear fit with Eq.(S52). $T = 293$ K.

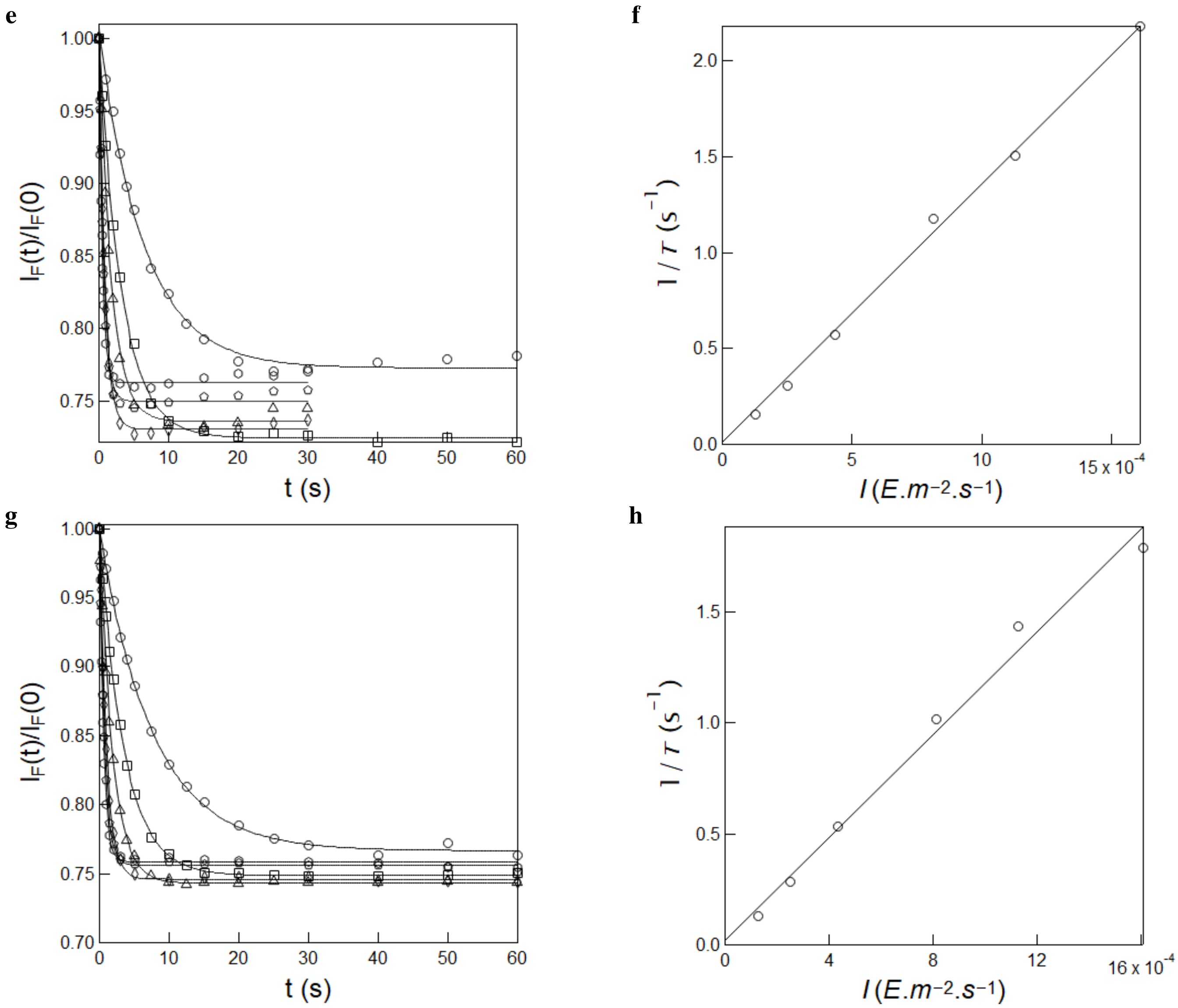

Figure S17: *Measurement of the effective cross section of photoisomerization of* ***HBR3Cl5F*** *and* ***HBR35DF*** under 480 nm illumination. **a,c**: Time evolution of the normalized fluorescence emission at 550 nm from 15 $\mu$M **HBR3Cl5F (e)** and **HBR35DF (g)** in 1$\times$ pH = 7.4 PBS buffer in a 54 $\mu$L cuvette (3 mm optical pathlength) upon irradiation at 480 nm at constant various light intensities (in $10^{-4}$ E.m$^{-2}$.s$^{-1}$): 1.3 (circles), 2.5 (squares), 4.4 (triangles), 8.2 (diamonds), 11.3 (pentagons) and 16.1 (hexagons) Markers: experimental data; solid lines: Monoexponential fit delivering $\tau$ (s): 6.4, 3.3, 1.7, 0.9, 0.7, and 0.4 (for **HBR3Cl5F**) and 7.7, 3.5, 1.9, 1.0, 0.7, and 0.6 (for **HBR35DF**); **f,h**: Extraction of the effective cross section of photoisomerization for **HBR3Cl5F (f)** and **HBR35DF (h)** from the $\tau$ values retrieved in **e,g**. Markers: experimental data; solid line: linear fit with Eq.(S52). $T = 293$ K.

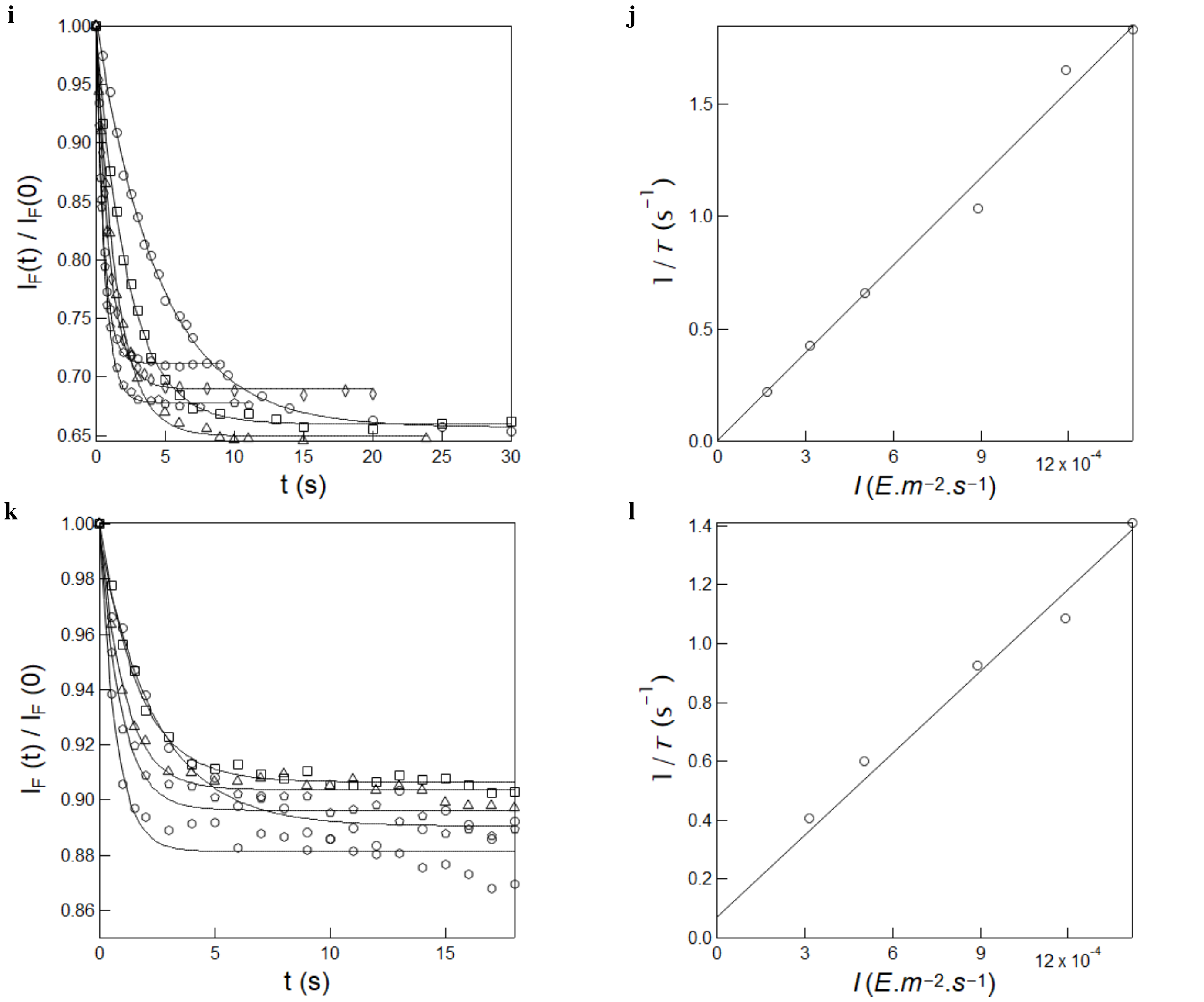

Figure S17: *Measurement of the effective cross section of photoisomerization of **HPAR3Cl** and **HPAR3F5OM** under 480 nm illumination.* **i,k**: Time evolution of the normalized fluorescence emission at 500 nm from 10 $\mu$M **HPAR3Cl (i)** and **HPAR3F5OM (k)** in 1× pH = 7.4 PBS buffer in a 54 $\mu$L cuvette (3 mm optical pathlength) upon irradiation at 480 nm at various constant light intensities (in $10^{-4}$ E.m$^{-2}$.s$^{-1}$): 1.7 (circles), 3.2 (squares), 5.0 (triangles), 8.9 (diamonds), 11.9 (pentagons) and 14.2 (hexagons) with **HPAR3Cl** and 3.2 (circles), 5.0 (squares), 8.9 (triangles), 11.9 (pentagons) and 14.2 (hexagons) with **HPAR3F5OM**. Markers: experimental data; solid lines: Monoexponential fit delivering $\tau$ (s): 4.5, 2.3, 1.5, 1.0, 0.6 and 0.5 (for **HPAR3Cl**) and 2.5, 1.7, 1.1, 0.9, and 0.7 (for **HPAR3F5OM**); **j,l**: Extraction of the effective cross section of photoisomerization for **HPAR3Cl (j)** and **HPAR3F5OM (l)** from the $\tau$ values retrieved in **i,k**. Markers: experimental data; solid line: linear fit with Eq.(S52). $T = 293$ K.

##### 3.3 Kinetics of thermal recovery from the photoisomerized state of the free fluorogen

To measure the effective rate constant associated with thermal recovery of the free fluorogens, 54 $\mu$L of 10 $\mu$M fluorogen solution in $1\times$ pH = 7.4 PBS buffer was repeatedly illuminated with 365 ($I$=13.1×10$^{-4}$ E.m$^{-2}$.s$^{-1}$), 405 ($I$ = 3.7×10$^{-4}$ E.s$^{-1}$.m$^{-2}$) or 480 ($I$=14.2×10$^{-4}$ E.m$^{-2}$.s$^{-1}$) nm LED, and the fluorescence signal was monitored at the respective emission maximum until the photostationary state was reached. These illumination steps were separated by increasing delay times ($\Delta t$), where solutions were kept at complete darkness. Recovery of the fluorescence signal was measured and plotted against the corresponding delay time $\Delta t$. The relaxation time associated with thermal recovery of the photoisomerized state was then extracted from the monoexponentially fitting of the dependence of the fluorescence recovery on the delay time $\Delta t$ by using Eq.(S51). The effective rate constant associated with thermal recovery was eventually retrieved as the inverse of the extracted characteristic time.

The fluorescence level recovered after each relaxation in the darkness was satisfactorily fitted monoexponentially with Eq.(S51) for most of the investigated fluorogens. Hence, we extracted $13.0 \pm 0.7$, $3359 \pm 336$, $96 \pm 5$, $54 \pm 5$, $16 \pm 1$, $7 \pm 0.4$, $238 \pm 15$, and $15 \pm 3$ s for the characteristic time of thermal return for **HBR3Cl** (Figure S18b), **HBR3CN** (Figure S18d), **HBR3Cl5F** (Figure S18f), and **HBR35DF** (Figure S18h), **HBT35DM** (Figure S18j) and **HBT35DOM** (Figure S18l),**HPAR3Cl** (Figure S18n) and **HPAR3F5OM** (Figure S18p) respectively, which finally yielded $k^{\Delta}_{F'F} = 0.08$, $3.0 \times 10^{-4}$, 0.01, 0.02, 0.06, 0.14, $4.2 \times 10^{-3}$, and 0.07 s$^{-1}$ at 293 K for **HBR3Cl**, **HBR3CN**, **HBR3Cl5F**, **HBR35DF**, **HBT35DM**, **HBT35DOM**, **HPAR3Cl** and **HPAR3F5OM** respectively.

In contrast, although exhibit photoisomerization under illumination at 365 nm, we noticed that the photoisomerized state of **HBO3M**, **HBO35DM**, **HBT3CN**, **HBP3CN** and **HBTH3CN** essentially did not recover over a time scale of an hour (Figure S19a–e).

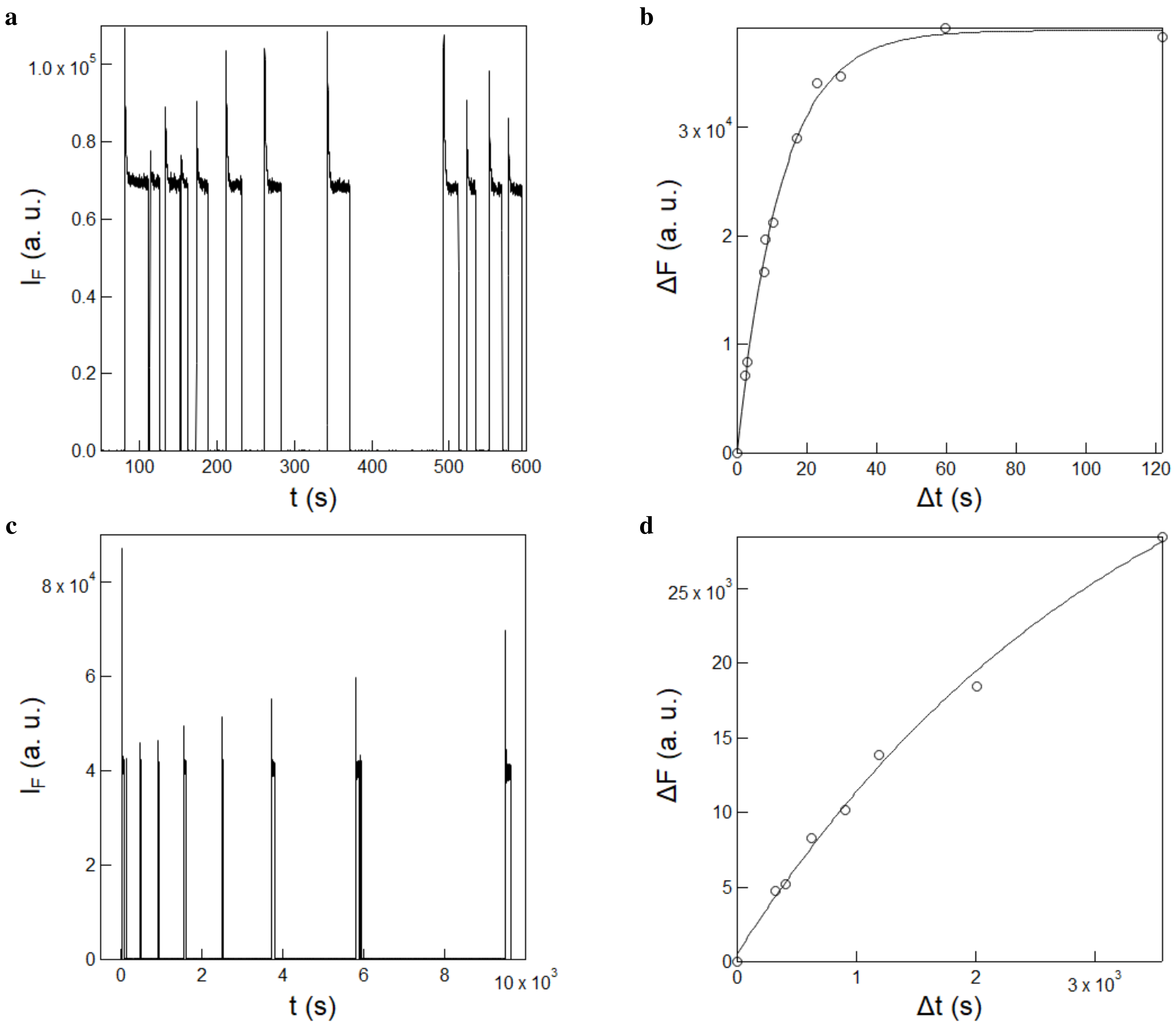

Figure S18: *Measurement of the rate constant of thermal recovery from the photoisomerized state of **HBR3Cl** and **HBR3CN***. **a,c**: Time evolution of the fluorescence emission at 550 nm from 10 $\mu$M **HBR3Cl** (**a**) and **HBR3CN** (**c**) in 1× pH = 7.4 PBS buffer exposed to a sequence of long enough 405 nm light pulses at 2.3×$10^{-4}$ E.m$^{-2}$.s$^{-1}$ to reach the photostationary state while varying the time of recovery in the dark in between the pulses; **b,d**: Extraction of the rate constant associated with the thermal return of the photoisomerized **HBR3Cl** (**b**) and **HBR3CN** (**d**) stereoisomer from the recovery of fluorescence signal against the delay time ($\Delta t$) retrieved in **a** and **c**. Markers: experimental data; solid line: monoexponential fit using Eq.(S51). $T = 293$ K.

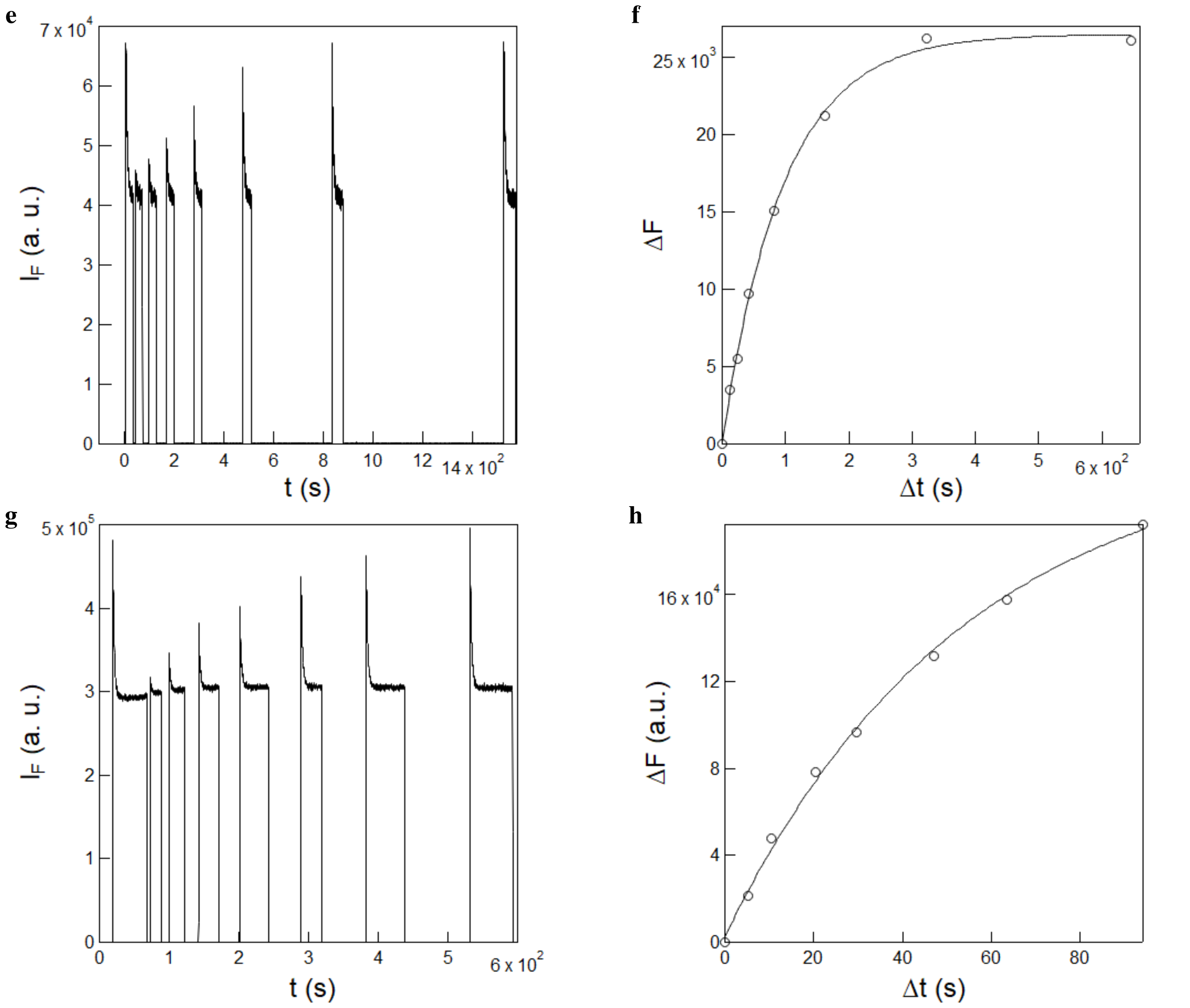

Figure S18: *Measurement of the rate constant of thermal recovery from the photoisomerized state of **HBR3Cl5F** and **HBR35DF**.* **e,g**: Time evolution of the fluorescence emission at 550 nm from 10 $\mu$M **HBR3Cl5F** (**e**) and **HBR35DF** (**g**) in $1\times$ pH = 7.4 PBS buffer exposed to a sequence of long enough 405 nm light pulses at 2.3×$10^{-4}$ E.m$^{-2}$.s$^{-1}$ to reach the photostationary state while varying the time of recovery in the dark in between the pulses; **f,h**: Extraction of the rate constant associated with the thermal return of the photoisomerized **HBR3Cl5F** (**f**) and **HBR35DF** (**h**) stereoisomer from the recovery of fluorescence signal against the delay time ($\Delta t$) retrieved in **e** and **g**. Markers: experimental data; solid line: monoexponential fit using Eq.(S51). $T = 293$ K.

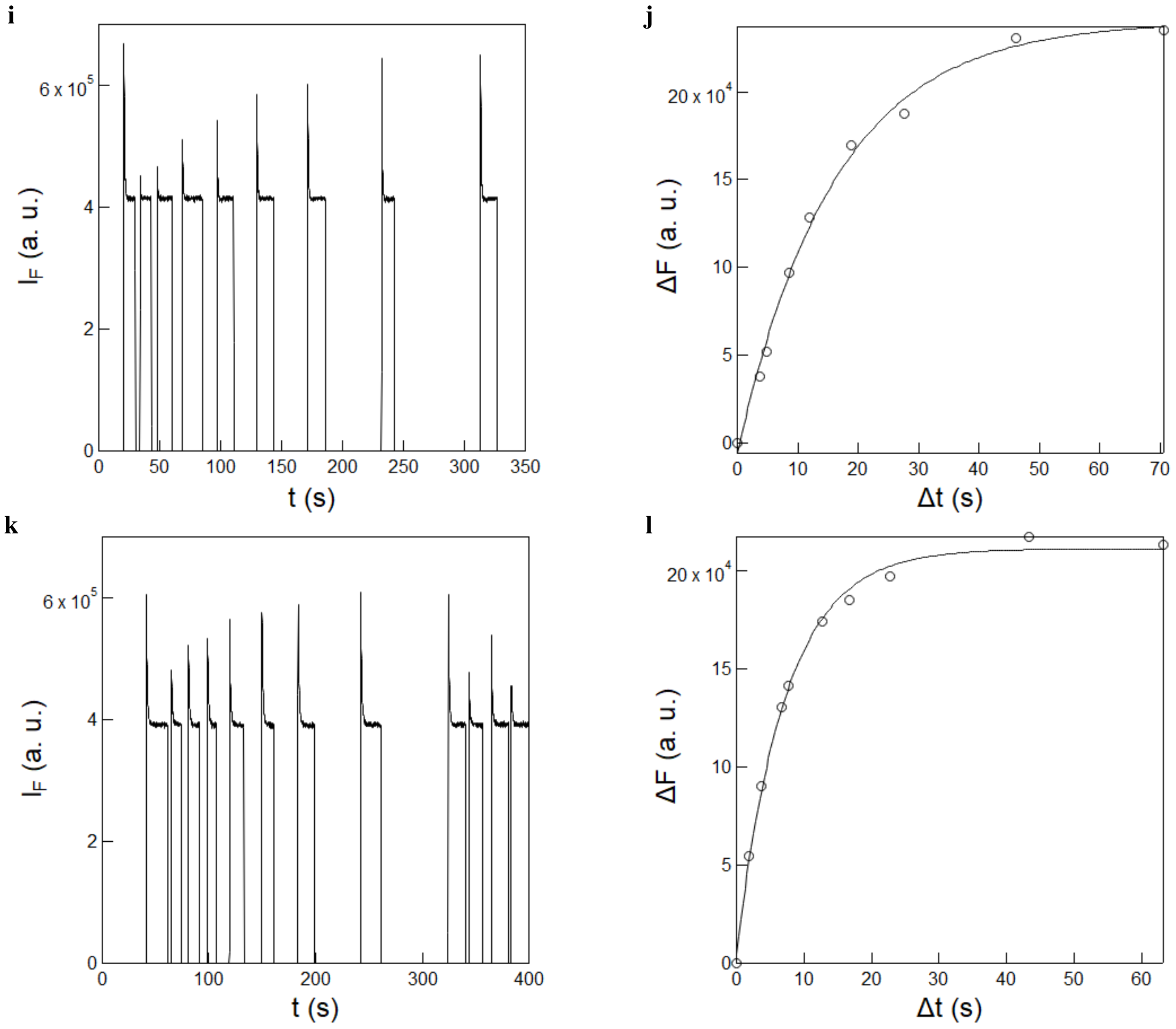

Figure S18: *Measurement of the rate constant of thermal recovery from the photoisomerized state of* ***HBT35DM*** *and* ***HBT35DOM***. **i,k**: Time evolution of the fluorescence emission at 500 nm from 10 $\mu$M **HBT35DM** (**i**) and **HBT35DOM** (**k**) in 1$\times$ pH = 7.4 PBS buffer exposed to a sequence of long enough 365 nm light pulses at 13.1$\times$10$^{-4}$ E.m$^{-2}$.s$^{-1}$ to reach the photostationary state while varying the time of recovery in the dark in between the pulses; **j,l**: Extraction of the rate constant associated with the thermal return of the photoisomerized **HBT35DM** (**j**) and **HBT35DOM** (**l**) stereoisomer from the recovery of fluorescence signal against the delay time ($\Delta t$) retrieved in **i** and **k**. Markers: experimental data; solid line: monoexponential fit using Eq.(S51). $T = 293$ K.

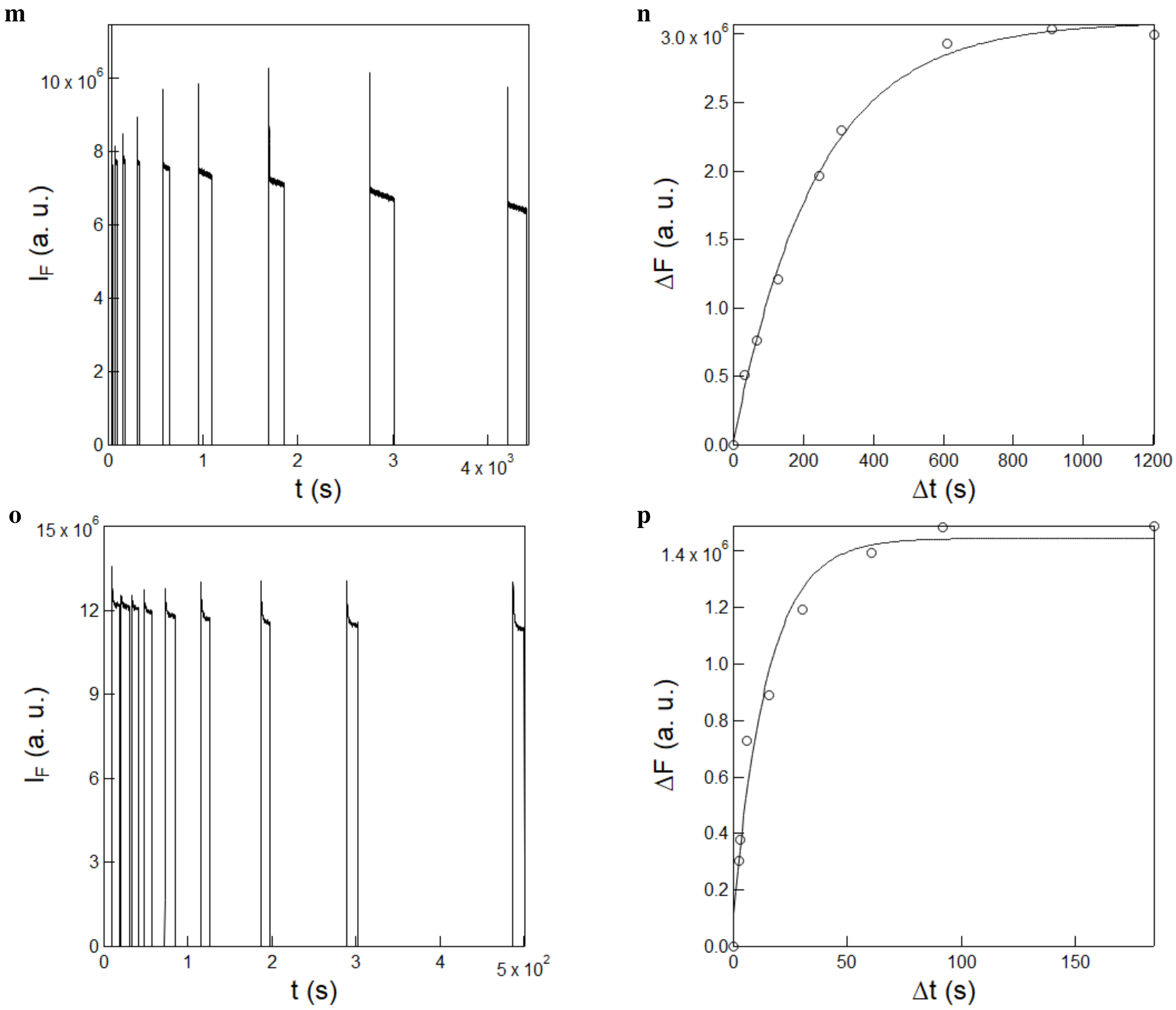

Figure S18: *Measurement of the rate constant of thermal recovery from the photoisomerized state of* ***HPAR3Cl*** *and* ***HPAR3F5OM***. **m,o**: Time evolution of the fluorescence emission at 660 nm from 10 $\mu$M **HPAR3Cl** (**m**) and **HPAR3F5OM** (**o**) in 1$\times$ pH = 7.4 PBS buffer exposed to a sequence of long enough 480 nm light pulses at 14.2$\times10^{-4}$ E.m$^{-2}$.s$^{-1}$ to reach the photostationary state while varying the time of recovery in the dark in between the pulses; **n,p**: Extraction of the rate constant associated with the thermal return of the photoisomerized **HPAR3Cl** (**n**) and **HPAR3F5OM** (**p**) stereoisomer from the recovery of fluorescence signal against the delay time ($\Delta t$) retrieved in **m** and **o**. Markers: experimental data; solid line: monoexponential fit using Eq.(S51). $T = 293$ K.

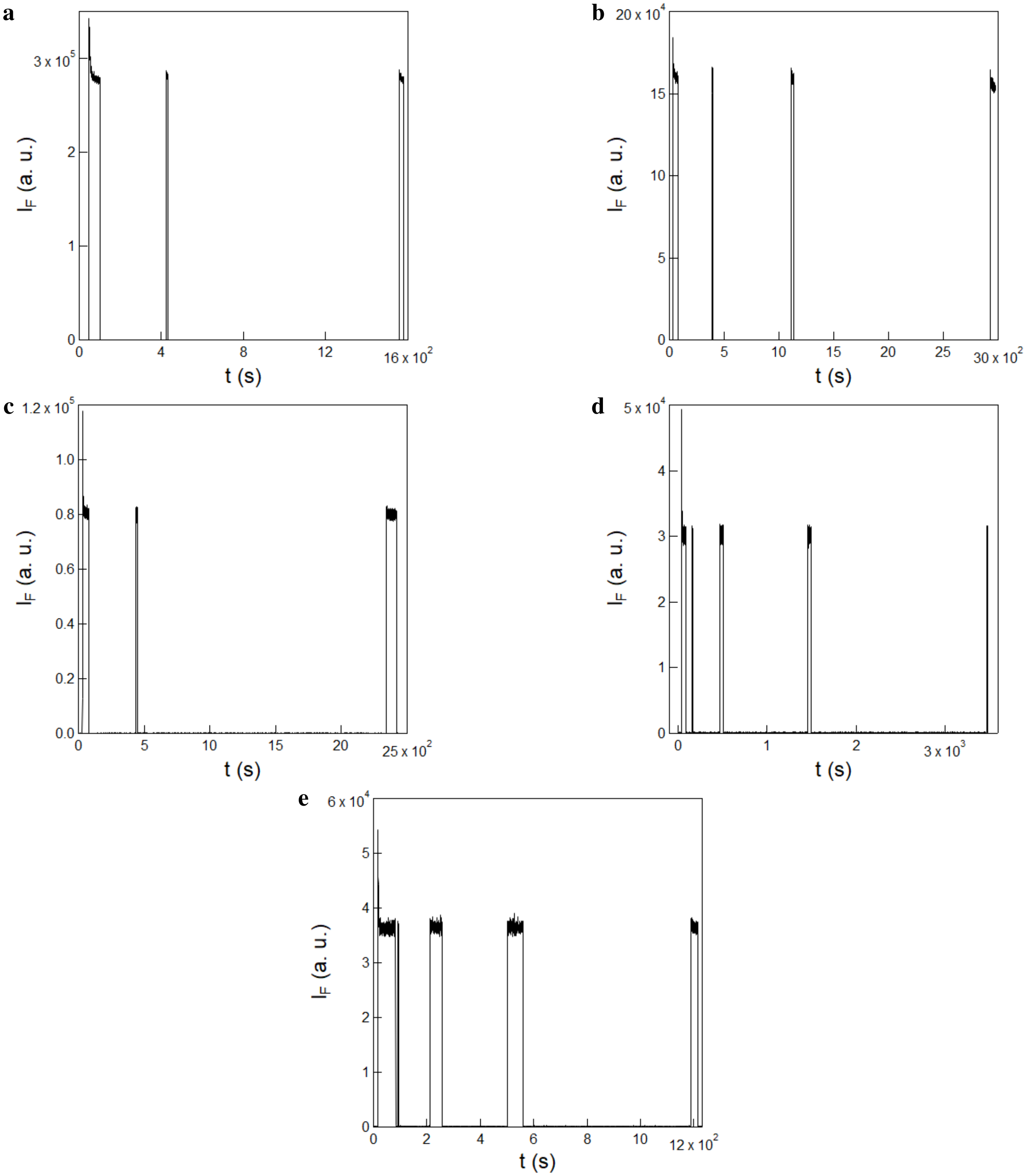

Figure S19: *Attempt to measure the rate constant of thermal recovery from the photoisomerized state of* ***HBO3M****,* ***HBO35DM****,* ***HBT3CN****,* ***HBP3CN*** *and* ***HBTH3CN***. Time evolution of the fluorescence emission at 500 nm from 20 $\mu$M **HBO3M**, **HBO35DM** and 10 $\mu$M **HBT3CN**, **HBP3CN** and **HBTH3CN** in 1$\times$ pH = 7.4 PBS buffer exposed to a sequence of long enough 365 nm (15.1×$10^{-4}$ E.m$^{-2}$.s$^{-1}$) for **HBO3M**, **HBO35DM** or 405 nm (1.1×$10^{-4}$ E.m$^{-2}$.s$^{-1}$) for **HBT3CN**, **HBP3CN** and **HBTH3CN** light pulses to reach the photostationary state while varying the time of recovery in the dark in between the pulses. $T$ = 293 K.

We conducted analogous photochemical experiments on **HBR3F5OM**, **HBIR3Cl**, **HBR25DM** and **HBR35DOM** using either 405 or 480 nm LED as the light source. Under identical conditions, we observed a constant fluorescence intensity over time for these fluorogens (Figure S20a,b). This observation may have led us to deduce that these fluorogens do not undergo photoisomerization. However, except for **HBR35DOM**, their photoisomerization was observed in imaging experiments performed with confocal microscopy. Hence, we rather concluded that these fluorogens possess extremely rapid thermal recovery (<1 s) so as to forbid to observe any fluorescence change upon applying a light jump at the light intensity ($10^{-3}$ E.m$^{-2}$.s$^{-1}$ range) employed.

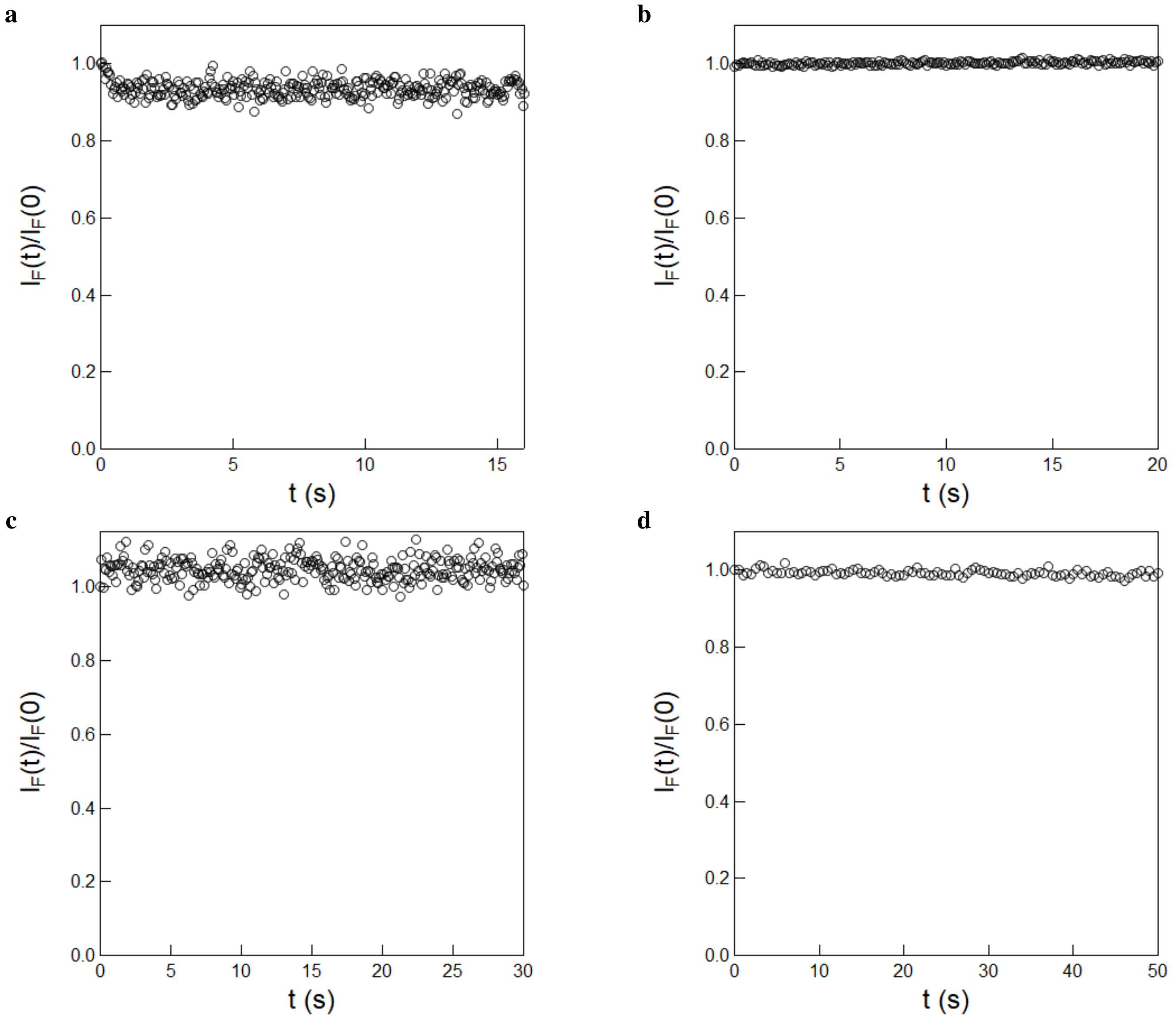

Figure S20: *An attempt to measure the rate constant of thermal recovery from the photoisomerized state of* ***HBR3F5OM****,* ***HBIR3Cl****,* ***HBR25DM*** *and* ***HBR35DOM***. Time evolution of the fluorescence emission at 550 nm from 10 $\mu$M **HBR3F5OM (a)**, **HBIR3Cl (b)**, **HBR25DM (c)** and **HBR35DOM (d)** in 1× pH = 7.4 PBS buffer in a 54 $\mu$L cuvette (3 mm optical pathlength) upon irradiation at 405 nm (for **HBR3F5OM**, **HBR25DM** and **HBR35DOM** ) and 480 nm (for **HBIR3Cl**) at constant light intensity. $T = 293$ *K*.

#### 3.4 Measurement of the proton exchange constants of the free fluorogens

To determine the proton exchange constant of the free fluorogens in their ($Z$) thermodynamically stable state, we used 2–4 $\mu$M fluorogen solutions in 10 mM universal buffer prepared as follows: To 100 mL of Milli-Q water were dissolved 82 mg of sodium acetate ($CH_3COONa$), 120 mg of sodium dihydrogenophosphate ($NaH_2PO_4$), and 518.2 mg of 2-(cyclohexylamino)ethane-1-sulfonic acid (CHES). The pH of the initial solution (pH = 5.90) was then adjusted to pH = 9 with 1 M NaOH. The dependence of the absorption spectrum obeying Eq.(S50) was globally analyzed by exploiting the SPECFIT/32TM Global Analysis System (Version 3.0 for 32-bit Windows systems)[14–17] to extract $K_{\mathbf{FH}}$.
Figures S21a–m display the pH-dependence of the absorption spectrum of **HBR3Cl**, **HBR3CN**, **HBR3Cl5F**, **HBR35DF**, **HBR3F5OM**, **HBIR3Cl**, **HBT35DOM**, **HBIR35DOM**, **HBT3CN**, **HBP3CN**, **HBTH3CN**, **HPAR3Cl**, and **HPAR3F5OM**. The extracted values of p$K_{\mathbf{FH}}$ are given in Table S2.

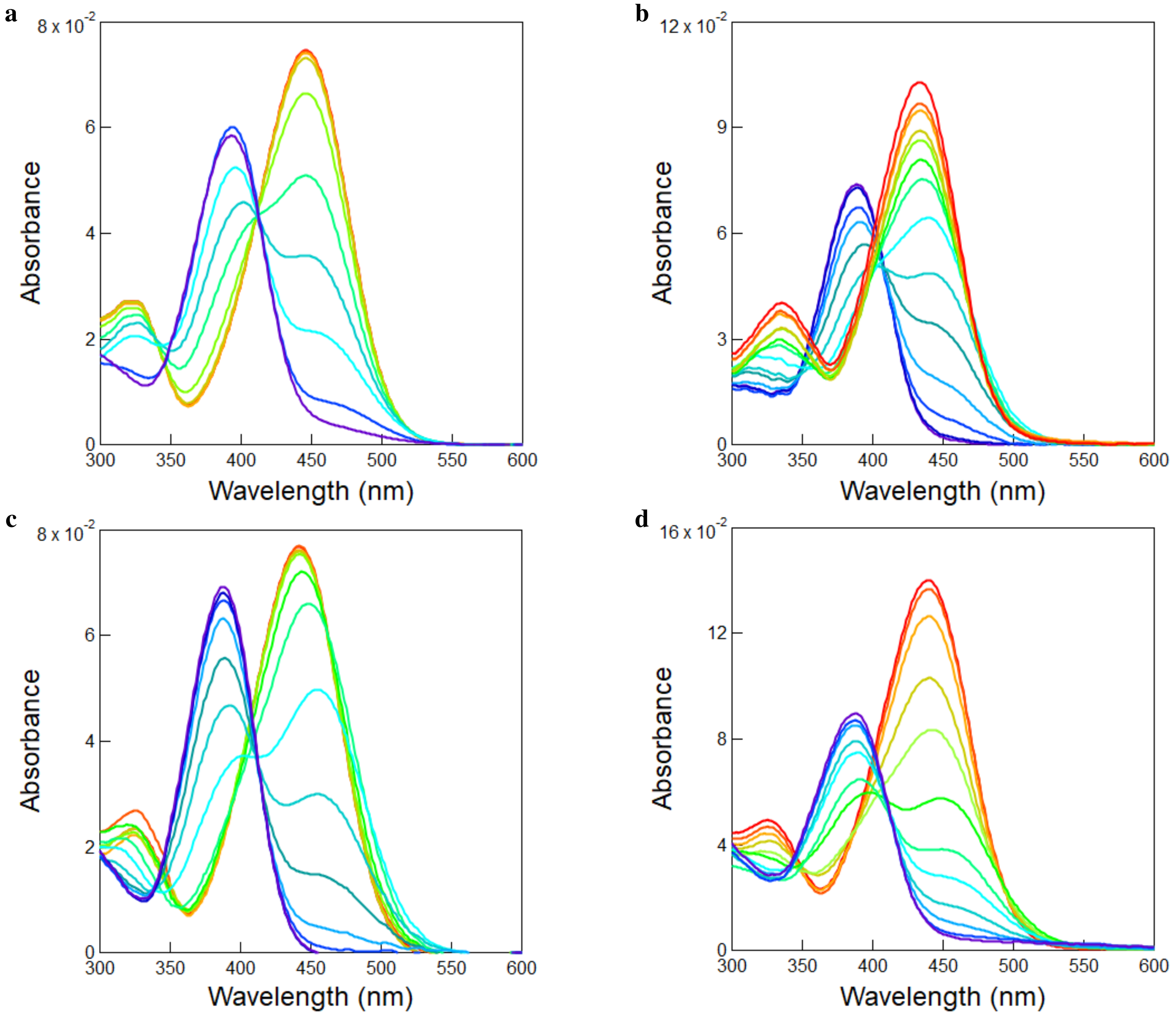

Figure S21: *Determination of the proton exchange constant $K_{FH}$ of the free fluorogens.* pH-dependence of the absorption spectrum of 2.4 $\mu$M **HBR3Cl (a)**, 3 $\mu$M **HBR3CN (b)**, 2.4 $\mu$M **HBR3Cl5F (c)** and 4 $\mu$M **HBR35DF (d)** in 10 mM universal buffer. pH = 9.9, 9.4, 9.0, 8.5, 7.7, 7.1, 6.7, 6.2, 5.4, 4.8 for **HBR3Cl**; pH = 9.0, 8.8, 8.4, 7.8, 7.5, 7.1, 6.7, 6.1, 5.6, 5.0, 4.6, 4.2, 3.1, 2.6 for **HBR3CN**; pH = 9.7, 8.8, 8.3, 7.5, 7.0, 6.6, 6.0, 5.2, 4.7, 4.3, 3.7, 3.3, 2.9, 2.7 for **HBR3Cl5F**; pH = 9.0, 8.5, 7.8, 7.1, 6.6, 5.7, 5.3, 4.9, 4.6, 4.2, 3.6, 3.0 for **HBR35DF**. The extracted values of p$K_{\mathbf{FH}}$ are 6.8, 5.1, 4.8 and 5.3 for **HBR3Cl**, **HBR3CN**, **HBR3Cl5F** and **HBR35DF** respectively. $T = 293$ K.

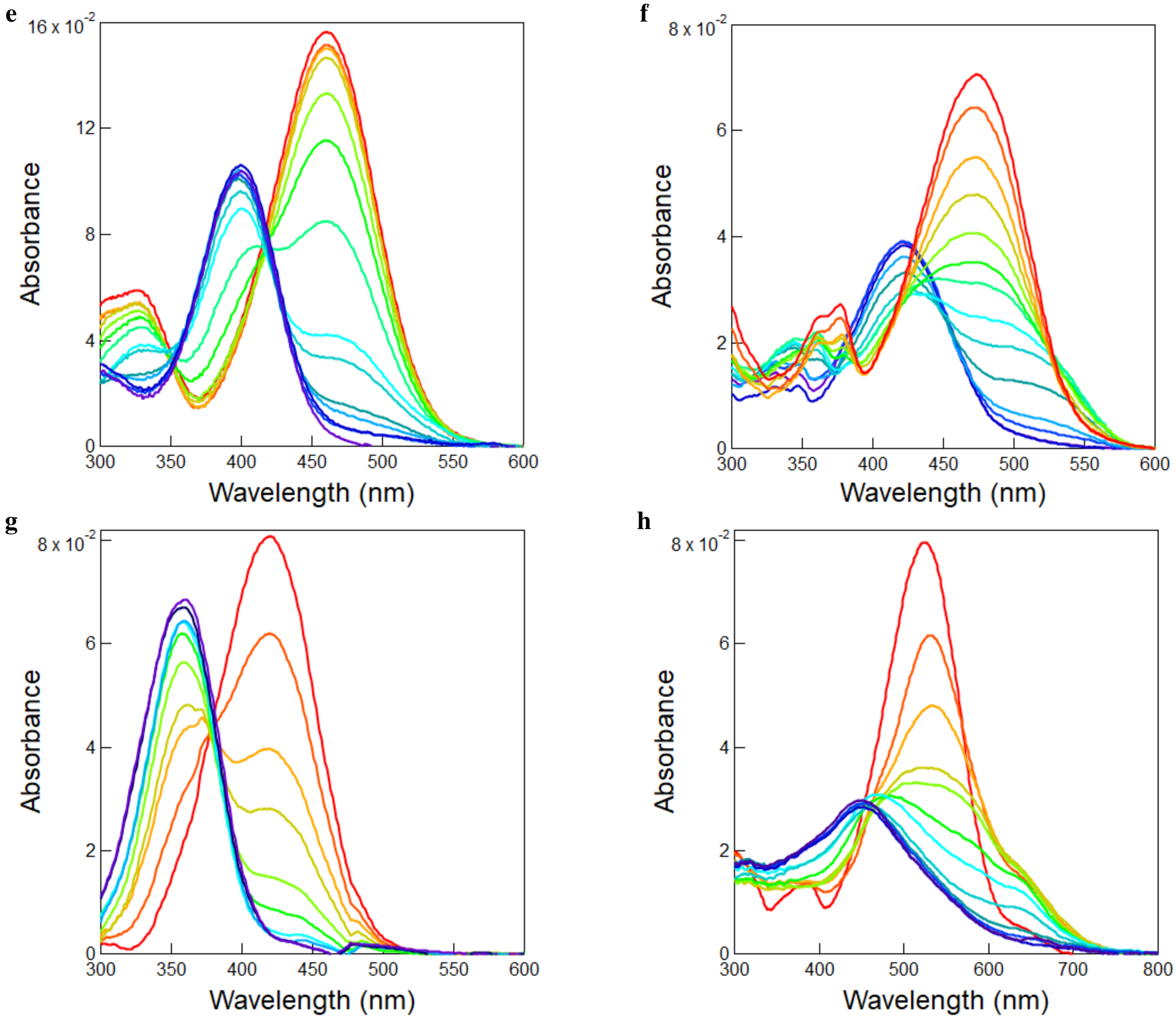

Figure S21: *Determination of the proton exchange constant $K_{FH}$ of the free fluorogens.* pH-dependence of the absorption spectrum of **HBR3F5OM (e)**, **HBIR3Cl (f)**, **HBT35DOM (g)** and **HBIR35DOM (h)** recorded at 3 $\mu$M in 10 mM universal buffer. pH = 9.9, 9.5, 9.0, 8.6, 8.0, 7.5, 7.0, 6.5, 6.1, 5.5, 5.1, 4.7, 3.9, 3.3 for **HBR3F5OM**, pH = 3.0, 3.8, 4.8, 5.4, 6.0, 6.3, 6.7, 6.9, 7.1, 7.3, 7.7, 8.1, 8.8, 9.1 for **HBIR3Cl**, pH = 9.6, 9.0, 8.6, 8.0, 7.7, 7.3, 6.9, 6.5, 5.5, 4.5 for **HBT35DOM** and pH = 9.8, 9.3, 8.9, 8.6, 7.9, 7.7, 7.1, 6.8, 6.3, 5.2, 4.6, 3.9 for **HBIR35DOM**. The extracted values of p$K_{\mathbf{FH}}$ are 6.9, 7.1, 8.6 and 8.4 for **HBR3F5OM**, **HBIR3Cl**, **HBT35DOM** and **HBIR35DOM** respectively. $T = 293$ K.

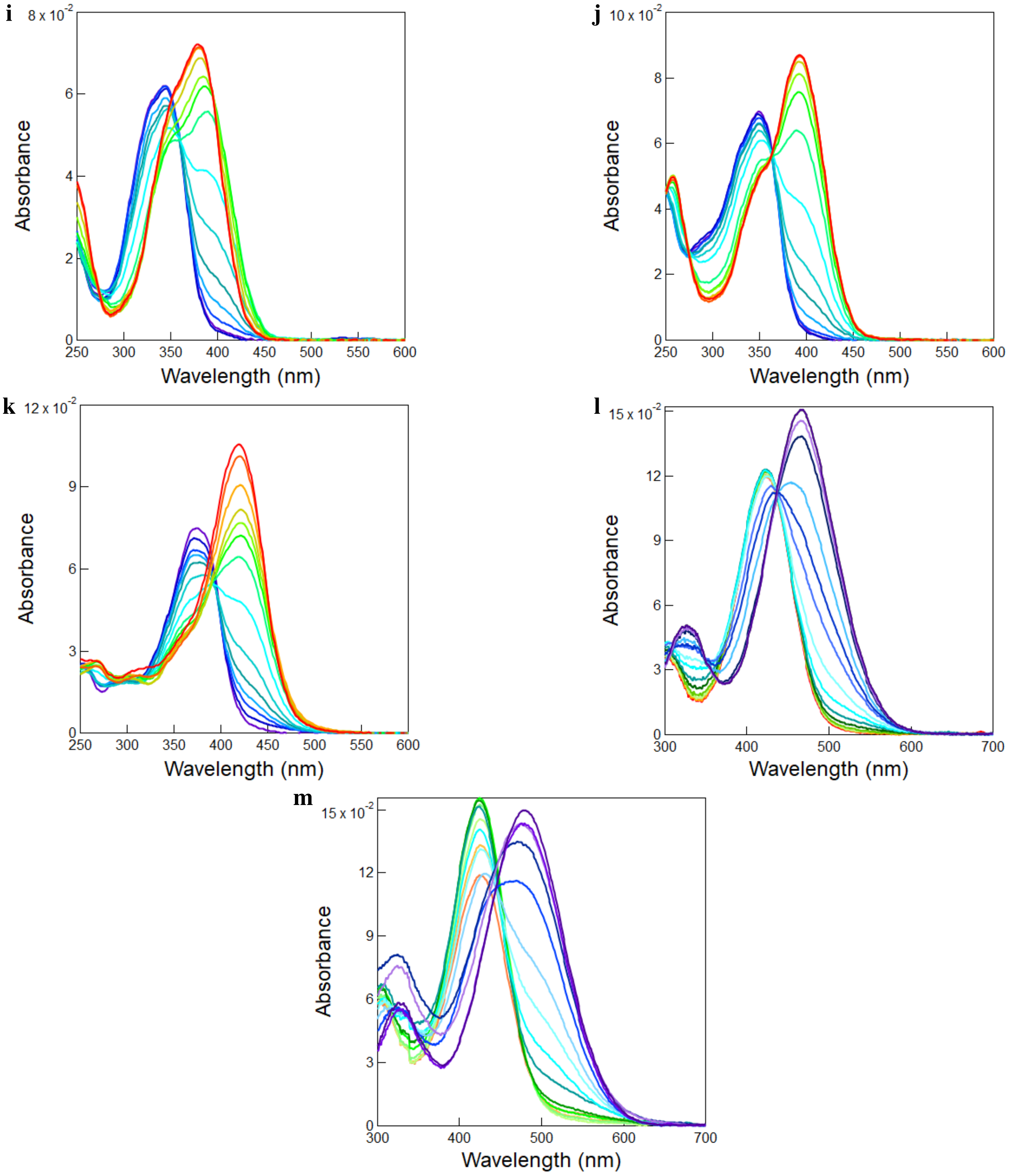

Figure S21: *Determination of the proton exchange constant $K_{FH}$ of the free fluorogens.* pH-dependence of the absorption of **HBT3CN (i)**, **HBP3CN (j)**, **HBTH3CN (k)** recorded at 3 $\mu$M and **HPAR3Cl (l)**, **HPAR3F5OM (m)** recorded at 4 $\mu$M in 10 mM universal buffer. pH = 9.0, 8.8, 8.4, 7.8, 7.5, 7.1, 6.7, 6.1, 5.6, 5.0, 4.6, 4.2, 3.1, 2.6 for **HBR3CN**; pH = 9.0, 8.6, 8.1, 7.4, 6.8, 6.5, 6.1, 5.5, 5.1, 4.8, 4.4, 3.9, 3.1, 2.6 for **HBT3CN**; pH = 9.0, 8.6, 8.1, 7.4, 6.9, 6.6, 6.2, 5.6, 5.1, 4.8, 4.5, 4.0, 3.2, 2.6 for **HBP3CN**, pH = 9.0, 8.7, 8.1, 7.3, 7.0, 6.6, 6.3, 5.8, 5.3, 4.9, 4.6, 4.3, 3.4, 2.7 for **HBTH3CN**, pH = 2.9, 3.1, 3.8, 4.2, 4.6, 4.6, 4.9, 5.2, 5.6, 5.9, 6.5, 6.8, 7.1, 7.2, 7.5, 8.5, 8.8, 9.1 for **HPAR3Cl** and pH = 3.8, 4.1, 4.7, 4.9, 5.4, 5.9, 6.5, 7.0, 7.3, 7.6, 8.1, 9.0 for **HPAR3F5OM**. The extracted p$K_a$ values are 5.3, 5.2, 5.6, 7.7 and 7.7 for **HBT3CN**, **HBP3CN**, **HBTH3CN**, **HPAR3Cl** and **HPAR3F5OM** respectively. $T$ = *293 K*.

Table S2: *Proton exchange properties of the (Z) stereoisomer of the free fluorogens.* Solvent: $1\times$ pH = 7.4 PBS buffer. $T = 293$ K.

| Fluorogens | $pK_a$ |
|---|---|
| **HBR3Cl5F** | 4.8 |
| **HBR3CN** | 5.1 |
| **HBP3CN** | 5.2 |
| **HBT3CN** | 5.3 |
| **HBR35DF** | 5.3 |
| **HBTH3CN** | 5.6 |
| **HBR3Cl** | 6.8 |
| **HBR3F5OM** | 6.9 |
| **HBIR3Cl** | 7.1 |
| **HPAR3Cl** | 7.7 |
| **HPAR3F5OM** | 7.7 |
| **HBIR35DOM** | 8.4 |
| **HBT35DOM** | 8.6 |

### 4 Investigation of the pFAST-fluorogen and nirFAST-fluorogen complexes

#### 4.1 Absorption and emission spectra

The absorption and emission spectra of the **pFAST**-**fluorogen** and **nirFAST**-**fluorogen** complexes in $1\times$ pH = 7.4 PBS buffer have been recorded in 54 $\mu$L quartz cuvette (3 mm optical pathlength) at protein concentration much higher than the nanomolar range of the dissociation constant and with an excess of protein over the fluorogen.

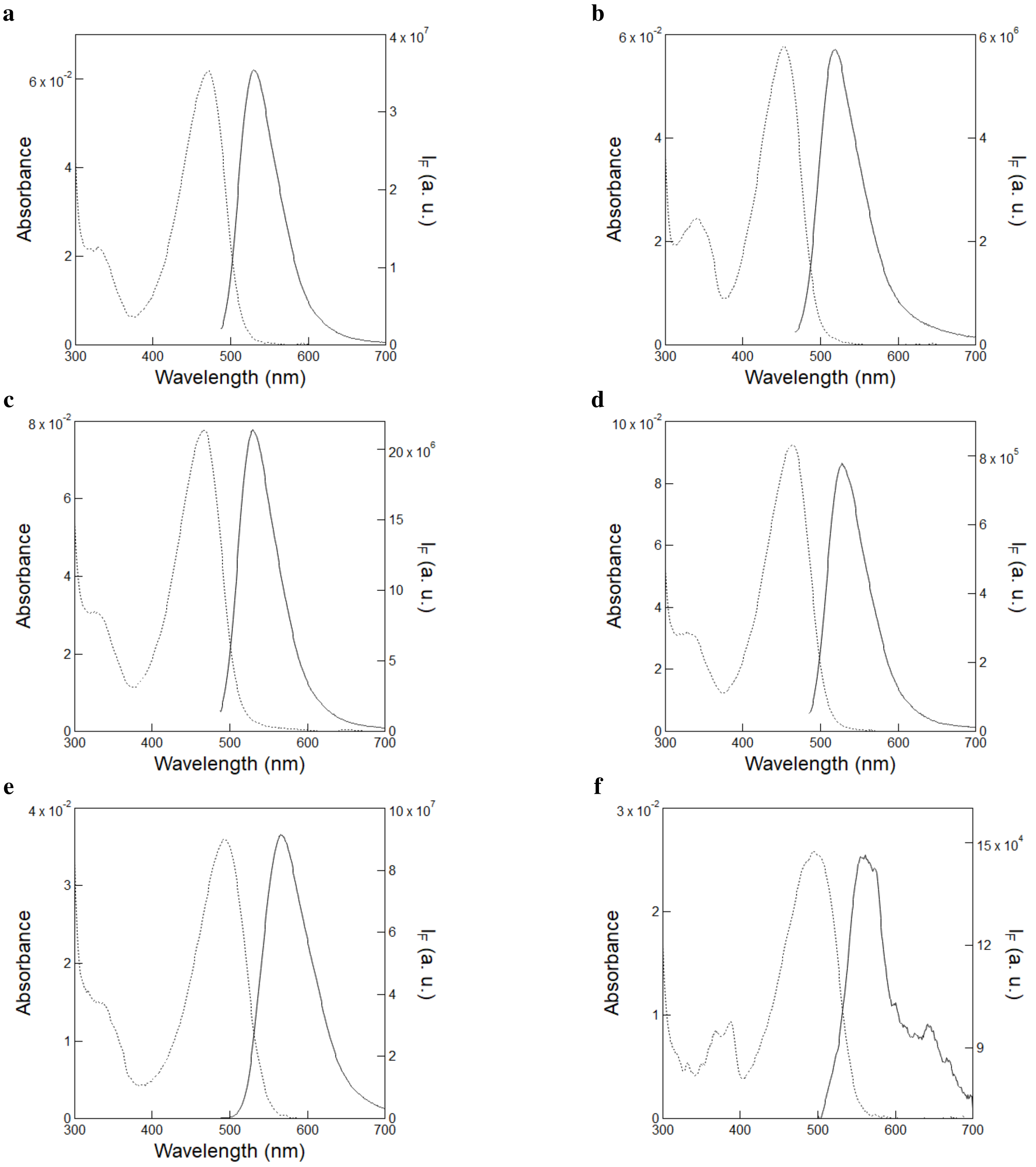

Figure S22: *UV-Vis absorption and emission spectra of the complexes between* ***pFAST*** *and the fluorogens.* The absorption (dotted line) and emission (solid line; $\lambda_{exc}$ = 480 nm) spectra with **HBR3Cl (a)**, **HBR3CN (b)**, **HBR3Cl5F (c)**, **HBR35DF (d)**, **HBR3F5OM (e)** and **HBIR3Cl (f)** were recorded at 5 $\mu$M (for **HBR3Cl**, **HBR3CN**, **HBR3Cl5F**, **HBR35DF**), 4 $\mu$M (**HBR3F5OM**), and 3 $\mu$M (**HBIR3Cl**) in the presence of 32 $\mu$M **pFAST** in 1× pH = 7.4 PBS buffer. $T$ = 293 K.

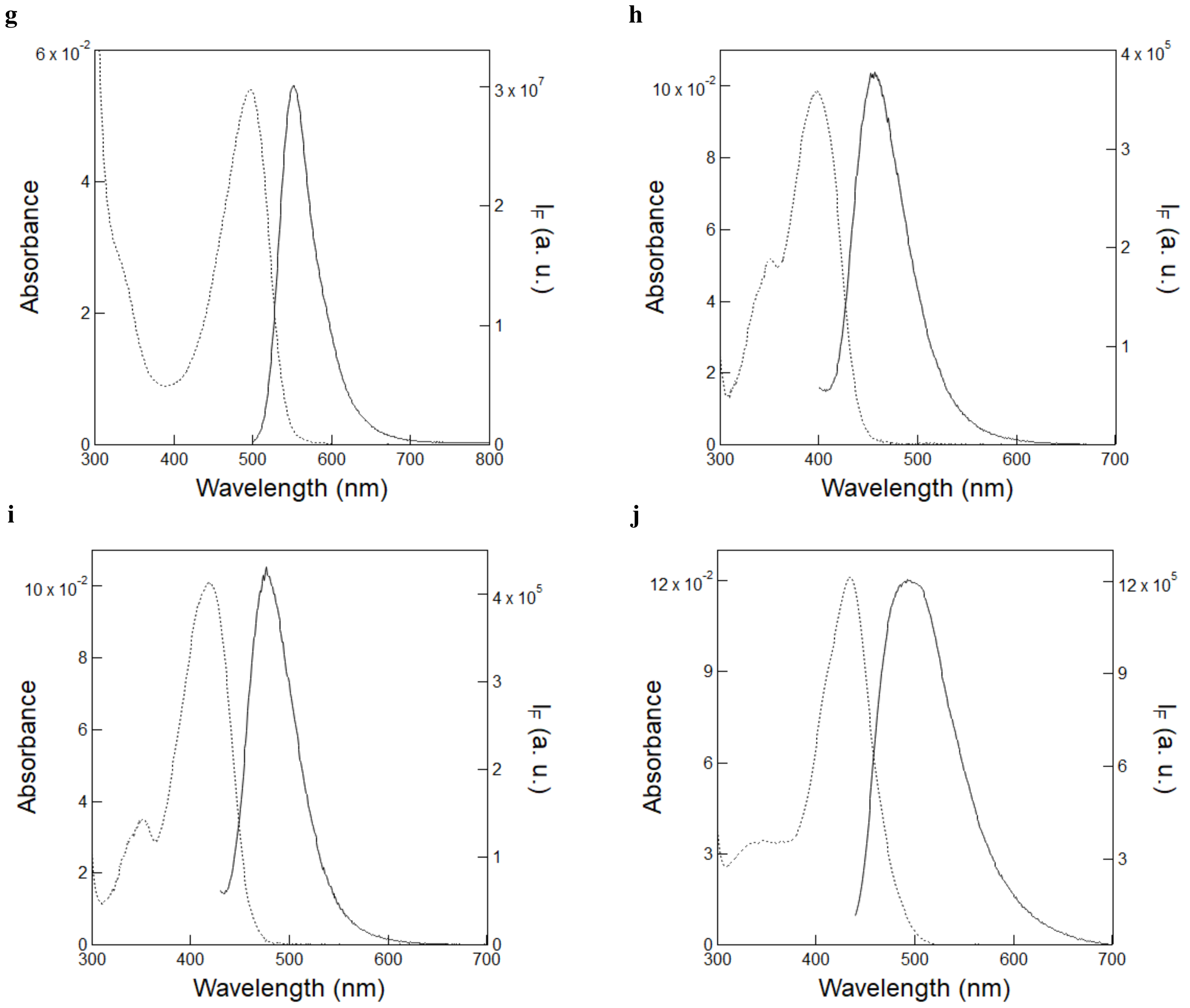

Figure S22: *UV-Vis absorption and emission spectra of the complexes between* ***pFAST*** *and the fluorogens.* The absorption (dotted line) and emission (solid line; $\lambda_{exc}$ = 480 nm (**HBR25DM**), $\lambda_{exc}$ = 380 nm (**HBT3CN**) and $\lambda_{exc}$ = 420 nm (**HBP3CN**, **HBT3CN**)) spectra with **HBR25DM (g)**, **HBT3CN (h)**, **HBP3CN (i)**, and **HBTH3CN (j)** were recorded at 4 $\mu$M **HBR25DM**, 10 $\mu$M **HBT3CN**, **HBP3CN**, and **HBTH3CN**, in the presence of 32 $\mu$M **pFAST** in 1$\times$ pH = 7.4 PBS buffer. $T$ = 293 K.

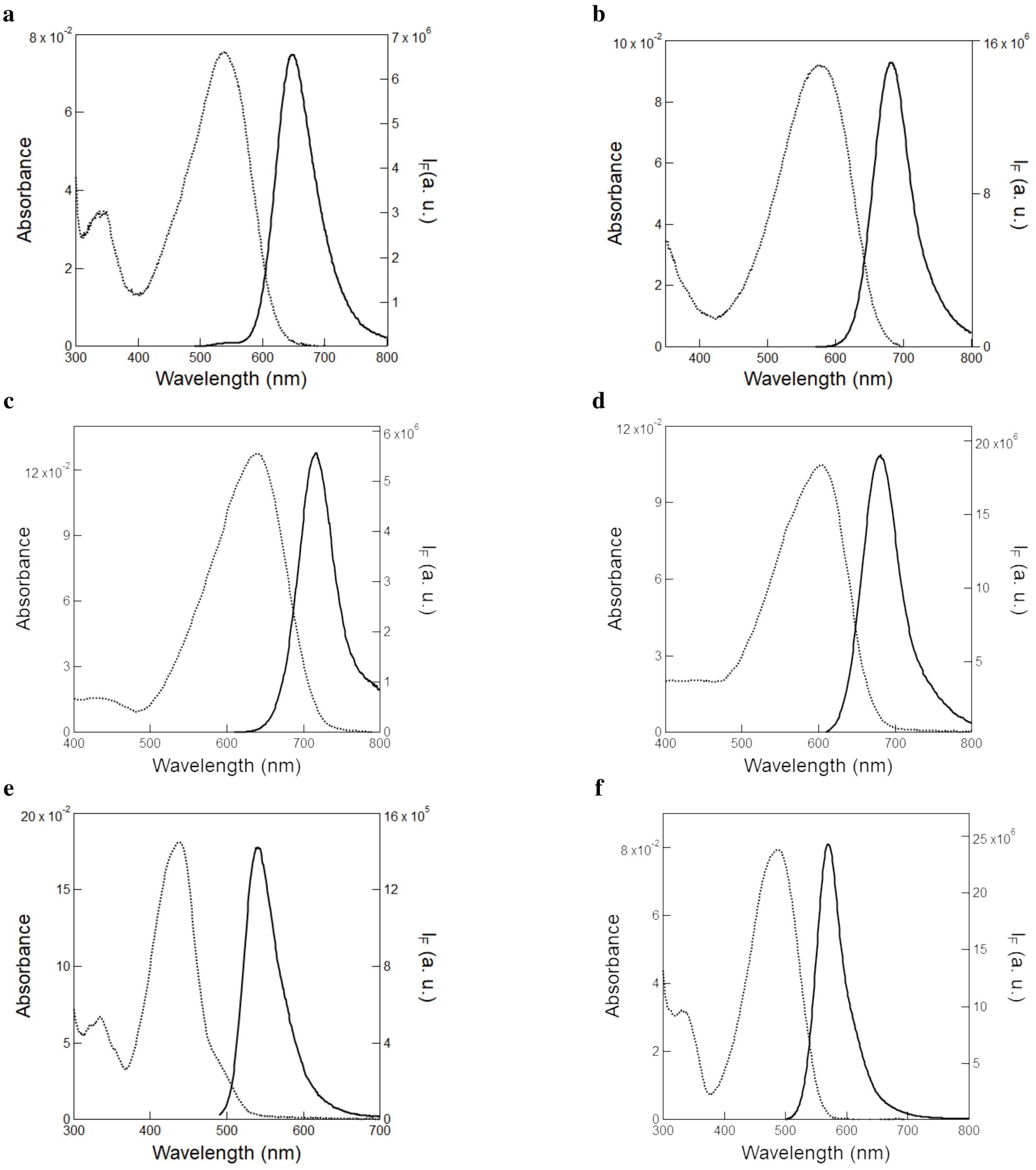

Figure S23: *UV-Vis absorption and emission spectra of the complexes between* ***nirFAST*** *and the fluorogens.* The absorption (dotted line) and emission (solid line) spectra with **HPAR3Cl (a)**, **HPAR3F5OM (b)**, **HPAR3OM (c)**, **HPAR35DOM (d)**, **HBR3Cl (e)** and **HBR3F5OM (f)** ($\lambda_{exc}$ = 480 nm for **(a)**, **(e)** and **(f)**, $\lambda_{exc}$ = 560 nm for **(b)** and $\lambda_{exc}$ = 600 nm for **(c)** and **(d)**) were recorded at 10 $\mu$M of fluorogens in the presence of 30 $\mu$M **nirFAST** in 1$\times$ pH = 7.4 PBS buffer. $T$ = 293 K.

**g**

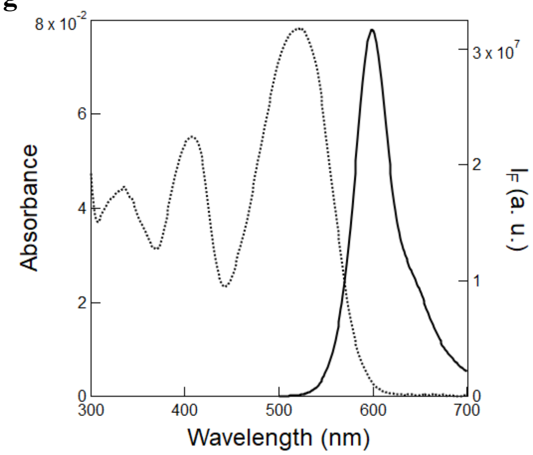

Figure S23: *UV-Vis absorption and emission spectra of the complexes between* ***nirFAST*** *and the fluorogens.* The absorption (dotted line) and emission (solid line) spectra with **HBR35DOM (g)** ($\lambda_{exc}$ = 480 nm) were recorded at 10 $\mu$M of fluorogen in the presence of 30 $\mu$M **nirFAST** in 1$\times$ pH = 7.4 PBS buffer. $T$ = 293 K.

Table S3: *Photophysical properties of the complex between **pFAST** and the (Z)-stereoisomer of the fluorogens.* Solvent: 1× pH = 7.4 PBS buffer. $T$ = 293 K.

| Fluorogens | $\lambda_{abs}^{max}$ (nm) | $\lambda_{em}^{max}$ (nm) | $\varepsilon$ ($mM^{-1}cm^{-1}$) | $\Phi$ | Brightness ($M^{-1}cm^{-1}$) |
|---|---|---|---|---|---|
| **HBT3CN** | 398 | 455 | 33 | – | – |
| **HBO3M** | 392 | 473 | 14 | 0.076 | 1000 |
| **HBP3CN** | 420 | 478 | 34 | – | – |
| **HBO35DM** | 411 | 483 | 6 | 0.23 | 1300 |
| **HBT35DM** | 433 | 499 | 33 | 0.10 | 3300 |
| **HBTH3CN** | 432 | 500 | 39 | – | – |
| **HBR3CN** | 454 | 516 | 44 | 0.02 | 880 |
| **HBR35DF** | 464 | 528 | 56 | 0.06 | 3360 |
| **HBR3Cl** | 471 | 530 | 41 | 0.08 | 3280 |
| **HBR3Cl5F** | 467 | 530 | 51 | 0.05 | 2550 |
| **HBT35DOM** | 449 | 532 | 60 | 0.27 | 16200 |
| **HBR3M** | 481 | 542 | 54 | 0.23 | 12400 |
| **HBR25DM** | 498 | 552 | 45 | 0.29 | 13000 |
| **HBP35DOM** | 487 | 554 | 37 | 0.33 | 12200 |
| **HBIR3Cl** | 497 | 557 | 29 | 0.002 | 50 |
| **HBR3F5OM** | 494 | 567 | 30 | 0.42 | 12600 |
| **HBR35DOM** | 520 | 600 | 44 | 0.35 | 15000 |
| **HBIR35DOM** | 562 | 616 | 40 | 0.10 | 4000 |

Abbreviations are as follows: $\lambda_{abs}^{max}$ and $\lambda_{em}^{max}$: wavelengths of maximum absorption and emission respectively; $\varepsilon$: molar absorption coefficient at $\lambda_{abs}^{max}$; $\Phi$: fluorescence quantum yield (see subsection 2.5.4 for details. Standard error: 10 %); Molecular brightness = $\Phi \times \varepsilon$. The photophysical data for **HBR3M**, **HBR25DM**, **HBR35DOM**, **HBIR35DOM**, **HBT35DM**, **HBT35DOM**, **HBO3M**, **HBO35DM** and **HBP35DOM** with **pFAST** were taken from.[4]

Table S4: *Photophysical properties of the complex between **nirFAST** and the (Z)-stereoisomer of the fluorogens.* Solvent: 1× pH = 7.4 PBS buffer. $T$ = 293 K.

| Fluorogens | $\lambda_{abs}^{max}$ (nm) | $\lambda_{em}^{max}$ (nm) | $\epsilon$ ($mM^{-1}cm^{-1}$) | $\Phi$ | Brightness ($M^{-1}cm^{-1}$) |
|---|---|---|---|---|---|
| **HBR3Cl** | 439 | 540 | 60 | – | – |
| **HBR3F5OM** | 487 | 570 | 26 | 0.15 | 4000 |
| **HBR35DOM** | 521 | 600 | 26 | 0.24* | 6240 |
| **HPAR3Cl** | 540 | 650 | 24 | 0.08 | 1920 |
| **HPAR3F5OM** | 580 | 680 | 30 | 0.14 | 4200 |
| **HPAR3OM** | 603 | 681 | 35 | 0.20* | 7000 |
| **HPAR35DOM** | 639 | 716 | 43 | 0.069* | 2970 |

Abbreviations are as follows: $\lambda_{abs}^{max}$ and $\lambda_{em}^{max}$: wavelengths of maximum absorption and emission respectively; $\varepsilon$: molar absorption coefficient at $\lambda_{abs}^{max}$; $\Phi$: fluorescence quantum yield (see subsection 2.5.4 for details. Standard error: 10 %); Molecular brightness = $\Phi \times \varepsilon$. *The data was taken from.[5]

##### 4.2 Measurement of the thermodynamic dissociation constant of the complex between pFAST or nirFAST and the thermodynamically stable state of the fluorogen ($K_d$)

###### 4.2.1 Complexes with pFAST

**4.2.1.1 HBR3Cl, HBR3Cl5F, HBR35DF, HBR3M, HBR25DM** 2.5 mL of 10 nM pFAST solution in 1× pH = 7.4 PBS buffer in a quartz cuvette (1×1 $cm^2$) were subjected to irradiation at 480 nm ($I = 1.1 \times 10^{-3}$ $E.m^{-2}.s^{-1}$) up to the photostationary state. The cuvette content was then kept in the dark for 5–10 min to get thermal recovery of the photoisomerized fluorogen before sequentially adding aliquots of 5 $\mu$L of 100 nM fluorogen and 10 nM pFAST in 1× pH = 7.4 PBS buffer and irradiating again the resulting solution with 480 nm light to reach the photostationary state.
We then subsequently fitted the $F_{tot}$-dependence of the fluorescence intensity $I_F$ at initial and final times of the investigated time window by using Eq. (S10).
Figure S24a–j displays the results for **HBR3Cl**, **HBR3Cl5F**, **HBR35DF**, **HBR3M** and **HBR25DM**. We retrieved for $K_d$ $0.09 \pm 0.01$ and $0.10 \pm 0.01$ nM (for **HBR3Cl**), $0.12 \pm 0.01$ and $0.16 \pm 0.01$ nM(for **HBR3Cl5F**), $0.14 \pm 0.01$ and $0.13 \pm 0.01$ nM (for **HBR35DF**), $3.00 \pm 0.20$ and $3.30 \pm 0.30$ (for **HBR3M**) and $0.80 \pm 0.05$ and $0.90 \pm 0.05$ nM (for **HBR25DM**) from processing the fluorogen concentration dependence of the fluorescence signal at initial and photostationary state.[c]

**4.2.1.2 HBR3F5OM, HBR35DOM, HBT35DM** Upon applying the same protocol on **HBR3F5OM**, **HBR35DOM** and **HBT35DM** (Figure S25a,c,e), we did not observe any time evolution of the fluorescence signal. Hence, we reduced analysis to processing the fluorogen concentration dependence of the observed fluorescence signal and retrieved $0.70 \pm 0.04$ (**HBR3F5OM**), $7.70 \pm 0.50$ (**HBR35DOM**) and $14.00 \pm 1.50$ (**HBT35DM**) nM for $K_d$.[d]

[c]We additionally extracted the total protein concentration from data fitting: 0.7, 0.9, 1.6, 4.6 and 5.0 nM were found for **HBR3Cl**, **HBR3Cl5F**, **HBR35DF**, **HBR3M** and **HBR25DM** respectively. Such values are lower than the nominal 10 nM concentration of protein, which could be attributed to nonspecific adsorption of the protein onto the glass surface.

[d]We additionally extracted the total protein concentration from data fitting: 7.8 (**HBR3F5OM**), 1.3 (**HBR35DOM**) and 5.7 nM (**HBR3F5OM**) was found, which again suggests nonspecific adsorption of the protein to occur onto the glass surface.

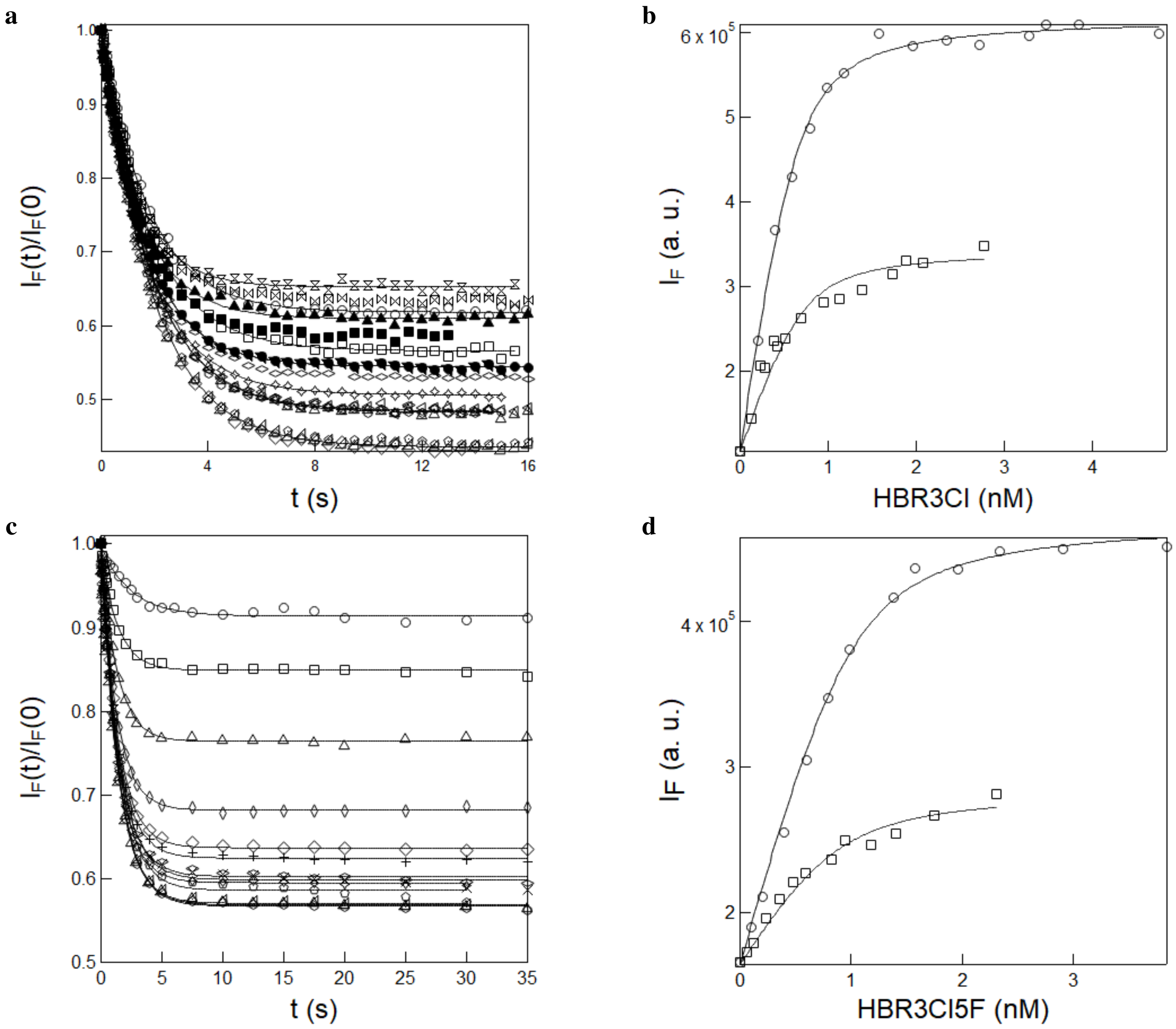

Figure S24: *Measurement of the thermodynamic dissociation constant of the complex between* ***pFAST*** *and the (Z)-stereoisomer of* ***HBR3Cl*** *and* ***HBR3Cl5F***. **a, c**: Time evolution of fluorescence emission at 530 nm from 10 nM **pFAST** solution in $1\times$ pH = 7.4 PBS buffer after successive addition of aliquots of 5 $\mu$L of 100 nM **HBR3Cl (a)** or 100 nM **HBR3Cl5F (c)** and 10 nM **pFAST** in $1\times$ pH = 7.4 PBS buffer upon constant illumination at 480 nm ($I = 1.1 \times 10^{-3}$ E.m$^{-2}$.s$^{-1}$) up to the photostationary state. Markers: experimental data; solid line: Monoexponential fit; **b, d**: Dependence of fluorescence intensity at initial (circles) and photostationary state (squares) on the concentration of **HBR3Cl (b)** and **HBR3Cl5F (d)** retrieved from **a** and **c** respectively. Markers: experimental data; solid line: Fit with Eq. (S10). $T = 293$ K.

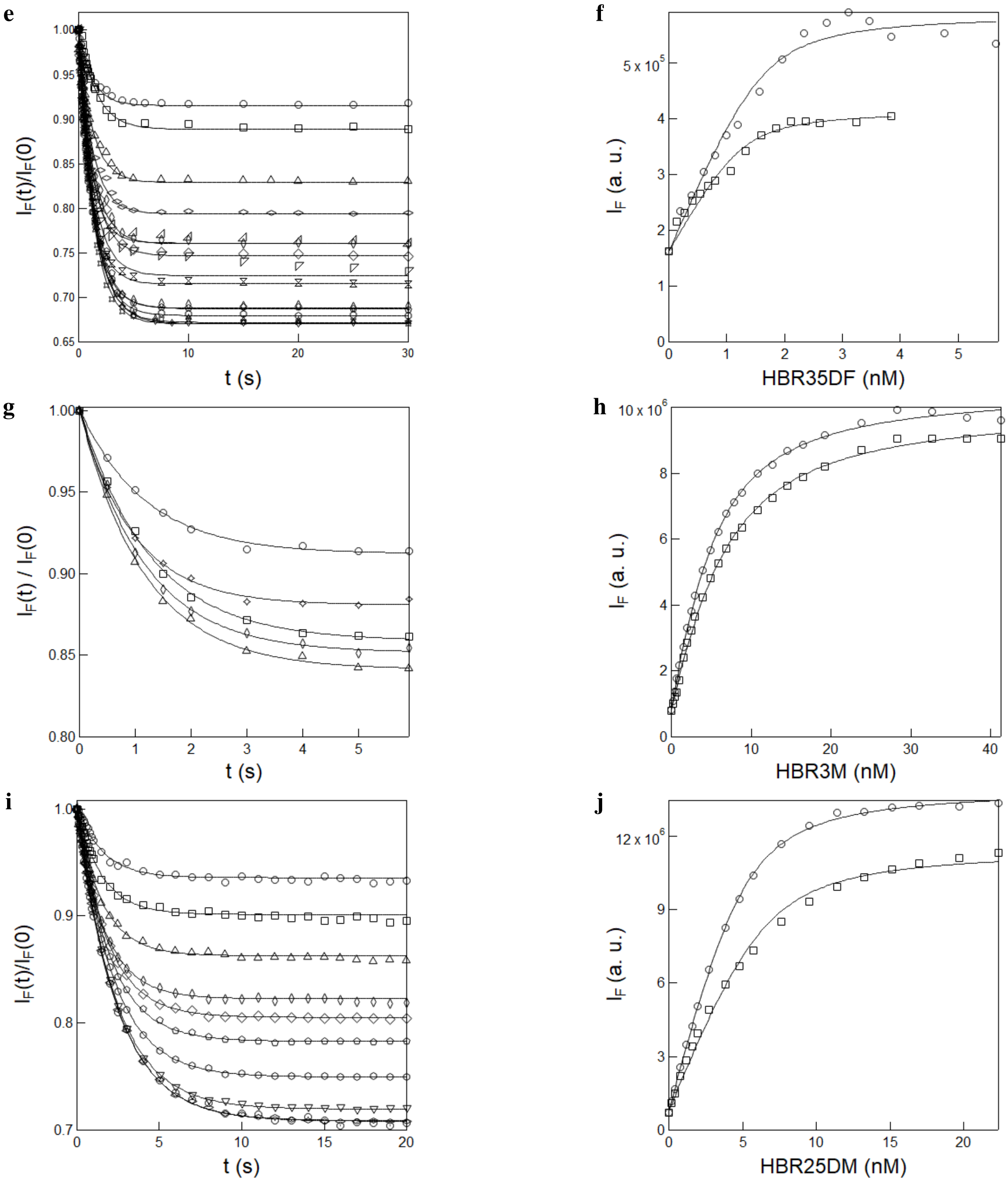

Figure S24: *Measurement of the thermodynamic dissociation constant of the complex between* ***pFAST*** *and the (Z)-stereoisomer of* ***HBR35DF****,* ***HBR3M*** *and* ***HBR25DM****.* **e,g,i**: Time evolution of fluorescence emission at 530 nm from 10 nM **pFAST** solution in $1\times$ pH = 7.4 PBS buffer after successive addition of aliquots of 5 $\mu$L of 100 nM **HBR35DF**, 500 nM **HBR3M** and **HBR25DM** containing 10 nM **pFAST** in $1\times$ pH = 7.4 PBS buffer upon constant illumination at 480 nm ($I = 1.1 \times 10^{-3}$ E.m$^{-2}$.s$^{-1}$) up to the photostationary state. Markers: experimental data; solid line: Monoexponential fit; **f**: Dependence of fluorescence intensity at initial (circles) and photostationary state (squares) on the concentration of **HBR35DF (f)**, **HBR3M (h)** and **HBR25DM (j)** retrieved from **e,g,i**. Markers: experimental data; solid line: Fit with Eq. (S10). $T = 293$ K.

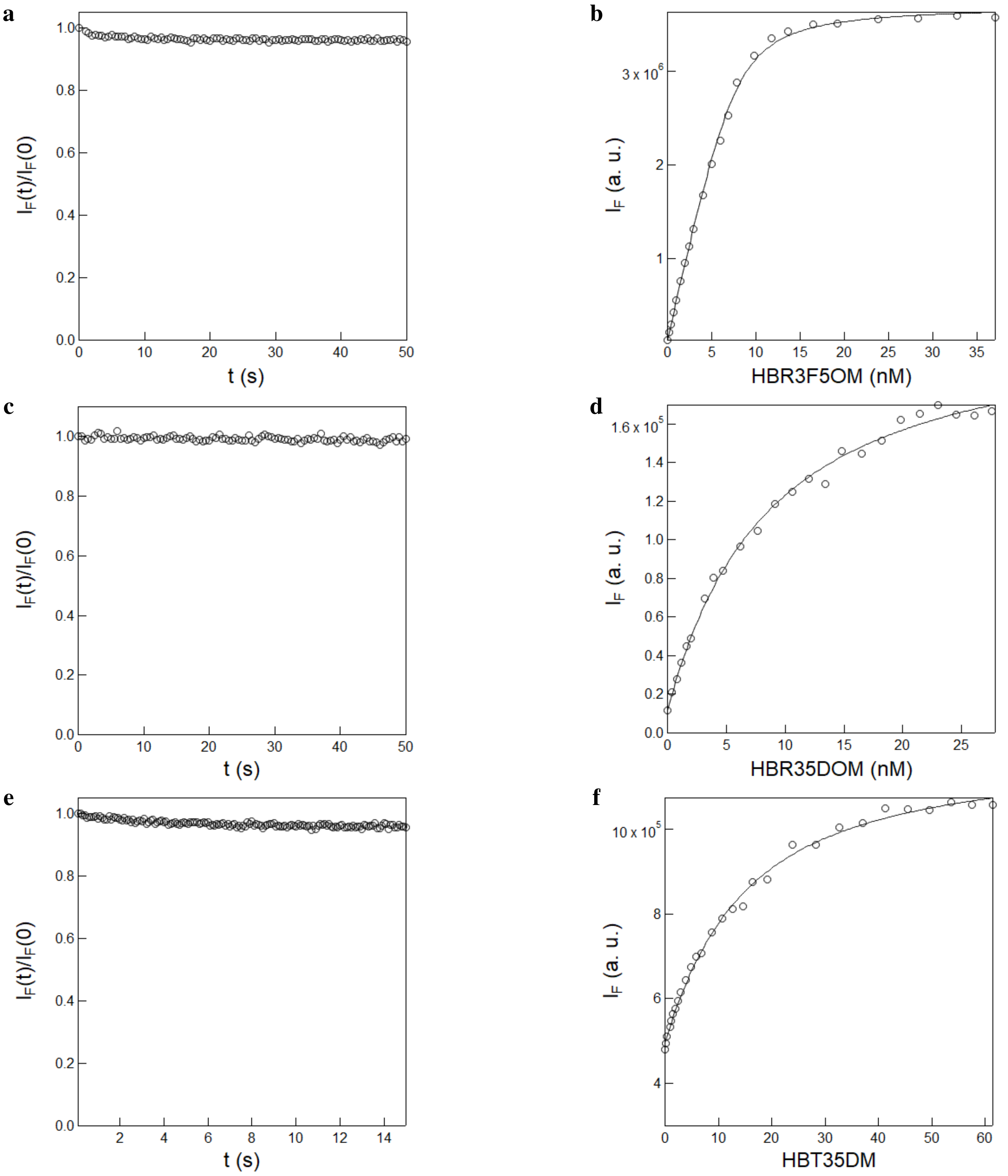

Figure S25: *Measurement of the thermodynamic dissociation constant of the complex between* ***pFAST*** *and the (Z)-stereoisomer of* ***HBR3F5OM****,* ***HBR35DOM*** *and* ***HBT35DM****.* **a,c,e**: Time evolution of fluorescence emission at 570 nm (for **HBR3F5OM**), 610 nm (**HBR35DOM**) and 500 nm (for **HBT35DM**) from 10 nM **pFAST** solution in 1× pH = 7.4 PBS buffer after addition of 5 $\mu$L of 500 nM **HBR3F5OM (a)**, **HBT35DM (e)** or 200 nM **HBR35DOM (c)** and 10 nM **pFAST** in 1× pH = 7.4 PBS buffer upon constant illumination at 480 nm ($I = 1.1 \times 10^{-3}$ E.m$^{-2}$.s$^{-1}$). Markers: experimental data; **b,d,f**: Dependence of the fluorescence intensity at initial state (circles) on the concentration of **HBR3F5OM (b)**, **HBR35DOM (d)** and **HBT35DM (f)**. Markers: experimental data; solid line: Fit with Eq. (S10). $T = 293$ K.

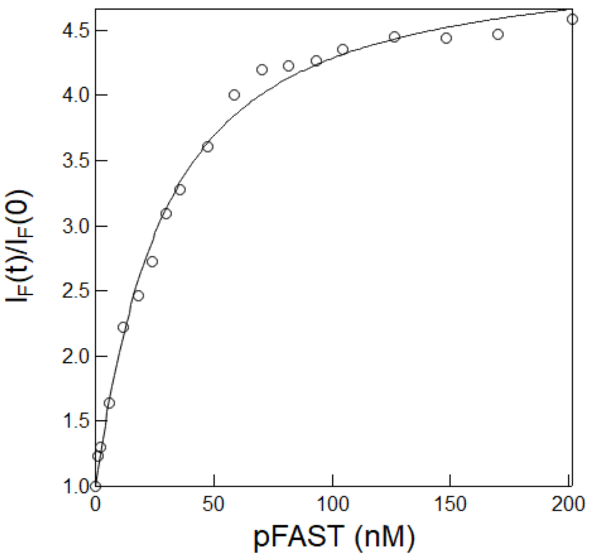

Figure S26: *Measurement of the thermodynamic dissociation constant of the complex between* ***pFAST*** *and the (Z)-stereoisomer of* ***HBR3CN***. Fluorogen concentration dependence of the initial fluorescence signal at 530 nm from titrating 10 nM **HBR3CN** solution in 1× pH = 7.4 PBS buffer after successive addition of aliquots of 3000 nM **pFAST** and 10 nM **HBR3CN** solution in 1× pH = 7.4 PBS buffer upon excitation using 480 nm Xe lamp. Markers: experimental data; solid line: Fit with Eq. (S10). $T = 293$ K.

**4.2.1.3 HBR3CN** The protocol described above has been modified for extracting $K_d$ with **HBR3CN**. Indeed, thermal recovery was too slow for **HBR3CN**. Hence, we performed a regular titration by using the Xenon lamp of our fluorimeter for reading out fluorescence without generating any significant photoisomerization due to its low intensity (Figure S26). Hence, we processed the protein concentration-dependence of the fluorescence signal and retrieved $22.7 \pm 1.50$ (for **HBR3CN**) for $K_d$.[e]

[e] We additionally retrieved from fitting 10 nM for the total fluorogen concentration with **HBR3CN**.

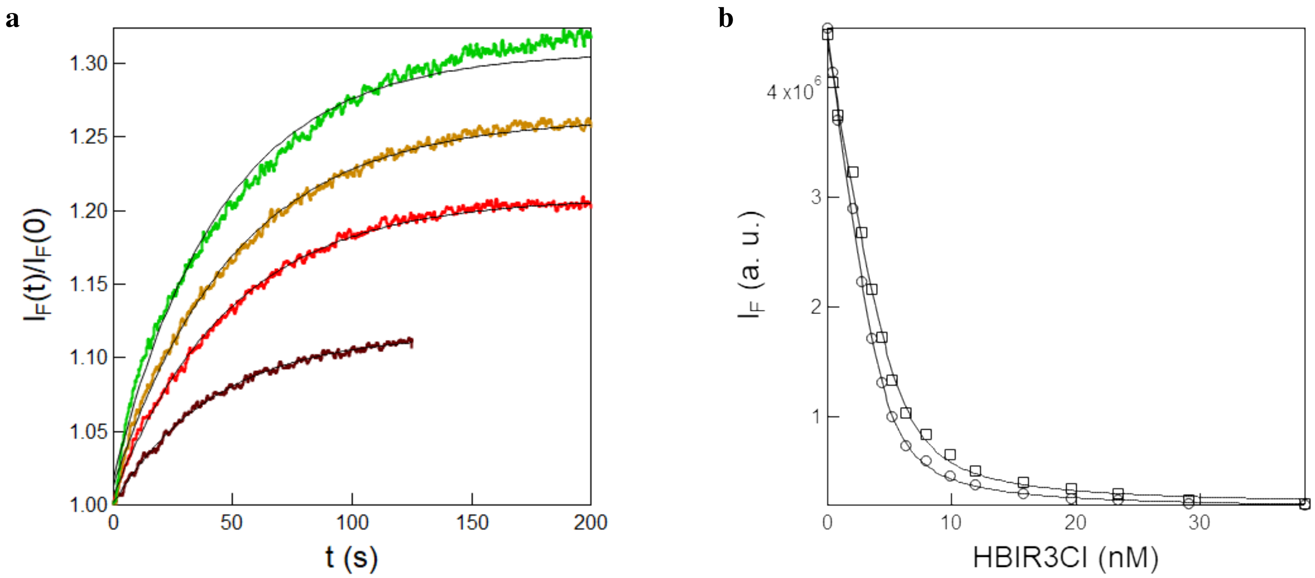

Figure S27: *Measurement of the thermodynamic dissociation constant of the complex between **pFAST** and the (Z)-stereoisomer of **HBIR3Cl**.* **a**: Time evolution of fluorescence emission at 610 nm from 10 nM **HBR35DOM** and **pFAST** solution in 1× pH = 7.4 PBS buffer after successive addition of aliquots of 5 $\mu$L of 1000 nM **HBIR3Cl** and 10 nM **pFAST** in 1× pH = 7.4 PBS buffer upon constant illumination at 480 nm ($I = 1.1 \times 10^{-3}$ E.m$^{-2}$.s$^{-1}$) up to the photostationary state. Markers: experimental data; solid line: Monoexponential fit; **b**: Dependence of fluorescence intensity at initial (circles) and photostationary state (squares) on the concentration of **HBIR3Cl** retrieved from **a**. Markers: experimental data; solid line: Fit with Eq.(S15). $T = 293$ K.

**4.2.1.4 HBIR3Cl** **HBIR3Cl** exhibits negligible fluorescence upon binding with **pFAST**. Therefore, in order to measure its affinity for **pFAST**, we designed a competition binding experiment by using the **HBR35DOM** nonphotoisomerizing fluorogen as a competitor.

Here, the titration experiments were performed by titrating 2.5 mL of 10 nM **HBR35DOM** and **pFAST** solution in 1× pH = 7.4 PBS buffer contained in a 3 mL quartz cuvette (1 cm optical path length) with 1000 nM **HBIR3Cl** and 10 nM **pFAST** solution in 1× pH = 7.4 PBS buffer.

After addition of each aliquot, the solution was subjected to irradiation at 480 nm ($I = 1.1 \times 10^{-3}$ E.m$^{-2}$.s$^{-1}$) up to the photostationary state (Figure S27a). The cuvette content was then kept in the dark for 5–10 min to get thermal recovery of the photoisomerized **HBIR3Cl** before proceeding to the next aliquot addition of **HBIR3Cl** and **pFAST** solution.

We retrieved $K_d = 0.40 \pm 0.04$ nM from processing the fluorogen concentration dependence of the fluorescence signal at both initial and photostationary state with Eq.(S15) (Figure S27b).[f]

[f] We additionally extracted the total **pFAST-HBR35DOM** concentration from data fitting: 4.5 nM was found. Such value is lower than the nominal 10 nM concentration of initial complex, which could be attributed to its nonspecific adsorption onto the glass surface.

##### 4.2.2 Complexes with nirFAST

**4.2.2.1 HPAR3Cl, HBR3F5OM** 2.5 mL of 20 nM **nirFAST** solution in 1× pH = 7.4 PBS buffer in a quartz cuvette (1×1 $cm^2$) were subjected to irradiation at 480 nm or 640 nm ($I = 1.1 \times 10^{-3}$ $E.m^{-2}.s^{-1}$) up to the photostationary state. The cuvette content was then kept in the dark for 5–10 min to get thermal recovery of the photoisomerized fluorogen before sequentially adding aliquots of 5 $\mu$L of 500 nM fluorogen and 20 nM **nirFAST** in 1× pH = 7.4 PBS buffer and irradiating again the resulting solution with 480 nm or 640 nm light to reach the photostationary state.

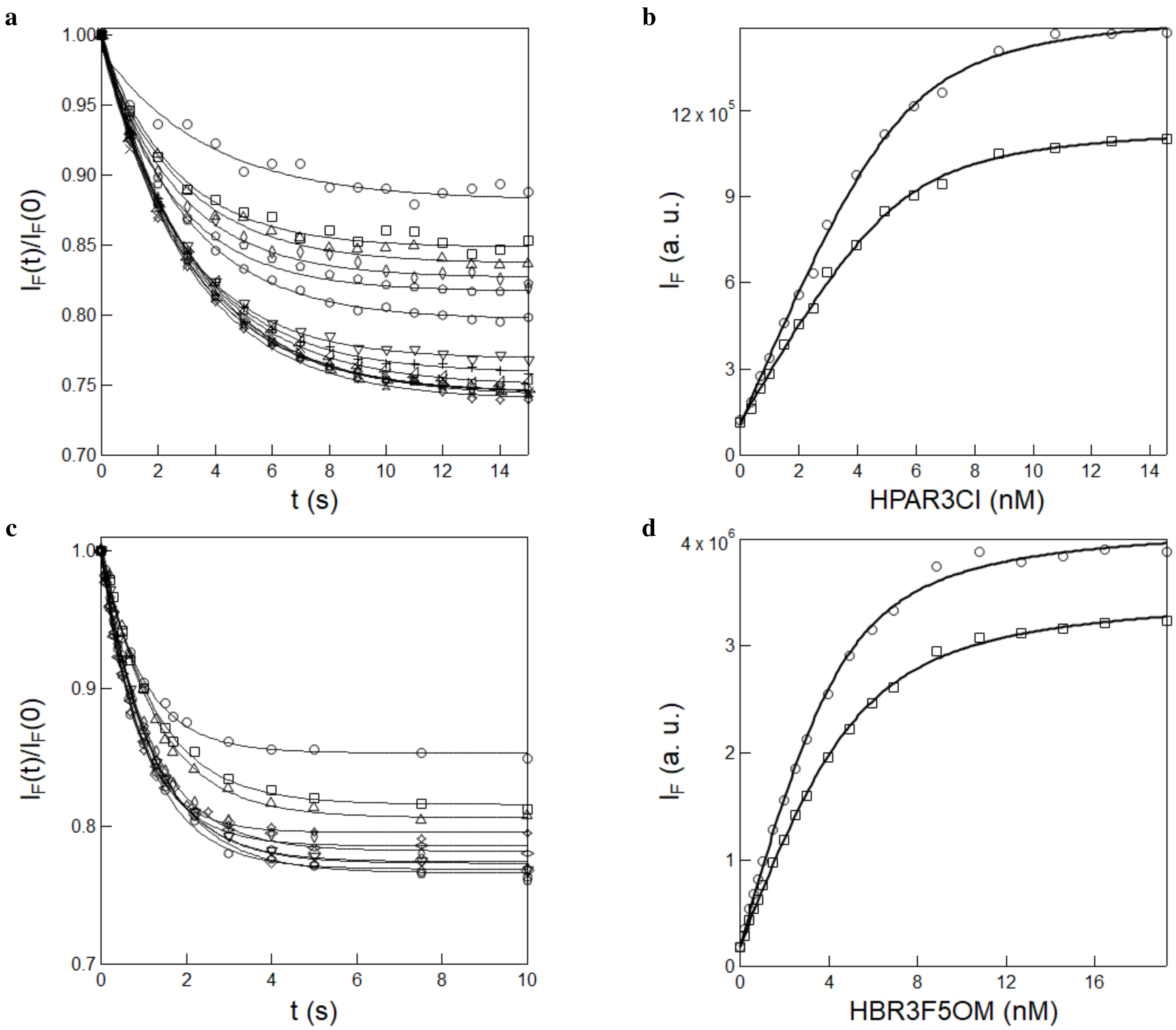

Figure S28: *Measurement of the thermodynamic dissociation constant of the complex between **nirFAST** and the (Z)-stereoisomer of **HPAR3Cl** and **HBR3F5OM***. **a,c**: Time evolution of fluorescence emission at 650 nm (for **HPAR3Cl**) and 570 nm (for **HBR3F5OM**) from 20 nM **nirFAST** solution in 1× pH = 7.4 PBS buffer after successive addition of aliquots of 5 $\mu$L of 500 nM **HPAR3Cl (a)** and **HBR3F5OM (c)** and 20 nM **nirFAST** in 1× pH = 7.4 PBS buffer upon constant illumination at 480 nm ($I = 1.1 \times 10^{-3}$ $E.m^{-2}.s^{-1}$) up to the photostationary state. Markers: experimental data; solid line: Monoexponential fit; **b,d**: Dependence of fluorescence intensity at initial (circles) and photostationary state (squares) on the concentration of **HPAR3Cl (b)** and **HBR3F5OM (d)**. Markers: experimental data; solid line: Fit with Eq. (S10). $T = 293$ K.

Figure S28 display the results for **HPAR3Cl** and **HBR3F5OM**. We retrieved for $K_d$ $0.52 \pm 0.07$ and $0.54 \pm 0.09$ nM for **HPAR3Cl** and $0.90 \pm 0.07$ and $1.10 \pm 0.09$ nM (for **HBR3F5OM**) from processing the fluorogen concentration dependence of the fluorescence signal at initial and photostationary state.[g]

**4.2.2.2 HBR3Cl, HBR35DOM, HPAR3F5OM, HPAR3OM, HPAR35DOM** Upon applying the same protocol on **HPAR3F5OM**, **HBR35DOM**, **HBR3Cl**, **HPAR3OM**, and **HPAR35DOM** (Figure S29a,c,e.g), we did not observe any time evolution of the fluorescence signal. Hence, we reduced analysis to processing the fluorogen concentration dependence of the observed fluorescence signal and retrieved $0.70 \pm 0.03$ (**HPAR3F5OM**), $6.70 \pm 0.40$ (**HBR35DOM**), $0.14 \pm 0.03$ (**HBR3Cl**), $0.70 \pm 0.06$ (**HPAR3OM**), and $1.6 \pm 0.20$ (**HPAR35DOM**) nM for $K_d$.[h]

**4.2.2.3 HBIR3Cl** **HBIR3Cl** exhibits negligible fluorescence upon binding with **nirFAST**. Therefore, we applied the protocol displayed in Figure S27 in order to measure its affinity for **nirFAST** by using the **HBR35DOM** nonphotoisomerizing fluorogen as a competitor.
We retrieved $K_d = 1.50 \pm 0.30$ nM from processing the fluorogen concentration dependence of the fluorescence signal at both initial and photostationary state with Eq.(S15) (Figure S30b).[i]

[g] We additionally extracted the total protein concentration from data fitting: 5.4 and 4.7 nM was found for **HPAR3Cl** and **HBR3F5OM** respectively. Such values are lower than the nominal 20 nM concentration of protein, which could be attributed to nonspecific adsorption of the protein onto the glass surface.

[h] We additionally extracted the total protein concentration from data fitting: 10, 3.5, 6, 1.8 and 2.1 nM were found for **HPAR3F5OM**, **HBR35DOM**, **HBR3Cl**, **HPAR3OM**, and **HPAR35DOM** respectively, which again suggests nonspecific adsorption of the protein to occur onto the glass surface.

[i] We additionally extracted the total **nirFAST-HBR35DOM** concentration from data fitting: 6.0 nM was found. Such value is lower than the nominal 10 nM concentration of initial complex, which could be attributed to its nonspecific adsorption onto the glass surface.

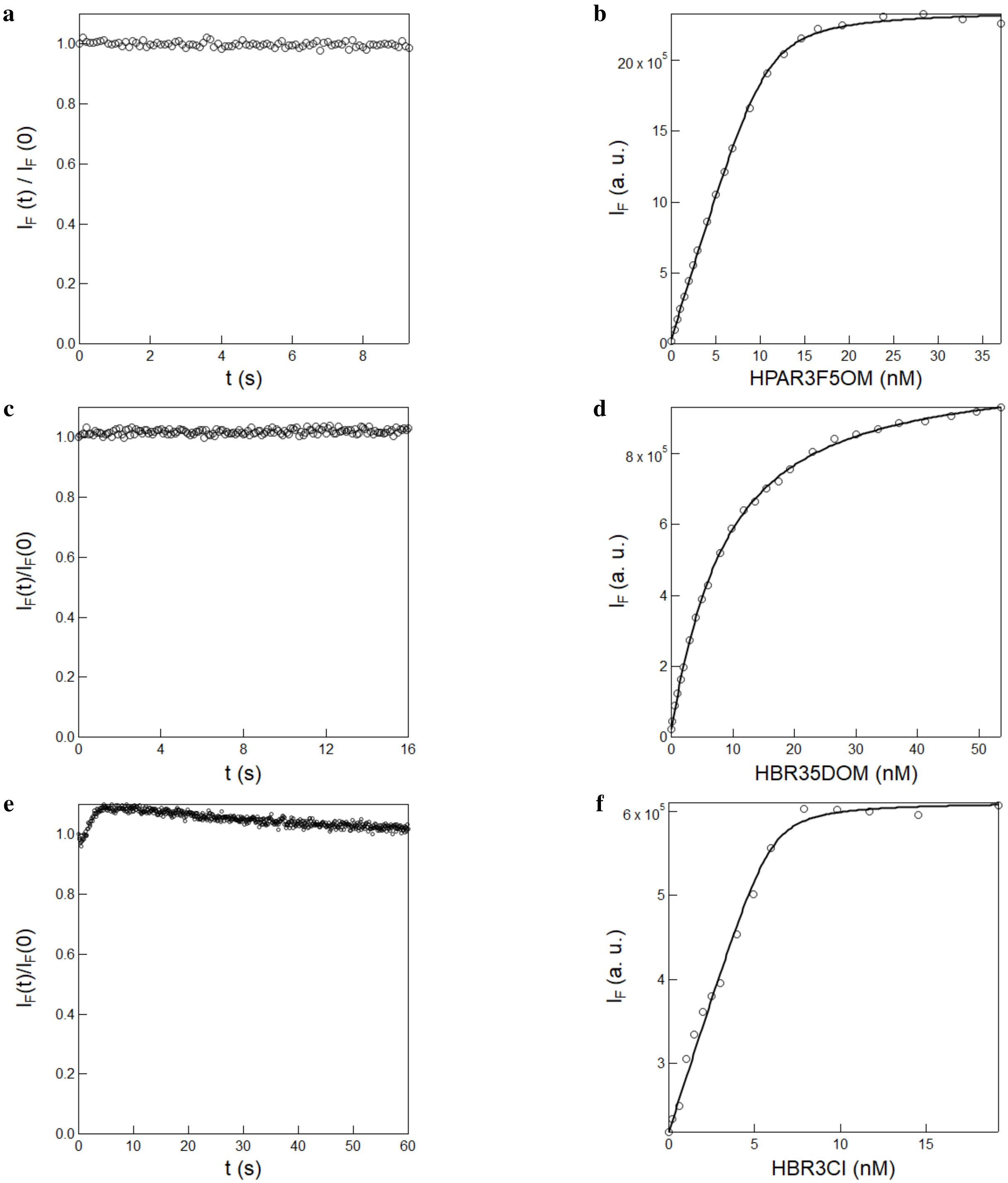

Figure S29: *Measurement of the thermodynamic dissociation constant of the complex between **nirFAST** and the (Z)-stereoisomer of **HPAR3F5OM**, **HBR35DOM**, and **HBR3Cl**.* **a, c, e**: Time evolution of fluorescence emission at 680 (**HPAR3F5OM**), 600 (**HBR35DOM**) and 540 (**HBR3Cl**) nm from 20 nM **nirFAST** solution in 1$\times$ pH = 7.4 PBS buffer after addition of aliquots of 100 $\mu$L of 500 nM **HPAR3F5OM (a)**, **HBR35DOM (c)**, or **HBR3Cl (e)** and 20 nM **nirFAST** in 1$\times$ pH = 7.4 PBS buffer upon constant illumination at 480 nm ($I = 1.1 \times 10^{-3}$ E.m$^{-2}$.s$^{-1}$). Markers: experimental data; **b, d**: Dependence of fluorescence intensity at initial state (circles) on the concentration of **HPAR3F5OM (b)**, **HBR35DOM (d)**, and **HBR3Cl (f)**. Markers: experimental data; solid line: Fit with Eq. (S10). $T = 293$ K.

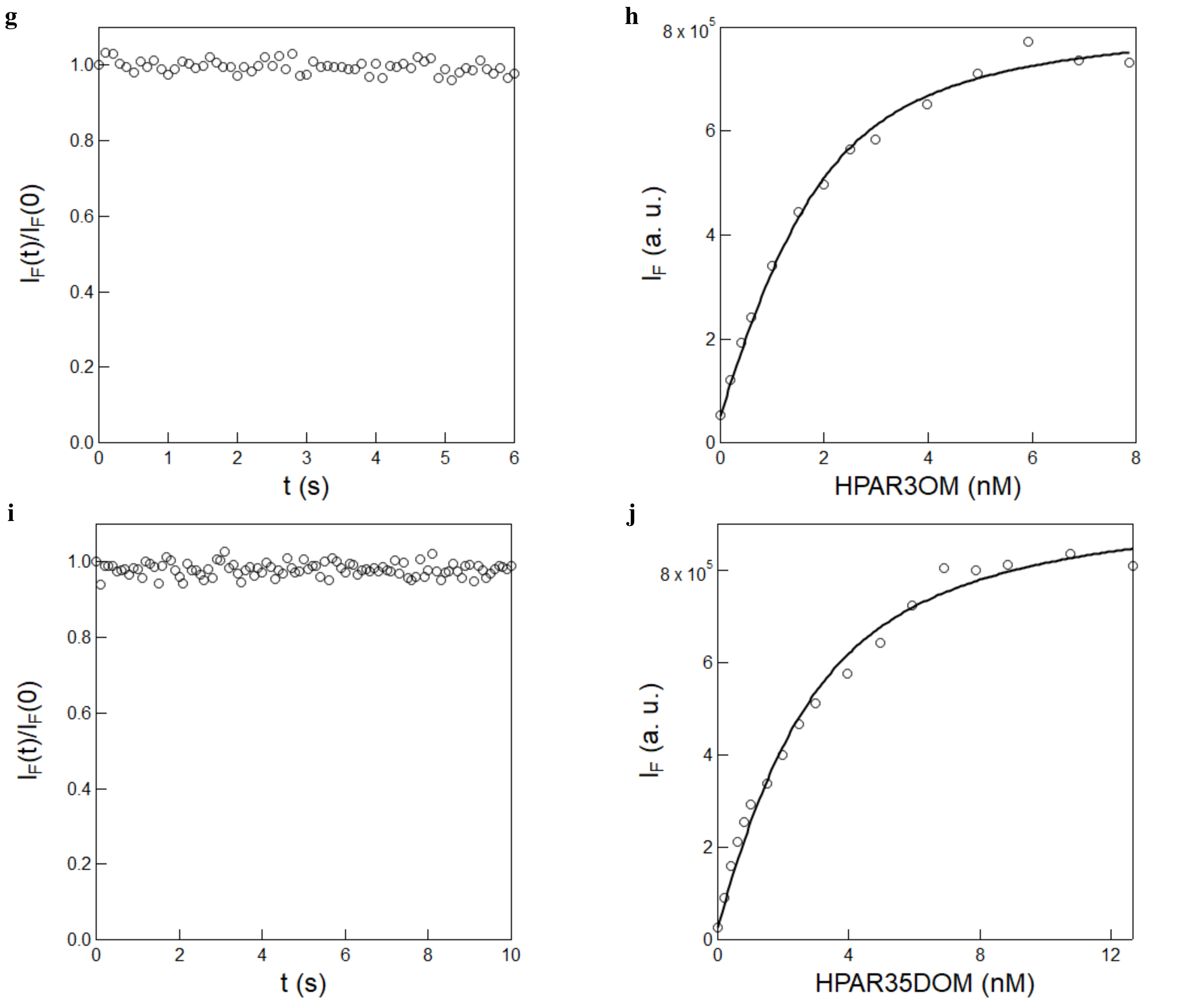

Figure S29: *Measurement of the thermodynamic dissociation constant of the complex between **nirFAST** and the (Z)-stereoisomer of **HPAR3OM** and **HPAR35DOM**.* **g,i**: Time evolution of fluorescence emission at 680 nm (**HPAR3OM**) and 715 nm (**HPAR35DOM**) from 20 nM **nirFAST** solution in 1× pH = 7.4 PBS buffer after addition of aliquots of 100 $\mu$L of 500 nM **HPAR3OM (g)**, **HPAR35DOM (i)** and 20 nM **nirFAST** in 1× pH = 7.4 PBS buffer upon constant illumination at 640 nm ($I = 1.1 \times 10^{-4}$ E.m$^{-2}$.s$^{-1}$). Markers: experimental data; **h,j**: Dependence of fluorescence intensity at initial state (circles) on the concentration of **HPAR3OM (h)** and **HPAR35DOM (j)**. Markers: experimental data; solid line: Fit with Eq. (S10). $T = 293$ K.

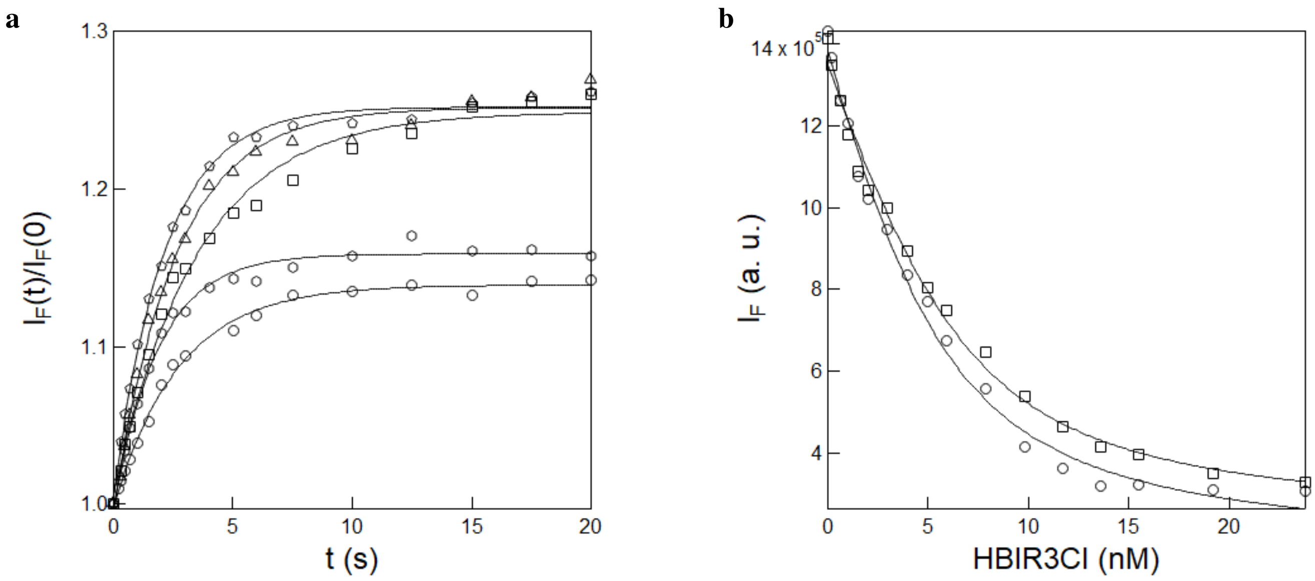

Figure S30: *Measurement of the thermodynamic dissociation constant of the complex between **nirFAST** and the (Z)-stereoisomer of **HBIR3Cl**.* **a**: Time evolution of fluorescence emission at 610 nm from 10 nM **HBR35DOM** and **nirFAST** solution in $1\times$ pH = 7.4 PBS buffer after successive addition of aliquots of 5 $\mu$L of 1000 nM **HBIR3Cl** and 10 nM **nirFAST** in $1\times$ pH = 7.4 PBS buffer upon constant illumination at 480 nm ($I = 1.1 \times 10^{-3}$ E.m$^{-2}$.s$^{-1}$) up to the photostationary state. Markers: experimental data; solid line: Monoexponential fit; **b**: Dependence of fluorescence intensity at initial (circles) and photostationary state (squares) on the concentration of **HBIR3Cl** retrieved from **a**. Markers: experimental data; solid line: Fit with Eq.(S15). $T = 293$ K.

Table S5: *Thermodynamic properties of the complex between* ***pFAST*** *and the (Z)-stereoisomer of the fluorogens.* Solvent: 1× pH = 7.4 PBS buffer. $T$ = 293 K.

| Fluorogens | $K_d$ (nM) |
|---|---|
| **HBR3Cl** | $0.10 \pm 0.01$ |
| **HBR3Cl5F** | $0.14 \pm 0.01$ |
| **HBR35DF** | $0.14 \pm 0.01$ |
| **HBIR3Cl** | $0.40 \pm 0.04$ |
| **HBR3F5OM** | $0.70 \pm 0.04$ |
| **HBR25DM** | $0.80 \pm 0.05$ |
| **HBR3M** | $3.0 \pm 0.2$ |
| **HBR35DOM** | $7.7 \pm 0.5$ |
| **HBT35DM** | $14.0 \pm 1.5$ |
| **HBR3CN** | $22.7 \pm 1.5$ |

Table S6: *Thermodynamic properties of the complex between* ***nirFAST*** *and the (Z)-stereoisomer of the fluorogens.* Solvent: 1× pH = 7.4 PBS buffer. $T$ = 293 K.

| Fluorogens | $K_d$ (nM) |
|---|---|
| **HBR3Cl** | $0.14 \pm 0.03$ |
| **HPAR3Cl** | $0.70 \pm 0.04$ |
| **HPAR3F5OM** | $0.70 \pm 0.03$ |
| **HPAR3OM** | $0.70 \pm 0.06$ |
| **HBR3F5OM** | $0.90 \pm 0.07$ |
| **HBIR3Cl** | $1.5 \pm 0.3$ |
| **HPAR35DOM** | $1.6 \pm 0.2$ |
| **HBR35DOM** | $6.7 \pm 0.4$ |

Abbreviation is as follows: $K_d$ designates the thermodynamic dissociation constant of the complex between **FAST** and the ($Z$)-stereoisomer of the fluorogens.

#### 4.3 Reference trajectory associated to replacing HBT35DM by HBR35DOM in pFAST in the CIE 1931 color space

To further appreciate the hue change generated from illuminating **pFAST** in the presence of a photoisomerizable fluorogen and a second fluorogen giving a **pFAST** complex of different color, we performed a titration experiment by recording the fluorescence emission spectrum from 700 nM **HBT35DM** and 100 nM **pFAST** in 1× pH = 7.4 PBS buffer upon adding aliquots of 10 $\mu$M **HBR35DOM** and 100 nM **pFAST** solution in the same solvent (Figure S31a). As anticipated, the fluorescence emission from the **pFAST:HBT35DM** complex is replaced by the one of the **pFAST:HBR35DOM** complex. As displayed in Figure S31b, the associated trajectory of the fluorescence emission hue in the CIE 1931 color space is linear and it essentially visits the whole distance between the two extrema associated with the **pFAST:HBT35DM** and **pFAST:HBR35DOM** complexes respectively.

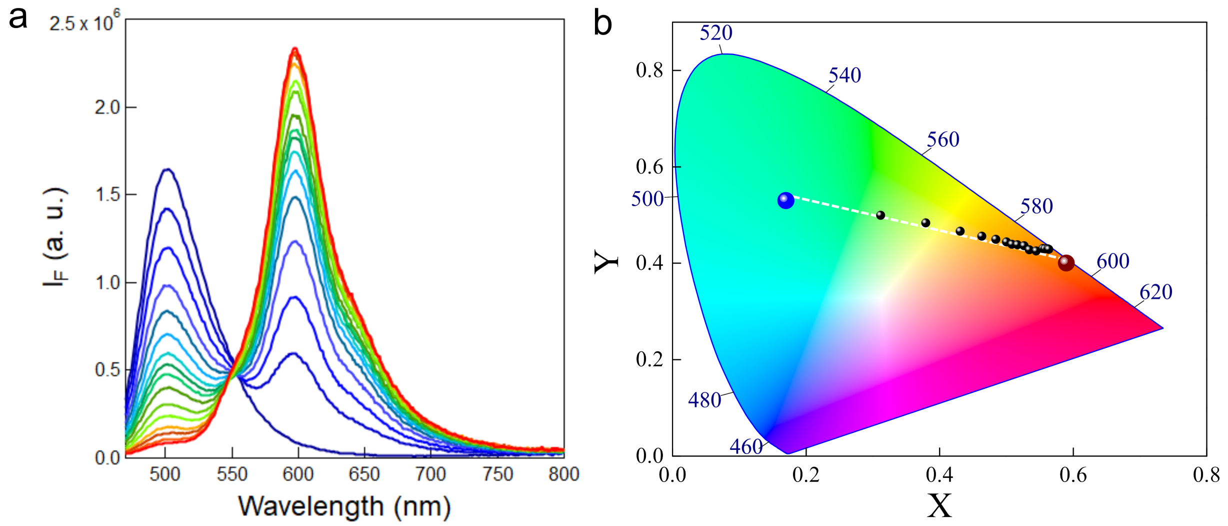

Figure S31: *Hue change originating from replacing a fluorogen by another one in a* ***pFAST*** *complex*. **a:** Evolution of the fluorescence emission spectra from 700 nM **HBT35DM** and 100 nM **pFAST** in 1× pH = 7.4 PBS buffer upon successive addition of aliquots of 10 $\mu$M **HBR35DOM** and 100 nM **pFAST** in 1× pH = 7.4 PBS buffer; **b:** Associated trajectory of the fluorescence emission hue in the CIE 1931 color space. The coordinates of the fluorescence emission from the pure complexes **pFAST:HBT35DM** and **pFAST:HBR35DOM** are displayed as blue and dark red circles respectively whereas the coordinates of the intermediate spectra obtained during the titration and aligned along a linear trajectory (dashed line) are displayed as black circles. $T = 298$ K.

### 5 Confocal microscopy experiments

#### 5.1 Experiments involving one fluorogen

##### 5.1.1 Photoejection of HBR25DM

As displayed in Figure S32, the fluorescence signal at the labeled nucleus decreased by up to 80 % upon repetitive scans in the presence of 0.1 $\mu$M of **HBR25DM** fluorogen with **pFAST**. Photoejection reversibility was evidenced by essentially recovering the initial fluorescence level upon introducing a sufficient delay (typically 120 s) before applying a new series of repetitive scans. Eventually, the fluorescence drop originating from photoejection could be abolished by increasing the fluorogen concentration above 1 $\mu$M, rejoining well established experimental conditions for **FAST** labeling.

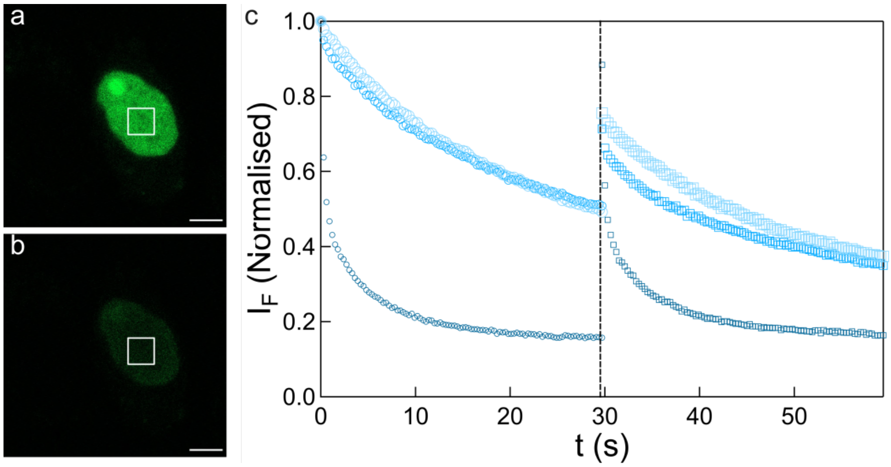

Figure S32: ***FAST** to generate a palette of negative ncRSFPs*. 488 nm light-induced fluorescence emission from **HBR25DM** (**a-c**) in **H2B-pFAST**-labeled living HeLa cells observed in confocal microscopy in the presence of the fluorogen in DMEM pH=7.4. Photoejection is evidenced at 0.1 $\mu$M concentration from the contrast at the nuclei between the initial (**a**) and photosteady state (**b**) images, its reversibility by the similar loss of fluorescence normalized by its initial value after two consecutive series of frame acquisition (circles and squares markers after the 1$^{st}$ and 2$^{nd}$ series respectively) in the region of interest at nuclei, and its concentration-dependence by the distinct decays observed at 0.1 (small black markers), 1 (medium grey markers), or 10 (large light grey markers) $\mu$M fluorogen concentration (**c**). Scale bar: 5 $\mu$m. $T = 298$ K. See Table S8 for the conditions of image acquisition.

##### 5.1.2 Photoejection of HPAR3Cl

As displayed in Figure S33, the fluorescence signal at the labeled nucleus decreased by up to 80 % upon repetitive scans in the presence of 0.1 $\mu$M of **HPAR3Cl** fluorogen with **nirFAST**. Photoejection reversibility was evidenced by essentially recovering the initial fluorescence level upon introducing a sufficient delay (typically 120 s) before applying a new series of repetitive scans.

#### 5.2 Experiments involving two fluorogens

##### 5.2.1 HPAR3OM:HBR35DOM

Figure S34 shows that 543 nm illumination of live HeLa cells expressing H2B-**nirFAST** conditioned with 0.5:0.25 $\mu$M **HPAR3OM**:**HBR35DOM** in confocal microscopy reversibly shifts the initial color of the fluorescence emission at the cell nucleus towards the red one resulting from replacing the initial fluorogen by **HBR35DOM** in the **nirFAST** complex.

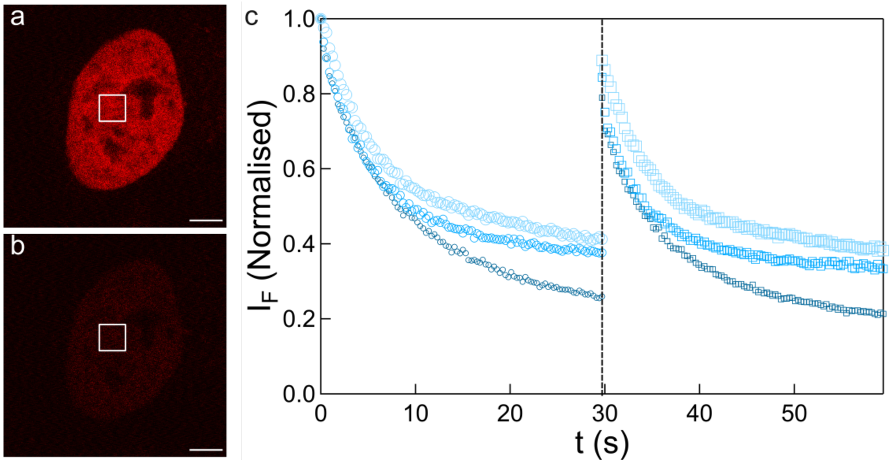

Figure S33: ***FAST** to generate a palette of negative ncRSFPs.* 488 nm light-induced fluorescence emission from **HPAR3Cl** (**a-c**) in **H2B-nirFAST**-labeled living HeLa cells observed in confocal microscopy in the presence of the fluorogen in DMEM pH=7.4. Photoejection is evidenced at 0.1 $\mu$M concentration from the contrast at the nuclei between the initial (**a**) and photosteady state (**b**) images, its reversibility by the similar loss of fluorescence normalized by its initial value after two consecutive series of frame acquisition (circles and squares markers after the 1st and 2nd series respectively) in the region of interest at nuclei, and its concentration-dependence by the distinct decays observed at 0.1 (small black markers), 1 (medium grey markers), or 10 (large light grey markers) $\mu$M fluorogen concentration (**c**). Scale bar: 5 $\mu$m. $T$ = 298 K. See Table S8 for the conditions of image acquisition.

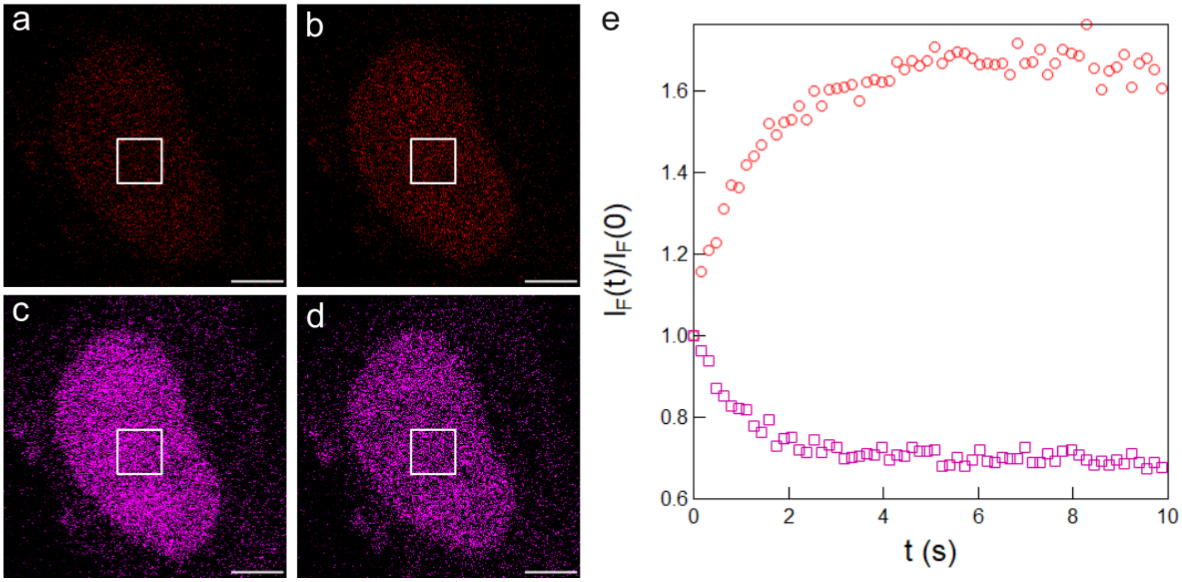

Figure S34: ***FAST** to generate a palette of reversibly photoconvertible fluorescent proteins.* 543 nm light-induced change of fluorescence hue from **HPAR3OM**:**HBR35DOM** in H2B-**nirFAST** labelled living HeLa cells observed in confocal microscopy in the presence of 0.5 $\mu$M **HPAR3OM** and 0.25 $\mu$M **HBR35DOM** in DMEM pH =7.4. Photoconversion of the fluorescence hue is evidenced from the fluorescence contrast at the nuclei between the initial (**a**,**c**) and photosteady state (**b**,**d**) images quantified at $I = 2.5 \times 10^3$ E·m$^{-2}$·s$^{-1}$ light intensity; **e**: Time evolution of the NIR (pink markers for **HPAR3OM**) and red (red markers for **HBR35DOM**) fluorescence normalized by its initial value at $I = 2.5 \times 10^3$ E·m$^{-2}$·s$^{-1}$ light intensity with 1 $\mu$s dwell time in the region interest at nuclei. Scale bar: 5 $\mu$m. $T$ = 298 K. See Table S8 for the conditions of image acquisition.

##### 5.2.2 HBR3F5OM:HPAR3OM

Figure S35 shows that 561 nm illumination of live HeLa cells expressing **H2B-nirFAST** conditioned with 1:1 $\mu$M **HBR3F5OM**:**HPAR3OM** in confocal microscopy reversibly shifts the initial hue of the fluorescence emission at the cell nucleus towards the greener one resulting from replacing the photoisomerized **HPAR3OM** fluorogen by **HBR3F5OM** in the **nirFAST** complex.

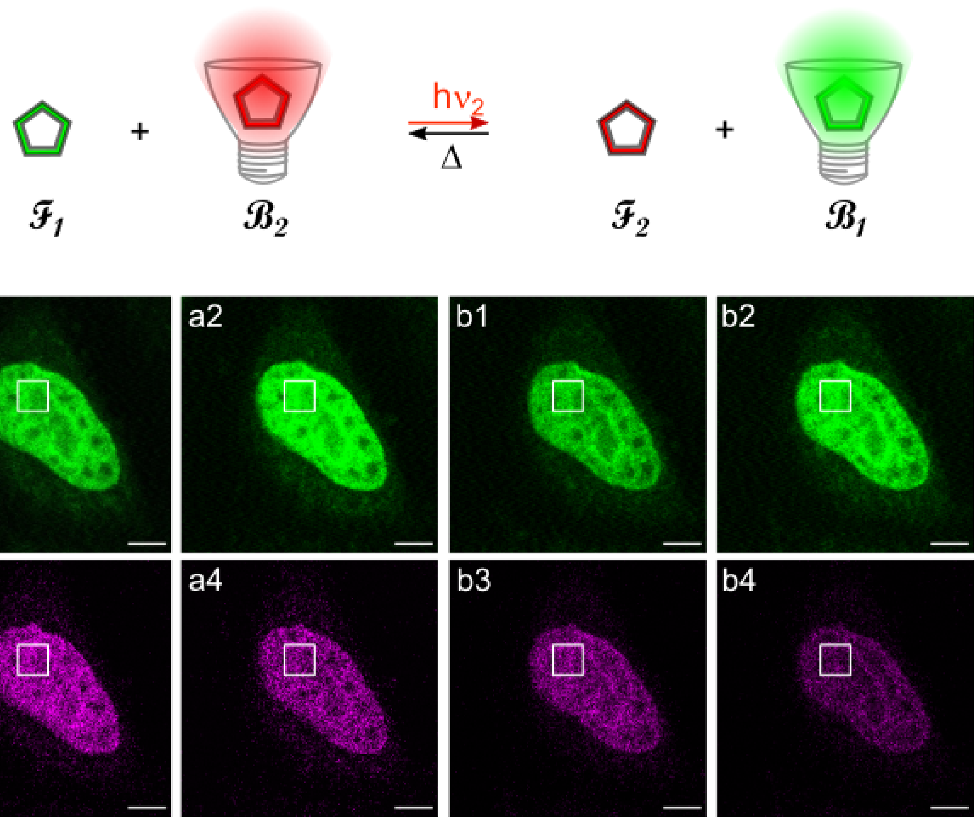

Figure S35: ***FAST*** *to generate a palette of reversibly photoconvertible fluorescent proteins*. 639 nm light ($I = 1.6 \times 10^4$ E·m$^{-2}$·s$^{-1}$)-induced change of fluorescence hue from **HBR3F5OM**:**HPAR3OM** in H2B-**nirFAST** labelled living HeLa cells observed in confocal microscopy in the presence of 1 $\mu$M **HBR3F5OM** and 1 $\mu$M **HPAR3OM** in DMEM pH =7.4. Photoconversion of the fluorescence hue is evidenced from the fluorescence contrast at the nuclei between the initial (**a1**,**a3)** and photosteady state (**a2**,**a4**) images upon exciting at 488 nm at $I = 1.7 \times 10^4$ E·m$^{-2}$·s$^{-1}$ light intensity. Evidence for reversibility is obtained from the similar changes of the fluorescence hue between the initial (**a1/b1** vs **a3/b3**) and photosteady (**a2/b2** vs **a4/b4**) states for two consecutive series of frame acquisition: the **HBR3F5OM** fluorescence signal increased by 10 and 12 % and the **HPAR3OM** one decreased by 24 and 29 % during the first and second series of frame acquisition respectively. Scale bar: 5 $\mu$m. $T = 298$ K. See Table S8 for the conditions of image acquisition.

#### 5.3 Comparative analyses in relation to multiplexed imaging

Refer to Table S7 and Figure S36 for a summary of comparative analyses performed to assess the robustness of the classification pipeline.

- *Six vs. seven vs. eight proteins:* The pipeline was run on the dataset containing six proteins augmented with experimental data corresponding to the protein iFAST and/or oFAST and/or YFAST, using identical segmentation, preprocessing, and training steps. When adding either YFAST or iFAST, the confusion matrix indicates that either additional protein mainly introduces confusion with greenFAST, which exhibits similar kinetic features. When adding oFAST, confusion mainly arises with tFAST and TsiFAST. When adding the three extra proteins, the overall accuracy drops from 98% (six proteins) to 75% (nine proteins) with a gradual performance decay as the proteins are added (see table S7);

- *One vs. two fluorescence channels:* The full analysis on the six proteins was repeated using only the red or green fluorescence channel to assess the gain of dual-channel information. We observed a drop of performance by approximately 20% when using only one channel. To reach performance comparable to the two-channel model (98% accuracy), we had to limit the dataset to 4 proteins (frFAST, greenFAST, nirFAST and tFAST) when using only the green channel (accuracy 97%) and to 3 proteins (pFAST, greenFAST and TsiFAST) when using only the red channel (accuracy 100%). See Figure S37 for the 2D representation of the separability of the proteins when using one channel only;

- *Normalization strategies:* The effect of different normalization strategies on classification performance was evaluated. We compared normalization to the first time point (used in the main analysis) to min-max normalization (0-1 scaling). The results indicate that normalizing to the first point yields better classification accuracy. A confusion between TsiFAST and pFAST appears with this normalisation, signalling that the amplitude change (and not only the kinetic constant) is an important component for the classification.

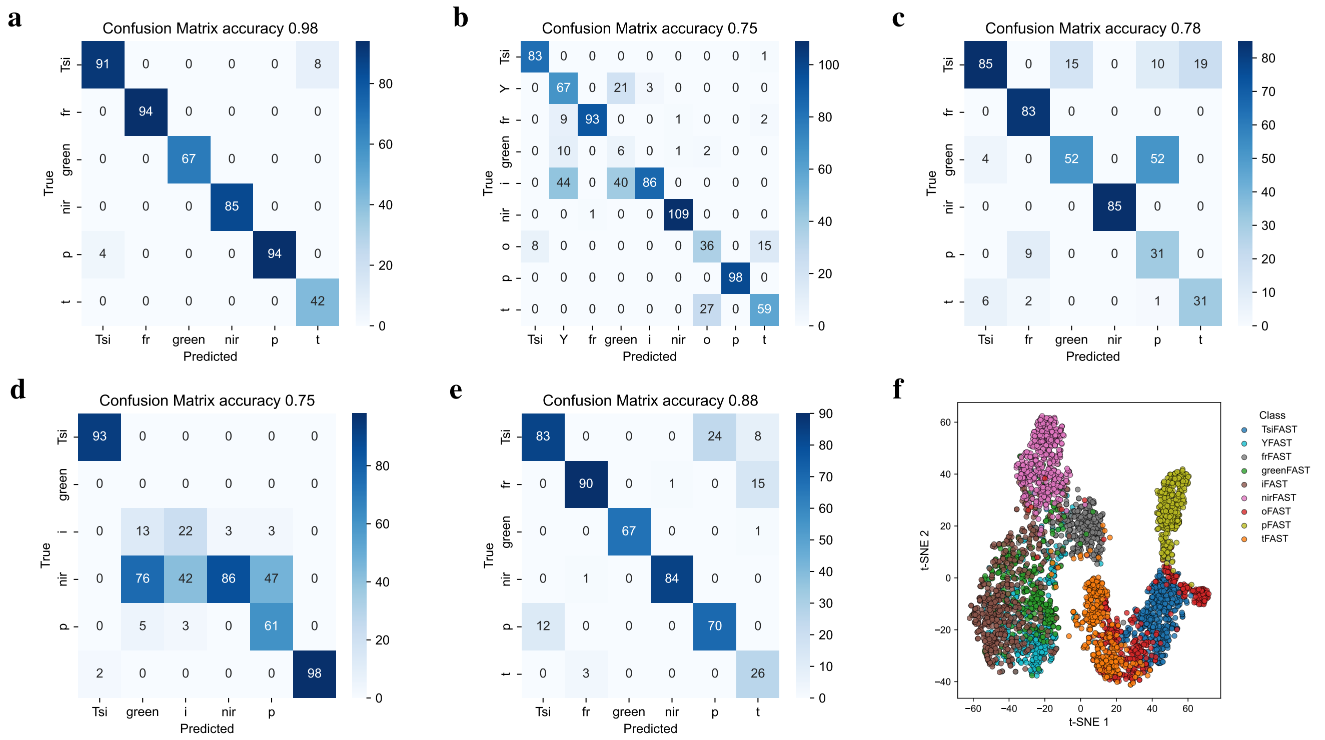

Figure S36: *Study of the parameters affecting the quality of the classification.* **a-e**: Confusion matrices for the classification by Linear Discriminant Analysis model after dimension reduction by PCA to eight components. **f**: A linear supervised reduction of dimension of the red and green traces (Linear Discriminant Analysis, dimension 150 reduced to dimension $n-1$, where $n$ is the number of proteins) is followed by a non-linear dimension reduction technique (t-SNE, dimension $n-1$ reduced to dimension 2) to display the overlap of the signatures of the nine proteins. The different pannels correspond to conditions described in Table S7.

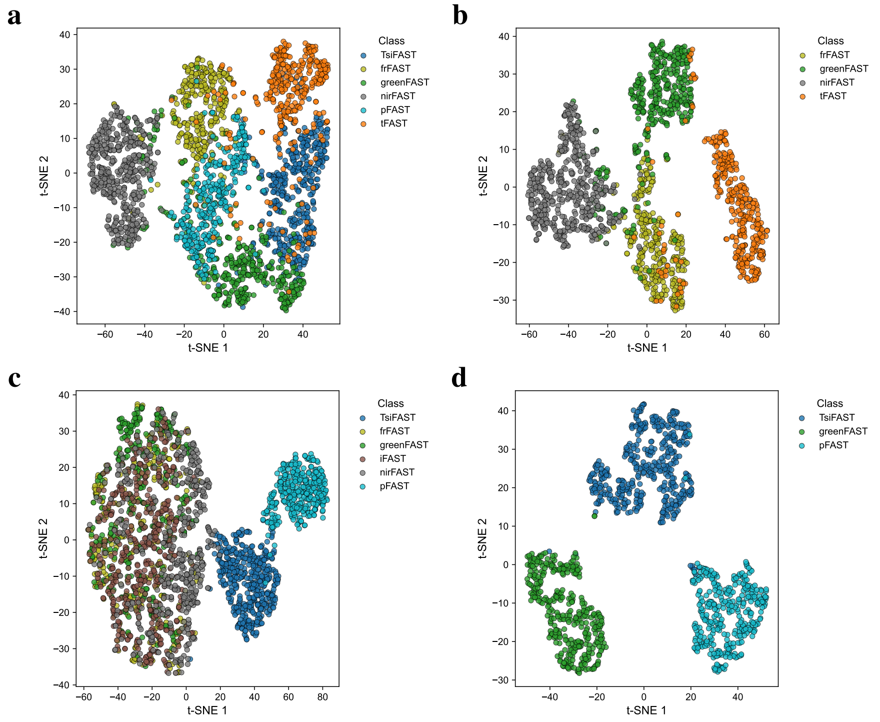

Figure S37: *2D visualisation of the distance between sample points per protein when only one channel is used (green or red)* A linear supervised reduction of dimension (Linear Discriminant Analysis, dimension 75 reduced to dimension $n-1$, where $n$ is the number of proteins) is followed by a non-linear dimension reduction technique (t-SNE, dimension $n-1$ reduced to dimension 2). The reduction is performed using only the green channel (**a** - six proteins, **b** - four proteins) or red channel (**c** - six proteins, **d** - three proteins). The separability of the six proteins is lower when channels are not combined (see Figure 6 of the Main Text for combined channels). The results of the corresponding classification are provided in Table S7.

| proteins | green | red | normalisation | accuracy | Related figure |
|---|---|---|---|---|---|
| 6 | ■ | ■ | first point | 98% | S36a |
| 7 (+ i) | ■ | ■ | first point | 90% | S36f |
| 7 (+ o) | ■ | ■ | first point | 92% | S36f |
| 7 (+ Y) | ■ | ■ | first point | 92% | S36f |
| 8 (no i) | ■ | ■ | first point | 79% | S36f |
| 8 (no o) | ■ | ■ | first point | 83% | S36f |
| 8 (no Y) | ■ | ■ | first point | 83% | S36f |
| 9 | ■ | ■ | first point | 75% | S36b,f |
| 6 | ■ | | first point | 78% | S36c, S37a |
| 4 | ■ | | first point | 97% | S37b |
| 6 | | ■ | first point | 75% | S36d, S37c |
| 3 | | ■ | first point | 100% | S37d |
| 6 | ■ | ■ | 0-1 | 88% | S36e |

Table S7: *Summary of comparative classification analyses.* Each row corresponds to a different condition tested. The last row correspond to confusion matrices in Figure S36.

#### 5.4 Experimental conditions for acquiring the images of the Main Text and the Supporting Information

All the experiments that led to deliver the images reported in the Main Text and in the Supporting Information have been reproduced at least twice.

Table S8: *Experimental conditions for acquiring the images of the Main Text and the Supporting Information* $T$ = 293 K.

| Figure | Fluorogen 1 ($C_1^{\dagger}$) | Fluorogen 2 ($C_2^{\dagger}$) | $I_1^{\pm}$ ($E{\cdot}m^{-2}{\cdot}s^{-1}$) | $I_2^{\mp}$ ($E{\cdot}m^{-2}{\cdot}s^{-1}$) | $\delta^*$ ($\mu$s) | $T^{\ddagger}$ (ms) | $Z^{\times}$ |
|---|---|---|---|---|---|---|---|
| Fig. 3a-c | **HBT35DM** (0.1, 1, 10) | – | $9.1 \times 10^3$ | – | 0.49 | 300 | 4 |
| Fig. 3d-f | **HBR3M** (0.1, 1, 10) | – | $1.7 \times 10^4$ | – | 0.49 | 300 | 4 |
| Fig. 3g-i | **HPAR3OM** (0.1, 1, 10) | – | $3.0 \times 10^4$ | – | 0.49 | 300 | 4 |
| Fig. 3j-l | **HPAR35DOM** (0.1, 1, 10) | – | $3.0 \times 10^4$ | – | 0.49 | 300 | 4 |
| Fig. 4a1-10,4b,5a1-6,7 | **HBR3Cl** (1) | **HBR35DOM** (0.5) | $4.9 \times 10^3$ | – | 0.98 | 150 | 4 |
| Fig. 4c1-4,4d | **HBR3Cl** (1) | **HBR35DOM** (2) | $7.5 \times 10^{-2}$ | – | – | 100 | – |
| Fig. 5b1-6 | **HBT35DM** (10) | **HBR35DOM** (0.25) | $6.5 \times 10^3$ | $2.6 \times 10^3$ | 0.49 | 300 | 4 |
| Fig. 5c1-6 | **HPAR3F5OM** (0.5) | **HBR35DOM** (0.5) | $4.8 \times 10^3$ | – | 1.00 | 150 | 4 |
| Fig. 6a1-5 | **HBIR3Cl** (1) | **HBR35DOM** (0.25) | $2.1 \times 10^3$ | – | 0.49 | 300 | 4 |
| Fig. 6b1-4 | **HBIR3Cl** (1) | **HPAR35DOM** (0.5) | $1.7 \times 10^4$ | $1.6 \times 10^4$ | 1.00 | 150 | 4 |
| Fig.S32 | **HBR25DM** (0.1, 1, 10) | – | $1.7 \times 10^4$ | – | 0.49 | 300 | 4 |
| Fig.S33 | **HPAR3Cl** (0.1, 1, 10) | – | $1.7 \times 10^4$ | – | 0.49 | 300 | 4 |
| Fig.S34 | **HPAR3OM** (0.5) | **HBR35DOM** (0.25) | $2.5 \times 10^3$ | – | 1.00 | 160 | 5 |
| Fig.S35 | **HBR3F5OM** (1) | **HPAR3OM** (1) | $1.6 \times 10^4$ | – | 1.00 | 150 | 4 |

† $C_i$ is the fluorogen concentration in $\mu$M.
* $\delta$ is the dwell time.
‡ $T$ is the duration of frame acquisition.
± $I_1$ is the irradiance of the first light in the illumination.
∓ $I_2$ is the irradiance of the second light in the illumination.
× $Z$ is the zoom factor.

[10] Ian Coghill, Fivos Pham, Mrinal Mandal, Alienor Lahlou, Yuriy Shpinov, Isabelle Aujard, Marie-Aude Plamont, Jacques Fattaccioli, Thomas Le Saux, and Ludovic Jullien. Spatiotemporal light modulation for optical sectioning, diffusion measurement and parallelized experiments with a dual-dmd microscope. 2026.

[11] Aliénor Lahlou, Hessam Sepasi Tehrani, Ian Coghill, Yuriy Shpinov, Mrinal Mandal, Marie-Aude Plamont, Isabelle Aujard, Yuxi Niu, Ladislav Nedbal, Dusan Lazár, et al. Fluorescence to measure light intensity. *Nature Methods*, 20(12):1930–1938, 2023.

[12] Yuriy Shpinov, Ian Coghill, Mrinal Mandal, Philippe Pelupessy, Isabelle Aujard, Thomas Le Saux, and Ludovic Jullien. A modular low-cost 3d uniformly illuminated mini-photoreactor with versatile in-situ analysis. *Advanced Optical Materials*, page 2403466, 2025.
